# Challenges and opportunities for strain verification by whole-genome sequencing

**DOI:** 10.1101/515338

**Authors:** Jenna E Gallegos, Sergei Hayrynen, Neil Adames, Jean Peccoud

## Abstract

Laboratory strains, cell lines, and other genetic materials change hands frequently in the life sciences. Despite evidence that such materials are subject to mix-ups, contamination, and accumulation of secondary mutations, verification of strains and samples is not an established part of many experimental workflows. With the plummeting cost of next generation technologies, it is conceivable that whole genome sequencing (WGS) could be applied to routine strain and sample verification in the future. To demonstrate the need for strain validation by WGS, we sequenced haploid yeast segregants derived from a popular commercial mutant collection and identified several unexpected mutations. We determined that available bioinformatics tools may be ill-suited for verification and highlight the importance of finishing reference genomes for commonly-used laboratory strains.

## The need for sample and strain verification

The frequent transfer of genetic materials between life science organizations introduces opportunities for quality control issues. Genetic mutations accumulate naturally over time, and human errors in labeling and sample preparation are unavoidable. Anecdotally, it is not uncommon for researchers to complain of samples exhibiting unexpected behaviors, only to later discover that the genetic material they’re working with is not as expected.

Laboratory strains, cell lines, and mutant collections exhibit considerable nucleotide variation and background mutations even among lines thought to be isogenic ^1–4^. Despite a growing awareness, cell-line contamination and misidentification is a persistent problem, particularly in mammalian cell research ^5–9^. Comparably, much less attention has been paid to the potential for similar issues in non-mammalian samples. Yet even commonly used plasmids have been shown to vary dramatically from their published sequence ^10^. The problem of plasmid verification has been addressed through the development of a web-based application for assembly of Sanger sequencing reads and alignment of the assembled plasmids with a reference^11^. Strain verification by a similar method will be orders of magnitude more challenging.

The methods currently used to verify samples/strains are biased towards a particular goal. For instance, diagnostic techniques such as PCR, targeted sequencing, or restriction enzyme-based methods are often used to identify whether or not a marker gene or known sequence variant is present, or for analysis of variable repeat regions such as in 16s rRNA profiling ^12,13^. These approaches are limited to particular regions of the genome or are insufficiently sensitive for capturing many types of sequence variations ^3,14,15^.

In addition to wasted time and reagents, undetected genetic variation can lead to severe consequences including delays in publishing or patenting and misplaced conclusions that result in product recalls or retractions^16,17^. Given reports that the life sciences are facing a reproducibility crisis ^18^, it is more important than ever for researchers to verify the samples and strains they work with.

## Applying whole genome sequencing to strain and sample verification

As the cost and turnaround time of next generation sequencing continues to decrease, sample and strain verification by whole genome sequencing (WGS) is becoming a more feasible approach ^12^. Many tools have been developed for assembling sequenced genomes and detecting variants by aligning sequencing reads to a reference genome ^19^, but their applicability to sample and strain verification has not been thoroughly explored.

In order to identify the practical obstacles that must be overcome to ultimately implement WGS as a regular part of genetics workflows, we used haploid yeast strains with an unexpected phenotype derived from a mutant collection as a test case.

As part of a series of yeast cell cycle experiments, we crossed two mutant lines from a knockout collection ^20^ to produce *cln3Δ mbp1Δ* double mutants. *CLN3* and *MBP1* are both involved in promoting cell cycle progression from G1 to S phase^21^, and *mbp1* and *cln3* knockouts are each individually known to result in an increased critical cell size at the start of S phase ^22,23^, so each of the single mutant parents and the double mutant progeny were all expected to exhibit an abnormally large G1 cell size. When the *cln3Δ::kanMX* and *mpb1Δ::natMX* mutant lines were crossed, half of the double mutant progeny had a wild-type G1 cell size. When we examined the *cln3* mutant strain, it too had a wild-type-like cell size. Three of the *cln3Δ mbp1Δ* double mutant segregants were sequenced using an Illumina MiSeq sequencer: one segregant, 1691, exhibited the unexpected wild-type-like phenotype, the others, 1693 and 1694 were used for comparison.

We conducted three different sequence analyses (Figure1): A) Reads were assembled both *de novo* and against the S288C reference genome. B) Variant finding tools were used to call single nucleotide polymorphisms (SNPs) that varied between strains 1691 and 1693. C) Copy number variant tools were used to confirm the presence of strain-specific deletions and marker gene insertions (INDELs) and check for additional structural variations. For each stage of the analysis, at least two different popular bioinformatic tools were tested. Known and unknown variants were confirmed visually (see supplementary Figures 1 & 2 for examples) by aligning trimmed reads to the S288C reference genome using Integrated Genomics Viewer (IGV) software (http://www.broadinstitute.org/igv).

**Figure 1:**
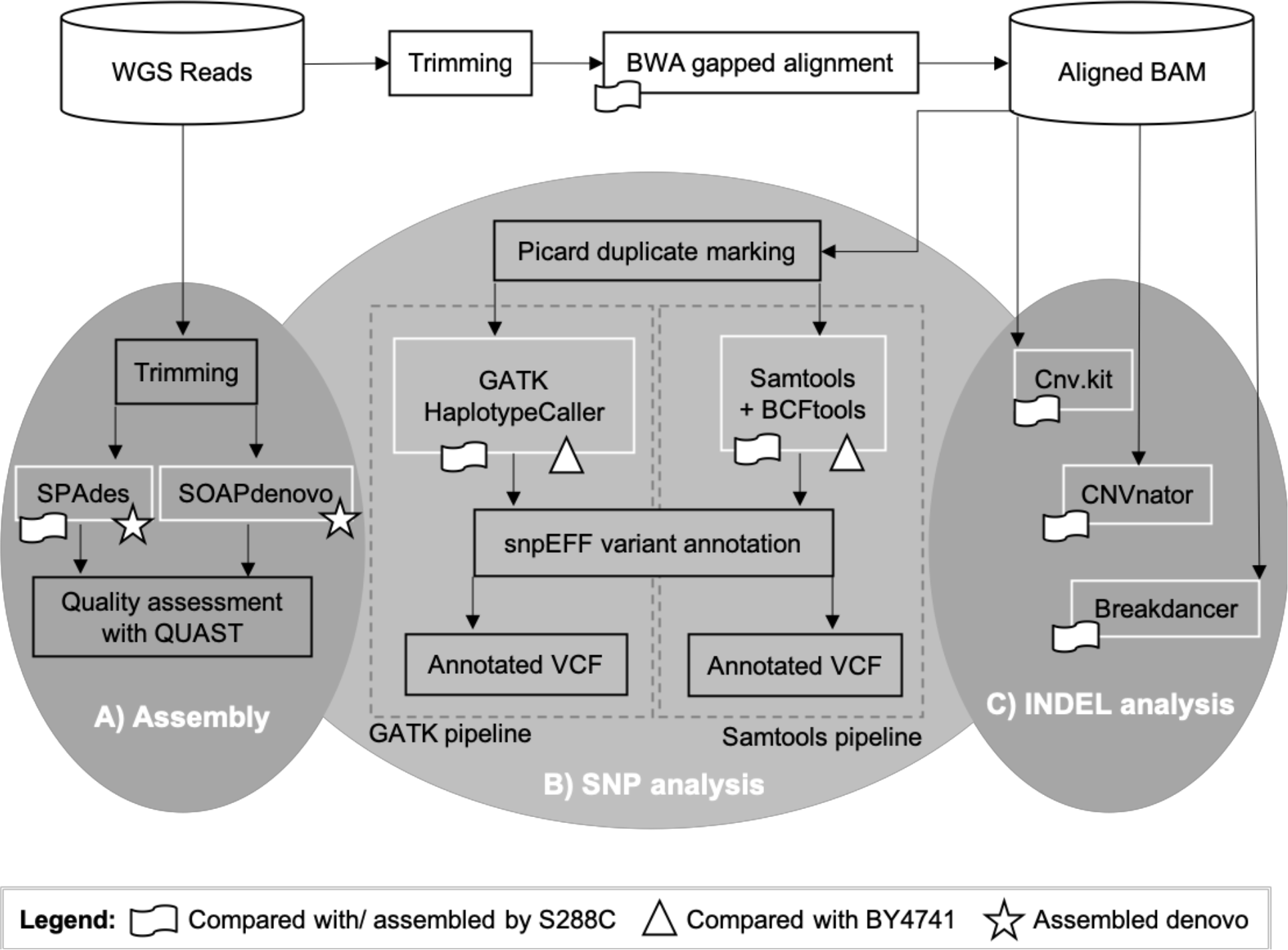
Data analysis pipeline. Software tools compared for each type of analysis are outlined in white. Icons are used to denote whether the analysis was conducted against the S288C reference genome (flag), the BY4741 draft genome (triangle), and or de novo (star). **A)** Reads were assembled using SPAdes^24^ (S288C reference-based and de novo assembly) and SOAPdenovo^25^ (de novo only). **B)** SNPs were analyzed using GATK and Samtools against the S288C reference strain and the BY4741 genetic background. **C)** INDELs and other structural variants were analyzed against the S288C reference using cnv.kit, CNVnator, and Breakdancer. Software versions and parameters used are detailed in Supplementary Information.

### Assembling a genome for verification

An ideal approach to sample verification by WGS would be to sequence the sample, assemble the genome, and then compare the assembled genome to the exact reference genome (the genetic background plus any known variations). Unfortunately, there is not a finished reference genome available for the genetic background used in our analysis (BY4741), despite the fact that it is a commonly used laboratory strain. We thus conducted reference-based assemblies using the closely related S288C genome.

Analyzing a genome assembled *de novo* has a number of additional advantages. For instance, reads that do not align to the reference, such as those for a marker gene insertion or transgene, might be trimmed from the analysis. As such, we repeated the assemblies *de novo*.

The quality of each assembly was compared using the QUAST quality assessment tool for genome assemblies ^26^ (Table 1 and Supplementary Table 2). In all metrics, the reference-based assembly exceeded the *de novo* assemblies, and SPAdes^24^ out-performed SOAPdenovo2^25^.

Although comparable to the previously published BY4741 draft genome^27^, the quality metrics of all assemblies varied markedly from the S288C reference in number of contigs and N50 values.

**Table 1:** Assembly Quality Metrics. Comparison of assembly metrics from different approaches.

| Method | Strain | # contigs | Total length | N50 | # misassembled contigs | Genome fraction (%) | # mismatches per 100 kbp | # indels per 100 kbp |
| --- | --- | --- | --- | --- | --- | --- | --- | --- |
| SPAdes (denovo) | 1691 | 216 | 11700934 | 168266 | 10 | 95.994 | 6.95 | 1.64 |
| SPAdes (denovo) | 1693 | 204 | 11701059 | 188980 | 7 | 96.026 | 7.94 | 1.60 |
| SPAdes (w/ ref) | 1691 | 98 | 11790726 | 356416 | 11 | 96.811 | 8.57 | 1.63 |
| SPAdes (w/ ref) | 1693 | 101 | 11774218 | 332829 | 7 | 96.658 | 10.11 | 1.73 |
| SOAPdenovo2 | 1691 | 283 | 11660175 | 156766 | 8 | 94.251 | 3.34 | 17.67 |
| SOAPdenovo2 | 1693 | 284 | 11642628 | 179827 | 10 | 94.457 | 3.17 | 12.25 |
| Reference | S288C | 17 | 12157105 | 924431 |  |  |  |  |

Cost is a major barrier to using WGS for sample verification, and cost directly relates to coverage. In this study, strains were sequenced on a MiSeq for a total of ∼2 million, 250bp paired-end reads, corresponding to a predicted 80x coverage of the 12 Mb yeast genome. Coverage metrics produced with Picard tools show that ∼95% of genome was covered with at least 30 reads in all three samples (http://broadinstitute.github.io/picard/).

To determine the minimal cost for which a comparable assembly might have been achieved, we repeated the SPAdes assemblies simulating varying levels of coverage by randomly subsampling the read library. For the majority of the quality assembly metrics compared for both reference-based and *de novo* assembly (Figure 2 and Supplementary Tables 3 and 4 respectively), the assembly quality began plateauing at around 500,000 read pairs.

Thus, for a haploid yeast genome, a predicted coverage of 10x-30x is sufficient for a draft genome assembly and investing in NGS coverage beyond 10x may not yield notably better assemblies. To generate a more complete genome, it may thus be more prudent to invest in long read sequencing such as PacBio and Oxford Nanopore technologies in order to conduct hybrid assemblies^28–31^, as opposed to increasing short read depth.

A meaningful direct comparison of our assembled genomes with the reference would require a more complete assembly than we were able to accomplish using short reads alone. As such, we conducted the remainder of our analysis by aligning trimmed reads to the reference genome.

**Figure 2:**
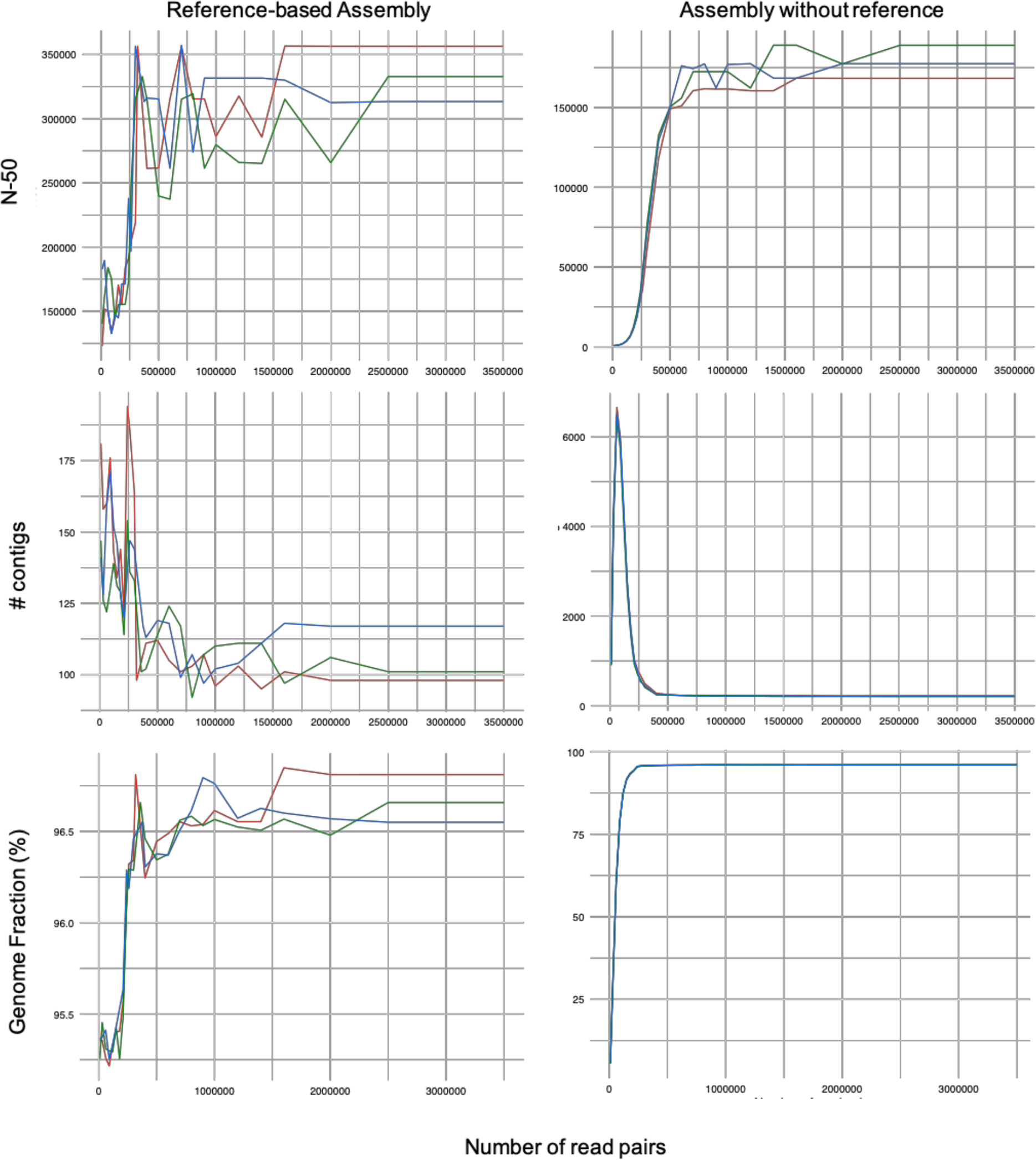
Assembly subsampling analysis. Comparison of the effect of sequencing depth on various metrics (from top to bottom: N50, total number of contigs, and genome fraction) for assemblies against the S288C reference (left) and de novo (right). In each case, the X-axis is number of read pairs. The red, green, and blue lines correspond to reads from 1691, 1693, and 1694 respectively. Additional metrics can be found in Supplementary Tables 3 and 4.

### Variant calling from WGS data

The selection of tools for variant calling can drastically influences the results ^19,32^. We called variants using GATK ^33^ and Samtools ^34^ and attempted to confirm each variant visually by inspecting the reads in IGV (see Supplementary Figure 1 for an example). To simplify the analysis, we focused on only those variants that were discordant between 1691 and 1693.

As has been previously observed ^32^, Samtools identified substantially fewer SNPs than GATK (Supplementary Tables 5 vs 6, respectively). This is likely due to the fact that GATK HaplotypeCaller does local reassembly in regions with genomic variation^35^.

Table 2 highlights all of the SNPs that could be confirmed by aligning the reads to the S288C reference genome in IGV. All of these were identified by both GATK and Samtools. The nine variants called by Samtools that were not also called by GATK (unshaded rows in Supplementary Table 5) could be confirmed for both strain 1691 and 1693 by visual inspection of the aligned reads and thus were not truly discordant. The 58 variants identified by GATK but not Samtools (unshaded and lightly shaded rows in Supplementary Table 6) could all either be confirmed for both strains (not discordant) or for neither (possibly not true SNPs). Only two of the variants that were called by both GATK and Samtools could not be confirmed by inspecting the reads in IGV. The combination of GATK and Samtools, as has previously been proposed ^32^, was thus a valuable approach for filtering out noise in this case.

Analysis with both tools was repeated using the draft genome for the genetic background strain of the parent, BY4741 (Supplementary Tables 7 and 8). This resulted in a substantially longer list of additional discordant SNPs, none of which could be validated by visual inspection of the reads in IGV.

The analysis using Samtools was also repeated using contigs generated by the SPAdes *de novo* assembly (Supplementary Table 9). This also resulted in a longer list of SNPs, but the quality of the calls could not be assessed, because the tool is designed to be used on dozens of reads, not a single contig. This common variant-finding tool is thus not well suited for use with draft genomes or assembled contigs, both of which could facilitate sample and strain verification.

**Table 2:** Consensus variants. Eight variants identified by both GATK and Samtools (using reference S288C). These are all the variants called by either tool that could be confirmed visually by aligning the reads to the reference in IGV (see Supplementary Figure 1 for an example).

| Gene | Function/Notes | Variant | differs from ref | Impact |
| --- | --- | --- | --- | --- |
| *GDC11* (YER025W) | eIF2 subunit (translation) | G > A | 1691 | missense |
| *PPX1* (YHR201C) | hydrolyzes inorganic phosphate | C > T | 1693 | missense |
| *TOR2* (YKL203C) | cell cycle | T > G | 1693 | missense |
| *DYN1* (YKR054C) | cell cycle | G > T | 1691 | missense |
| *SLA2* (YNL243W) | cell cycle | C > T | 1691 | missense |
| *CDC1* (YDR182W) | cell cycle | G > T | 1693 | missense |
| *SCH9* (YHR205W) | protein kinase | C > T | 1693 | synonymous |
| Intergenic | Near Met17 deletion | T > A | 1691 | intergenic |

Of the confirmed SNPs, half occurred in cell cycle related genes (Table 2). None of these are likely to account for the unexpected phenotype observed for 1691. The two variants that were specific to 1691 were located in genes *DYN1* and *SLA2*, both of which are important for cytoskeletal functions ^36,37^. *dyn1* or *sla2* loss-of-function slows the cell cycle and would not relieve the cell cycle delay in G1 caused by the *cln3* mutation^36,37^. However, any of these SNPs could potentially impact the interpretation of cell-cycle experiments.

### INDEL analysis

The segregants sequenced were expected to differ from the S288C reference strain at several auxotrophic marker deletions ^38^, at their mating type genes (the reference strain is *MATα* mating type while 1691 and 1693 are *MAT***a**), and by marker gene insertions at *CLN3* and *MBP1* (shaded cells in Table 3). To confirm the presence or absence of these structural changes in 1691 and 1693 we performed copy-number variant analyses using three different tools: Breakdancer^39^, CNVkit^40^, and CNVnator^41^.

**Table 3:**
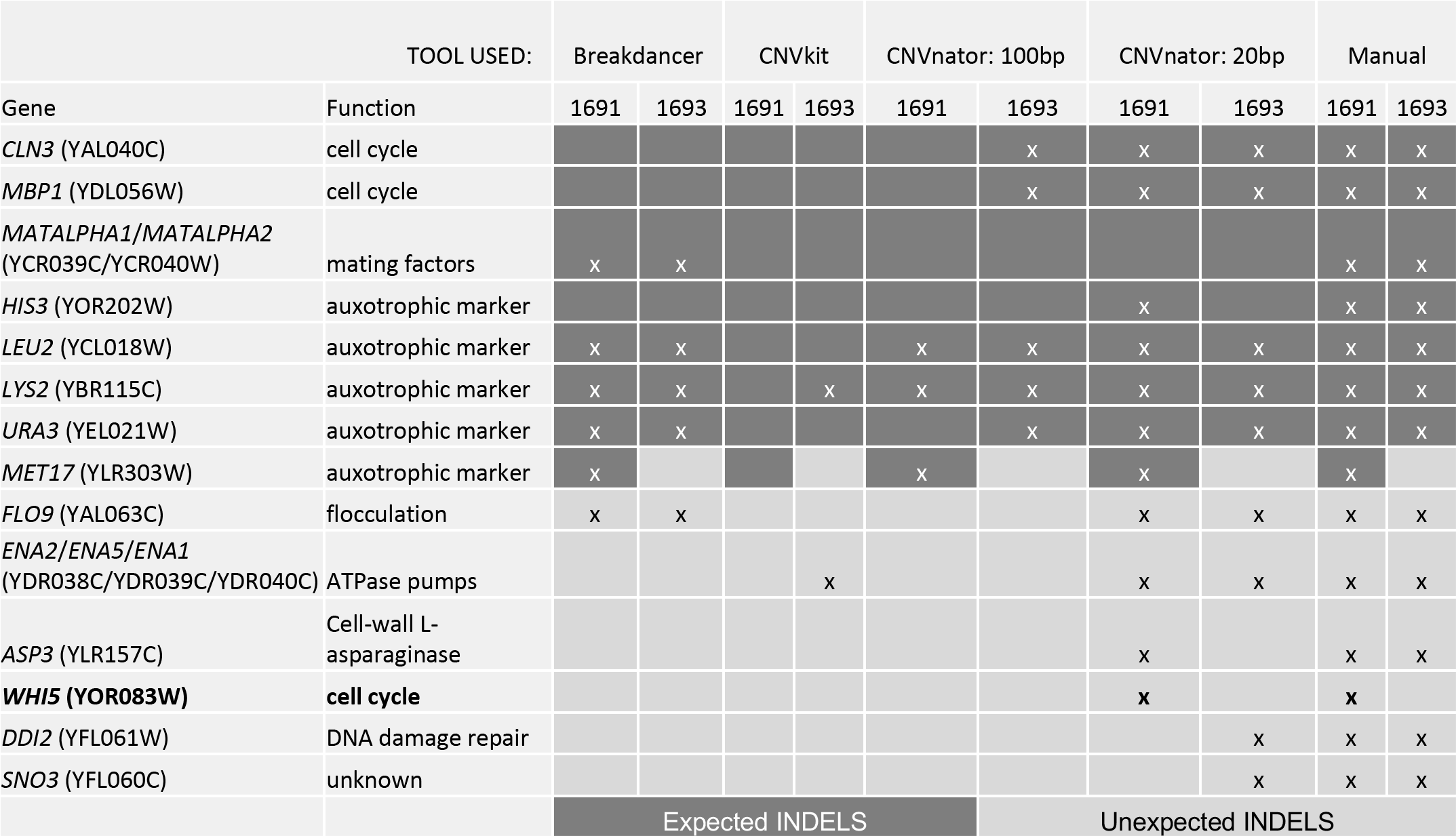
INDEL identification. Identifies which of the INDEL analysis tools succeeded in identifying various expected (dark grey shading) and unexpected (light grey shading) INDELS. For CNVnator, the analysis was conducted twice: once with a bin size of 100bp (recommended for 30x coverage) and once with a bin size of 20bp. Only those INDELS which could be confirmed manually by aligning the reads to the reference genome (see Supplementary Figure 2 for an example) were included. INDELS associated with variable repeat regions such as transposons, telomeres, and ribosomal RNA genes were also excluded. The only unexpected INDEL that differed between 1691 and 1693 occurred in the open reading frame of WHI5 (bolded).

| TOOL USED: |  | Breakdancer |  | CNVkit |  | CNVnator: 100bp |  | CNVnator: 20bp |  | Manual |  |
| --- | --- | --- | --- | --- | --- | --- | --- | --- | --- | --- | --- |
| Gene | Function | 1691 | 1693 | 1691 | 1693 | 1691 | 1693 | 1691 | 1693 | 1691 | 1693 |
| *CLN3* (YAL040C) | cell cycle |  |  |  |  |  | x | x | x | x | x |
| *MBP1* (YDL056W) | cell cycle |  |  |  |  |  | x | x | x | x | x |
| *MATALPHA1/MATALPHA2* (YCR039C/YCR040W) | mating factors | x | x |  |  |  |  |  |  | x | x |
| *HIS3* (YOR202W) | auxotrophic marker |  |  |  |  |  |  | x |  | x | x |
| *LEU2* (YCL018W) | auxotrophic marker | x | x |  |  | x | x | x | x | x | x |
| *LYS2* (YBR115C) | auxotrophic marker | x | x |  | x | x | x | x | x | x | x |
| *URA3* (YEL021W) | auxotrophic marker | x | x |  |  |  | x | x | x | x | x |
| *MET17* (YLR303W) | auxotrophic marker | x |  |  |  | x |  | x |  | x |  |
| *FLO9* (YAL063C) | flocculation | x | x |  |  |  |  | x | x | x | x |
| *ENA2/ENA5/ENA1* (YDR038C/YDR039C/YDR040C) | ATPase pumps |  |  |  | x |  |  | x | x | x | x |
| *ASP3* (YLR157C) | Cell-wall L-asparaginase |  |  |  |  |  |  | x |  | x | x |
| ***WHI5* (YOR083W)** | **cell cycle** |  |  |  |  |  |  | x |  | x |  |
| *DDI2* (YFL061W) | DNA damage repair |  |  |  |  |  |  |  | x | x | x |
| *SNO3* (YFL060C) | unknown |  |  |  |  |  |  |  | x | x | x |
|  |  | Expected INDELS |  |  |  |  | Unexpected INDELS |  |  |  |  |

Table 3 lists all of the INDELs that could be confirmed visually by aligning the reads to the S288C reference genome in IGV (see Supplementary Figure 2 for an example, duplications would be difficult to confirm in this manner). CNVkit found only one of the eight known INDELs in the two segregants but was one of the few tools to identify an event in the ATPase pump genes *ENA1*, *ENA2*, and *ENA5*. Without exploring the parameters to identify more known variants, Breakdancer and CNVnator performed comparably. It was only when we reduced the CNVnator search window down to just 20 bp that we were able to identify most of the known INDELs, even though most of the deletions are hundreds of bp in length (Table 3). These parameters also resulted in by far the longest list of variant calls (Supplementary Table 12 versus Supplementary Tables 10, 11, and 13), most of which occurred in known repetitive regions (lightly shaded rows in Supplementary Table 12).

Only one of the unexpected INDELs, which CNVnator identified as a 1660bp deletion in *WHI5*, differed between 1691 and 1693. *WHI5*, a G1 repressor, controls budding yeast cell size ^42–44^. Deletion of *WHI5* results in smaller cell-size upon entering S-phase, and has been previously shown to partially suppress the large cell phenotype seen in *cln3* mutants ^42^. It is, therefore, very likely that the unexpected wild-type-like phenotype observed for 1691 is due to suppression by a mutation in *WHI5*. A visual inspection of the aligned reads at *WHI5* (Figure 5) suggested that there is a transposon insertion interrupting the *WHI5* reading frame in 1691. The fact that half of the progeny in the cross exhibited the same cell size phenotype as 1691 suggests that the transposon insertion was already present in the *cln3Δ::kanMX* mutant parent obtained from the knock-out collection.

**Figure 3:**
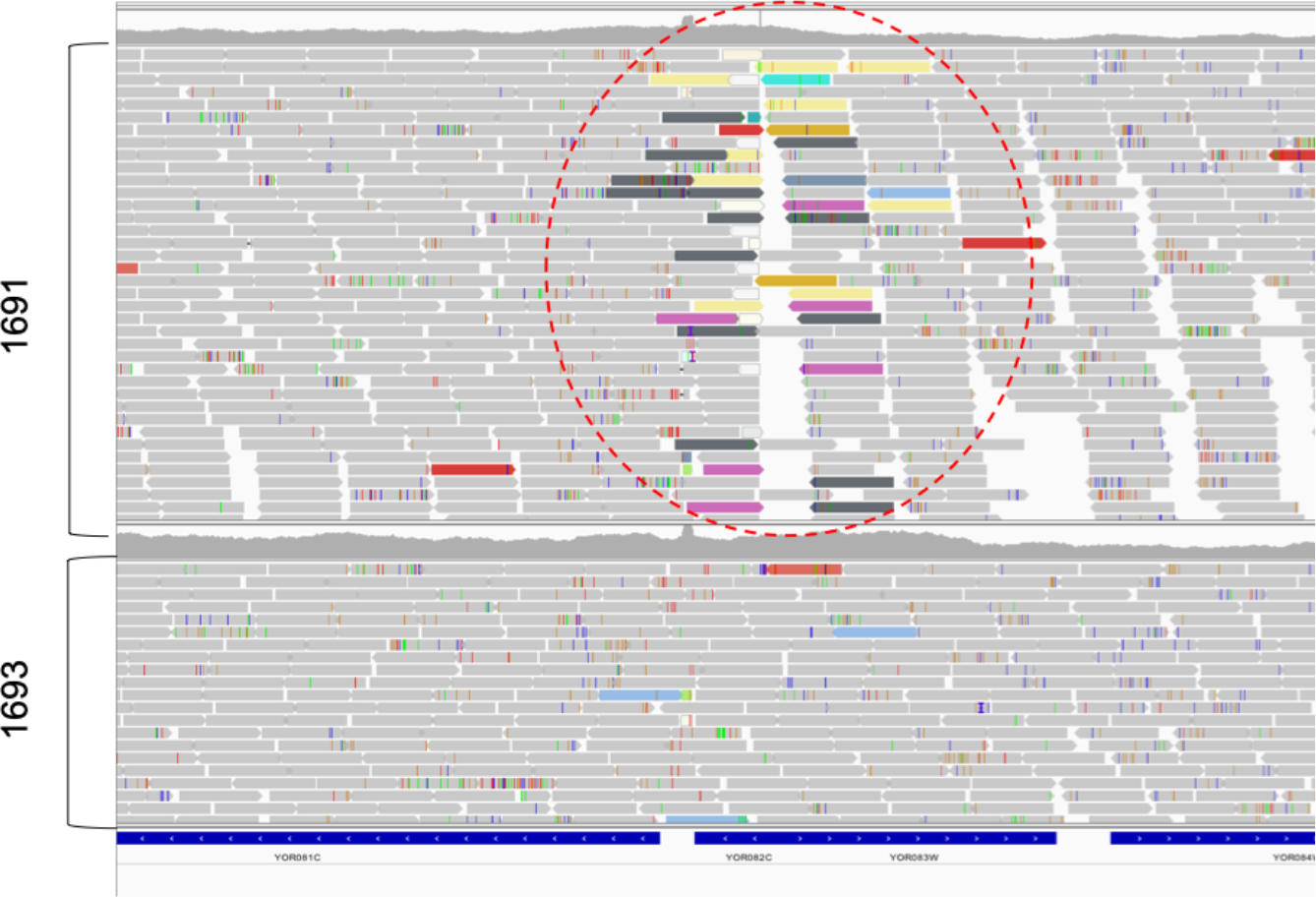
Reads aligned at the WHI5 locus. Each elongated block arrow is a different read. Reads that are colored (not grey) indicate that the mate pair matches a different location in the genome. For each of the colored reads highlighted in the red dashed circle, the mate pair matches a transposon (TY elements).

## Conclusion

Using WGS, we identified eight unexpected SNPs and one unexpected INDEL that differed between segregants derived from a commercial mutant collection. Because this mutant collection is popular in studies of the budding yeast cell cycle, it is pertinent that four of the SNPs identified occurred in cell cycle related genes. This test case demonstrates the value of verifying strains and cell lines from mutant collections by WGS. While this approach was successful in identifying the mutation that was likely the cause for an unexpected phenotype, there may be more changes to the genome that were missed. Clearly, unexpected mutations in common laboratory cell lines cannot be ignored, but the technology needed to get a clear vision of the magnitude of the problem is underdeveloped.

The variant-finding tools used in this analysis were not ideally suited to verification workflows. Most data analysis pipelines, including those described here, rely on ad-hoc or heuristic decision points that require an advanced understanding of the software tools used for analysis ^19^. Analyzing the results required manually validating the calls by visualizing the reads, as well as looking up the function of each individual gene – processes that are tedious, time consuming, and potentially error-prone. Additionally, the SNPs/INDELs called differed dramatically depending on the tools and parameters used. None of the tools and parameters tested successfully identified all of the known INDELs (Table 3). It was only when we adjusted the parameters to find the known INDELS, that we identified a large transposon insertion in an important gene. In conclusion, the software tools used, selected for their popularity in the bioinformatics community, could not reliably return expected outcomes, were individually too narrow in focus, and collectively too sensitive to parameters to be integrated into a consistent pipeline for verification by WGS.

Before WGS can be used for routine sample and strain validation, genome finishing also needs to be streamlined and made more affordable, such that reference genomes are available for all commonly used laboratory strains. Using short reads only, it was difficult to resolve repetitive regions of the genome. This is evident in the fact that most contigs were flanked by transposons. Read alignment across repetitive regions was also ambiguous (for example, Supplementary Figure 2), complicating the variant analysis; many of variants called were located within transposable elements and telomeres, and near genes encoding tRNAs and ribosomal RNAs (lightly shaded rows in Supplementary Tables 10-13). PacBio sequencing would likely have provided an improved resolution, but it remains prohibitively expensive for verification purposes. And while Oxford Nanopore sequencers are affordable, the reagents and flow cells are costly, and the associated software and algorithms are even less accessible to life science researchers without bioinformatics expertise.

In light of the findings presented in this paper, we would like to suggest a call to action for the development of tools and approaches specifically focused on verification by WGS, in order to ultimately implement WGS as a regular part of genetics workflows, such that all genetic materials are verified by WGS prior to experimentation to improve experimental reproducibility.

## Acknowledgements

The research reported in this publication was supported by NIH Award R01GM078989, NSF Award #1832320 “EAGER: Modeling DNA Manufacturing Processes Using Extensible Attribute Grammars”, NSF Award #1759900 “Collaborative Research: ABI Innovation: Automated Prioritization and Design of Experiments to Validate and Improve Mathematical Models of Molecular Regulatory Systems”, and Colorado State University’s Office of the Vice President for Research Catalyst for Innovative Partnerships Program.

## Author Contributions

JG wrote the manuscript text, prepared all figures and tables, and managed the project. SH conducted the bioinformatics analysis, prepared the GitHub documentation, and generated the graphs used in Figure 2. NA carried out the yeast genetic experiments and prepared the samples for WGS. JP devised and oversaw the project.

## Data Availability

All data needed to repeat the analysis described in this manuscript as well as descriptions of the software tools and parameters used is available in the GitHub repository….

