## Supplemental Methods for "Challenges and opportunities for strain verification by whole-genome sequencing"

### Generating and Phenotyping Yeast Mutants

The mutant collection from which the parental strains used in this study were obtained was generated using background strains BY4741 and BY4742, which differ only in their mating type and the auxotrophic markers *MET15* and *LYS2*[1]. Both were derived from *Saccharomyces cerevisiae* strain FY2 which is a direct descendent of S288C. Both BY4741 and BY4742 are known to differ from S288C by the deletion of four auxotrophic markers. According to a recently prepared draft genome, BY4741 additionally differs from S288C by fewer than 5 SNPs per 100,000 bp[2].

One of the haploid parents obtained from the collection has the *CLN3* ORF replaced with a marker for G418 resistance (*MAT***a** *cln3Δ::kanMX MBP1*). The other has the *MBP1* ORF replaced with a marker for nourseothricin resistance by marker switching of the commercial deletion strain (*MAT*[illegible] *CLN3 mbp1Δ::natMX*[3]). These strains were crossed to yield the *MAT***a** haploid progeny we analyzed by whole-genome sequencing.

| **Strain ID** | **Genotype** | **Source** |
|---|---|---|
| 964 (YSC1021-551214) | *MAT***a** *cln3Δ::kanMX his3Δ1 leu2Δ0 met15Δ0 ura3Δ0* | [4] |
| 975 (switched YSC1021-550669) | *MATα mbp1Δ::natMX his3Δ1 leu2Δ0 lys2Δ0 ura3Δ0* | This Study |
| 1691 | *MAT***a** *cln3Δ::kanMX mbp1Δ::natMX his3Δ1 leu2Δ0 met15Δ0 lys2Δ0 ura3Δ0* | This Study |
| 1693 | *MAT***a** *cln3Δ::kanMX mbp1Δ::natMX his3Δ1 leu2Δ0 lys2Δ0 ura3Δ0* | This Study |
| 1694 | *MAT***a** *cln3Δ::kanMX mbp1Δ::natMX his3Δ1 leu2Δ0 ura3Δ0* | This Study |

### Sequencing

Each of the strains was sequenced on a Miseq for a total of ~2 million, 250bp paired-end reads, corresponding to a predicted 80x coverage of the 12Mb yeast genome. Coverage metrics produced with Picard tools show that ~95% of genome was covered with at least 30 reads in all three samples (http://broadinstitute.github.io/picard/). In addition, the percentage of aligned reads versus total reads was in the range of 97-98% for all three samples.

### Analysis

The following software tools were used in the described analysis: FastQC (https://www.bioinformatics.babraham.ac.uk/projects/fastqc/), Trimgalore! (https://www.bioinformatics.babraham.ac.uk/projects/trim_galore/), cutadapt (https://cutadapt.readthedocs.io/en/stable/), bwa (http://bio-bwa.sourceforge.net)[5], Picard (https://broadinstitute.github.io/picard/), ENSEMBL (http://www.ensembl.org

)[6], CNVnator (https://github.com/abyzovlab/CNVnator)[7], Breakdancer (http://breakdancer.sourceforge.net/)[8], cnv.kit (https://cnvkit.readthedocs.io/en/stable/)[9], Samtools (http://www.htslib.org/doc/samtools.html)[10], BCFtools (https://samtools.github.io/bcftools//bcftools.html)[10], Genome Analysis Toolkit (GATK) (https://software.broadinstitute.org/gatk/)[11], snpEFF (http://snpeff.sourceforge.net/)[12], SOAPdenovo2 (http://soap.genomics.org.cn/soapdenovo.html)[13], SPAdes (http://cab.spbu.ru/software/spades/)[14], and QUAST (http://bioinf.spbau.ru/quast)[15], BLAST (http://doi.org/10.1186/1471-2105-10-421). Default parameters were used unless otherwise noted.

FastQC was used to calculate and visualize sequence quality metrics before and after trimming with Trimgalore!. Samples were aligned to Saccharomyces cerevisiae genome assembly R64-1-1 (Ensembl release 92), corresponding to strain S288C (baker's yeast), using BWA (v. 0.7.15) with default parameters. Alignment quality of the resulting bam files was assessed using Picard (v.2.9).

CNVnator (v0.3.3) was used for structural variant calling with a bin size of 20 and 100. In addition, copy number variation was assessed with CNVkit (v0.9.3) and Breakdancer (v. 1.3.6).

CNVkit was run with automatic binning and with p-value threshold of 0.000005 for accepting segments and their breakpoints. Copy numbers were called with log2 ratio thresholds of -1.0000000, 0.5849625, 1.3219281, 1.8073549, and 2.1699250 for copy numbers 0 to 4, with thresholds being the upper limits of log2 coverage ratio for each copy number. Thresholds were calculated by adding 0.5 to the integer copy number value for rounding, dividing by ploidy (1), and log2 transforming the result.

Breakdancer was run with default parameters and configuration files generated using script bam2cfg.pl, included in its distribution.

Variant calling was run both with the GATK pipeline and a pipeline consisting of Samtools (v1.8) and Bcftools (v.1.8).

The GATK pipeline was built on GATK version v4.beta.5, except for function CombineGVCFs, which was run with GATK version v3.8, as there was no working version of this function in GATK 4 at the time of setting up the analyses. GATK was run with default parameters and using GATK HaplotypeCaller for calling variants with *—sample_ploidy* set to 1.

Variant calling with Samtools and Bcftools was run with *—ploidy* set to 1 and using multi-allelic calling mode (Bcftools flag *-m*).

Variants called with GATK and Bcftools were annotated using snpEFF (v.4.3T), a software for variant annotation and predicting effects. Samples aligned to R64-1-1 were annotated using a database for strain S288C provided by snpEFF authors. Samples aligned to BY4741 were annotated using snpEFF database custom built on annotation files downloaded from Saccharomyces Genome Database[16].

*De novo* assemblies were annotated by BLASTing assembled contigs against the reference genome of strain S288C. In addition, variant calling was performed using our *de novo* assembled genome as a reference. Called variants were annotated by BLASTing their flanking sequences against the reference genome of S288C to find corresponding gene annotations. Function *blastn* of BLAST toolbox (v2.7.1+) was run with following parameters: *-outfmt 6 -max_target_seqs 1 -max_hsps 1 -num_threads 10 -strand plus.*

De novo assemblies were performed with SOAPdenovo2 (v. 2.04) and SPAdes (v. 3.9.0), with SPAdes used for downstream analyses and assembly by subsampling. Reference-based de novo assembly, with S288C chromosome sequences used as trusted contigs for i.a. gap closure and repeat resolution, with and without subsampling was also performed with SPAdes (v. 3.9.0).

Qualities of all assemblies were assessed using QUAST, with the reference genome of strain S288C used for benchmarking.
