## Supplementary Data for "Challenges and opportunities for strain verification by whole-genome sequencing"

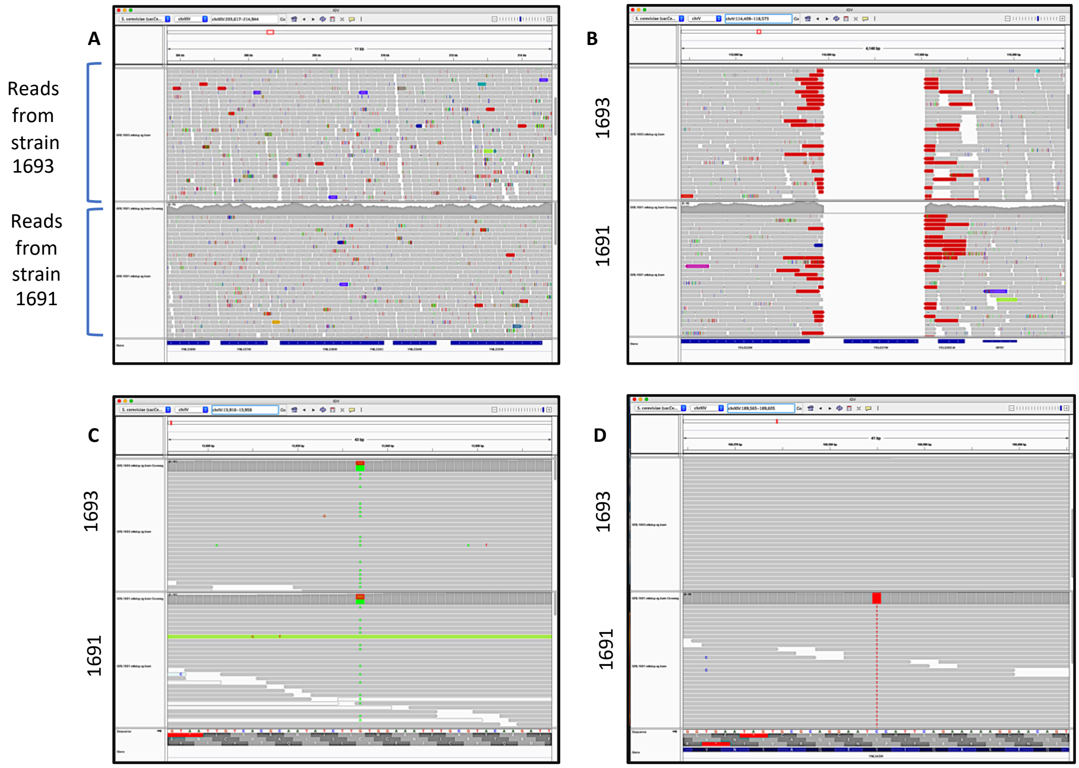


**Supplementary Figure 1. Example IGV screenshots.** A) Normal alignment where the majority of the reads match the reference genome. B) Deletion C) Non-discordant SNP (present in reads from both 1691 and 1693). D) Discordant SNP (present in 1691 only). Grey bars indicate a matched alignment. Colored bars indicate that the mate pair matches another region of the genome. White bars indicate that the mate pair matches multiple regions of the genome. Colored lines indicate a SNP in the read.

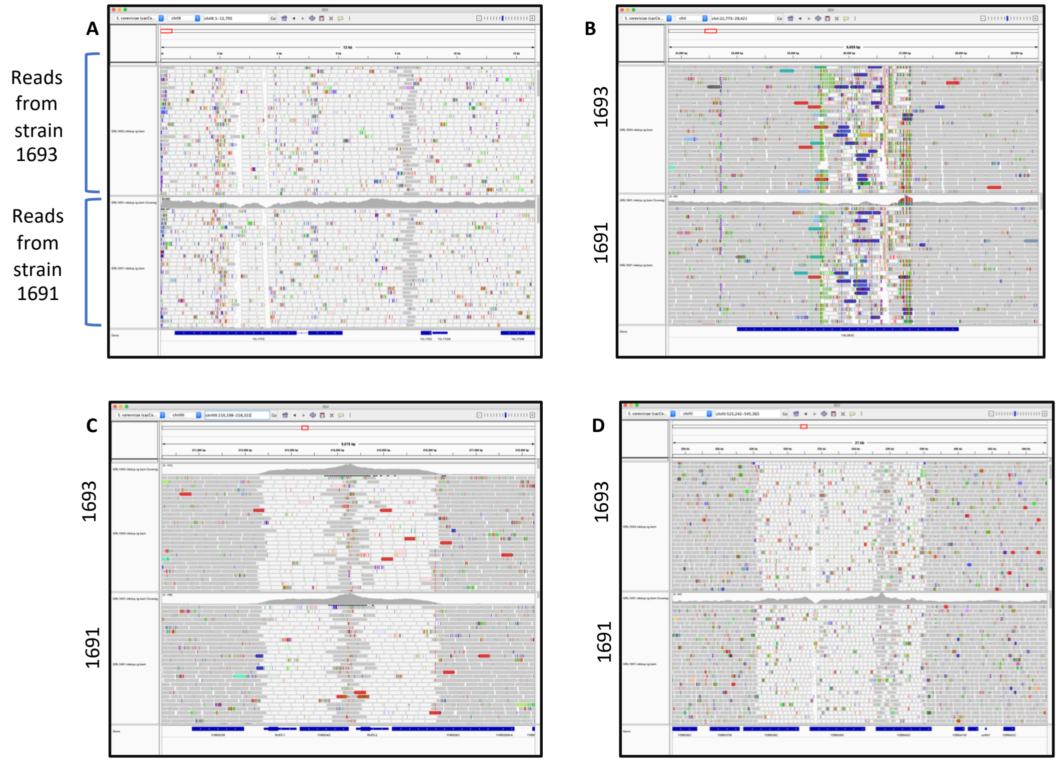


**Supplementary Figure 2. Example IGV screenshots at positions for which a structural variant was called by one or more of the CNV-finding tools.** A) Telomere, B) Transposon, C) ribosomal RNA genes, D) unexpected structural variant. Grey bars indicate a matched alignment. Colored bars indicate that the mate pair matches another region of the genome. White bars indicate that the mate pair matches multiple regions of the genome. Colored lines indicate a SNP in the read.

**Supplementary Table 1: Read Alignment Quality Metrics**

| Library | 1691 | 1693 | 1694 |
|---|---|---|---|
| CATEGORY | FIRST_OF_PAIR | FIRST_OF_PAIR | FIRST_OF_PAIR |
| TOTAL_READS | 1809588 | 2105594 | 2127090 |
| PF_READS | 1809588 | 2105594 | 2127090 |
| PCT_PF_READS | 1 | 1 | 1 |
| PF_NOISE_READS | 0 | 0 | 0 |
| PF_READS_ALIGNED | 1789810 | 2062497 | 2096632 |
| PCT_PF_READS_ALIGNED | 0.98907 | 0.979532 | 0.985681 |
| PF_ALIGNED_BASES | 440082553 | 507794906 | 515044878 |
| PF_HQ_ALIGNED_READS | 1613811 | 1891572 | 1890738 |
| PF_HQ_ALIGNED_BASES | 397272706 | 466108280 | 465047908 |
| PF_HQ_ALIGNED_Q20_BASES | 382132654 | 448663996 | 447045117 |
| PF_HQ_MEDIAN_MISMATCHES | 0 | 0 | 0 |
| PF_MISMATCH_RATE | 0.002786 | 0.002807 | 0.003026 |
| PF_HQ_ERROR_RATE | 0.00273 | 0.002754 | 0.002965 |
| PF_INDEL_RATE | 0.000066 | 0.000064 | 0.000066 |
| MEAN_READ_LENGTH | 247.193158 | 247.410225 | 246.869764 |
| READS_ALIGNED_IN_PAIRS | 1769206 | 2037963 | 2065868 |
| PCT_READS_ALIGNED_IN_PAIRS | 0.988488 | 0.985327 | 0.985327 |
| BAD_CYCLES | 0 | 0 | 0 |
| STRAND_BALANCE | 0.500084 | 0.501316 | 0.500933 |
| PCT_CHIMERAS | 0.012503 | 0.008604 | 0.013095 |
| PCT_ADAPTER | 0.00007 | 0.000059 | 0.000051 |

**Supplementary Table 2: Assembly metrics generated by QUAST.**

| Method | SPAdes | | | | | | SOAPdenovo2 | | |
|---|---|---|---|---|---|---|---|---|---|
| | w/out ref | | | w/ ref | | | SOAPdenovo2 | | |
| Sample | 1691 | 1693 | 1694 | 1691 | 1693 | 1694 | 1691 | 1693 | 1694 |
| # contigs (>= 0 bp) | 609 | 573 | 652 | 395 | 369 | 478 | 1714 | 1720 | 1853 |
| # contigs (>= 1000 bp) | 162 | 158 | 157 | 81 | 82 | 88 | 210 | 197 | 194 |
| # contigs (>= 5000 bp) | 109 | 101 | 101 | 65 | 68 | 73 | 114 | 109 | 111 |
| # contigs (>= 10000 bp) | 100 | 93 | 94 | 63 | 67 | 69 | 103 | 96 | 96 |
| # contigs (>= 25000 bp) | 86 | 77 | 80 | 58 | 61 | 60 | 87 | 81 | 79 |
| # contigs (>= 50000 bp) | 67 | 62 | 66 | 49 | 51 | 53 | 69 | 65 | 62 |
| Total length (>= 0 bp) | 11786208 | 11782596 | 11812579 | 11854059 | 11835283 | 11851133 | 11875194 | 11864959 | 11894585 |
| Total length (>= 1000 bp) | 11662495 | 11668214 | 11669841 | 11780107 | 11762442 | 11748430 | 11610547 | 11583184 | 11595004 |
| Total length (>= 5000 bp) | 11556713 | 11555484 | 11552077 | 11748249 | 11731823 | 11711253 | 11422187 | 11413516 | 11431730 |
| Total length (>= 10000 bp) | 11493176 | 11501635 | 11503714 | 11735246 | 11726310 | 11684834 | 11352487 | 11328934 | 11335570 |
| Total length (>= 25000 bp) | 11233635 | 11237257 | 11249578 | 11643857 | 11636863 | 11525169 | 11077229 | 11063858 | 11059529 |
| Total length (>= 50000 bp) | 10534688 | 10681523 | 10719345 | 11346208 | 11273804 | 11277876 | 10417960 | 10462340 | 10431188 |
| # contigs | 216 | 204 | 217 | 98 | 101 | 117 | 283 | 284 | 286 |
| Largest contig | 515082 | 552891 | 552579 | 707891 | 666561 | 666918 | 555842 | 555360 | 557703 |
| Total length | 11700934 | 11701059 | 11712421 | 11790726 | 11774218 | 11767469 | 11660175 | 11642628 | 11658908 |
| Reference length | 12157105 | 12157105 | 12157105 | 12157105 | 12157105 | 12157105 | 12157105 | 12157105 | 12157105 |
| GC (%) | 38.09 | 38.09 | 38.10 | 38.10 | 38.10 | 38.10 | 38.07 | 38.10 | 38.09 |
| Reference GC (%) | 38.15 | 38.15 | 38.15 | 38.15 | 38.15 | 38.15 | 38.15 | 38.15 | 38.15 |
| N50 | 168266 | 188980 | 177479 | 356416 | 332829 | 313412 | 156766 | 179827 | 182909 |
| NG50 | 159840 | 181742 | 176940 | 356416 | 279390 | 261450 | 151193 | 176254 | 177551 |
| N75 | 95888 | 103340 | 102845 | 159749 | 155699 | 151954 | 95467 | 98165 | 103264 |
| NG75 | 88934 | 97820 | 95604 | 152072 | 147383 | 142745 | 89027 | 91701 | 95470 |
| L50 | 21 | 18 | 20 | 12 | 13 | 14 | 23 | 21 | 19 |
| LG50 | 22 | 19 | 21 | 12 | 14 | 15 | 25 | 22 | 21 |
| L75 | 44 | 39 | 42 | 24 | 26 | 28 | 47 | 43 | 41 |
| LG75 | 48 | 43 | 45 | 26 | 28 | 30 | 51 | 47 | 45 |
| # misassemblies | 10 | 7 | 8 | 11 | 7 | 6 | 10 | 12 | 11 |
| # misassembled contigs | 10 | 7 | 8 | 11 | 7 | 6 | 8 | 10 | 7 |
| Misassembled contigs length | 1326701 | 1098654 | 1346051 | 1523523 | 1593156 | 1422580 | 1518239 | 1250942 | 1173710 |
| # local misassemblies | 15 | 15 | 15 | 12 | 9 | 10 | 139 | 154 | 179 |
| # unaligned mis. contigs | 0 | 0 | 0 | 0 | 0 | 0 | 11 | 9 | 15 |
| # unaligned contigs | 7 + 0 part | 6 + 0 part | 15 + 1 part | 7 + 0 part | 6 + 0 part | 15 + 1 part | 7 + 30 part | 4 + 17 part | 5 + 23 part |
| Unaligned length | 15089 | 14240 | 23033 | 15089 | 14240 | 23033 | 85376 | 49219 | 69053 |
| Genome fraction (%) | 95.994 | 96.026 | 96.014 | 96.811 | 96.658 | 96.55 | 94.251 | 94.457 | 94.405 |
| Duplication ratio | 1.001 | 1.001 | 1.001 | 1.001 | 1.001 | 1.001 | 1.01 | 1.01 | 1.01 |
| # N's per 100 kbp | 0.30 | 1.09 | 0.47 | 0.00 | 0.54 | 0.00 | 1477.35 | 1250.29 | 1332.77 |
| # mismatches per 100 kbp | 6.95 | 7.94 | 7.58 | 8.57 | 10.11 | 8.28 | 3.34 | 3.17 | 2.50 |
| # indels per 100 kbp | 1.64 | 1.60 | 1.80 | 1.63 | 1.73 | 1.62 | 17.67 | 12.25 | 11.53 |
| Largest alignment | 515082 | 552760 | 552579 | 707891 | 666459 | 666819 | 554237 | 553332 | 553953 |
| Total aligned length | 11680223 | 11682390 | 11683547 | 11772862 | 11754537 | 11740664 | 11519639 | 11525166 | 11519379 |
| NA50 | 155132 | 181111 | 177479 | 326499 | 279390 | 313412 | 151105 | 171897 | 177215 |
| NGA50 | 151977 | 177322 | 174318 | 279454 | 265699 | 261450 | 147050 | 169460 | 175719 |
| NA75 | 93593 | 102845 | 95727 | 152072 | 150952 | 147386 | 92273 | 93825 | 102686 |
| NGA75 | 88598 | 96035 | 93593 | 134042 | 122851 | 125395 | 87977 | 91150 | 92313 |
| LA50 | 22 | 19 | 20 | 12 | 14 | 14 | 25 | 22 | 21 |
| LGA50 | 23 | 20 | 22 | 13 | 15 | 15 | 26 | 23 | 22 |
| LA75 | 46 | 41 | 44 | 25 | 27 | 29 | 50 | 46 | 43 |
| LGA75 | 50 | 44 | 47 | 27 | 30 | 31 | 54 | 50 | 47 |

**Supplementary Table 3. Quality metrics for subassemblies by reference.** Assemblies were repeated using random susamplings of reads to artificially simulate a decrease in sequencing depth. Assembly was performed using SPAdes against the S288C reference and metrics were calculated by Quast.

| sample | reads | # N's per 100 kbp | # contigs | # contigs (>=1000 bp) | # contigs (>=5000 bp) | # contigs (>=25000 bp) | # contigs (>=50000 bp) | Largest contig | # indels per 100 kbp | # misassembled contigs | # misassemblies | # mismatches per 100 kbp | # unaligned contigs | Genome fraction (%) | L50 | LA50 | LG50 | LGA50 | N50 | NA50 | NG50 | NGA50 | Total length | Total length (>=1000 bp) | Total length (>=10000 bp) | Total length (>=50000 bp) |
|---|---|---|---|---|---|---|---|---|---|---|---|---|---|---|---|---|---|---|---|---|---|---|---|---|---|---|
| | 10000 | 0 | 181 | 172 | 143 | 109 | 77 | 414717 | 2.46 | 14 | 18 | 34.76 | 5 + 1 part | 95.355 | 29 | 33 | 31 | 35 | 123010 | 114237 | 118247 | 112469 | 11610511 | 11604282 | 11441842 | 9944968 |
| | 30000 | 0 | 158 | 153 | 128 | 91 | 71 | 513280 | 3.42 | 16 | 18 | 71.42 | 5 + 2 part | 95.355 | 24 | 27 | 26 | 29 | 151569 | 136305 | 144428 | 133732 | 11613846 | 11610603 | 11437229 | 10367513 |
| | 60000 | 0 | 160 | 154 | 135 | 103 | 77 | 414762 | 4.4 | 18 | 23 | 82.96 | 6 + 1 part | 95.26 | 26 | 30 | 28 | 32 | 150252 | 118242 | 126842 | 115975 | 11614519 | 11610400 | 11480639 | 10167403 |
| | 90000 | 0 | 176 | 171 | 144 | 109 | 78 | 437853 | 3.5 | 19 | 22 | 68.41 | 2 + 1 part | 95.216 | 29 | 32 | 32 | 35 | 132662 | 115974 | 118247 | 110122 | 11599133 | 11595757 | 11446780 | 9911236 |
| | 120000 | 0 | 143 | 140 | 121 | 98 | 73 | 451555 | 4.31 | 21 | 27 | 58.2 | 3 + 2 part | 95.332 | 24 | 29 | 26 | 32 | 144047 | 123673 | 140081 | 117671 | 11616977 | 11615140 | 11497374 | 10300818 |
| | 150000 | 0 | 134 | 133 | 115 | 85 | 66 | 583019 | 3.84 | 23 | 29 | 41.65 | 3 + 3 part | 95.401 | 21 | 26 | 23 | 28 | 170535 | 144844 | 162924 | 135058 | 11626588 | 11625929 | 11515723 | 10449723 |
| | 180000 | 0 | 144 | 142 | 127 | 92 | 67 | 607528 | 3.18 | 18 | 21 | 30.91 | 3 + 2 part | 95.406 | 22 | 26 | 24 | 28 | 155130 | 127199 | 148906 | 124488 | 11626664 | 11625421 | 11494342 | 10207655 |
| | 210000 | 0 | 122 | 121 | 105 | 84 | 65 | 607530 | 3.62 | 20 | 25 | 30.83 | 3 + 1 part | 95.564 | 20 | 25 | 21 | 27 | 183986 | 143991 | 176753 | 134377 | 11637689 | 11636993 | 11540193 | 10640391 |
| | 240000 | 0.74 | 194 | 150 | 110 | 81 | 63 | 666956 | 2.11 | 8 | 9 | 14.53 | 3 + 1 part | 96.08 | 19 | 20 | 20 | 22 | 191172 | 167037 | 190158 | 157711 | 11724151 | 11693532 | 11533040 | 10592716 |
| | 260000 | 0 | 186 | 131 | 107 | 83 | 62 | 666957 | 2.38 | 5 | 5 | 14.72 | 3 + 0 part | 96.322 | 17 | 18 | 18 | 19 | 203669 | 192335 | 193660 | 189658 | 11749437 | 11715417 | 11604558 | 10567983 |
| | 300000 | 0.09 | 164 | 109 | 89 | 69 | 57 | 743400 | 2.09 | 5 | 5 | 11.89 | 3 + 1 part | 96.341 | 14 | 14 | 15 | 15 | 219212 | 219028 | 219028 | 204362 | 11750245 | 11716658 | 11619700 | 10960019 |
| | 318795 | 0 | 98 | 81 | 65 | 58 | 49 | 707891 | 1.63 | 11 | 11 | 8.57 | 7 + 0 part | 96.811 | 12 | 12 | 12 | 13 | 356416 | 326499 | 356416 | 279454 | 11790726 | 11780107 | 11735246 | 11346208 |
| 1691 | 400000 | 2.02 | 111 | 93 | 79 | 62 | 53 | 666974 | 1.9 | 7 | 7 | 8.7 | 3 + 2 part | 96.245 | 14 | 14 | 14 | 15 | 261295 | 261295 | 261295 | 237090 | 11734126 | 11721326 | 11677255 | 11083735 |
| | 500000 | 2.02 | 112 | 95 | 79 | 62 | 53 | 666973 | 1.77 | 6 | 6 | 9.45 | 3 + 1 part | 96.445 | 13 | 14 | 14 | 14 | 261678 | 260651 | 260651 | 260651 | 11750473 | 11739395 | 11660041 | 11146031 |
| | 600000 | 0 | 105 | 86 | 71 | 58 | 50 | 666905 | 1.81 | 8 | 9 | 8.35 | 6 + 1 part | 96.488 | 13 | 13 | 13 | 14 | 315403 | 315403 | 315403 | 266278 | 11757279 | 11744732 | 11689586 | 11199370 |
| | 700000 | 0 | 101 | 79 | 68 | 57 | 51 | 666986 | 1.71 | 9 | 9 | 8.9 | 6 + 1 part | 96.553 | 12 | 13 | 13 | 14 | 356231 | 315416 | 315416 | 261294 | 11761476 | 11748016 | 11711402 | 11340083 |
| | 800000 | 1.89 | 103 | 84 | 70 | 56 | 49 | 666630 | 1.71 | 10 | 11 | 8.17 | 6 + 1 part | 96.53 | 13 | 13 | 13 | 14 | 315416 | 315416 | 315416 | 282532 | 11758687 | 11746399 | 11699357 | 11236933 |
| | 900000 | 0 | 107 | 84 | 67 | 57 | 51 | 552578 | 1.76 | 10 | 11 | 10.71 | 6 + 1 part | 96.538 | 13 | 13 | 13 | 14 | 315416 | 315416 | 315416 | 312474 | 11758759 | 11744404 | 11696314 | 11352308 |
| | 1000000 | 1.89 | 96 | 80 | 67 | 58 | 52 | 666631 | 1.8 | 7 | 7 | 8.64 | 6 + 1 part | 96.614 | 13 | 14 | 14 | 15 | 286028 | 284556 | 284556 | 261293 | 11765863 | 11755565 | 11721630 | 11356816 |
| | 1200000 | 0 | 103 | 83 | 72 | 62 | 53 | 666818 | 1.59 | 10 | 10 | 8.52 | 7 + 1 part | 96.553 | 13 | 13 | 13 | 14 | 317696 | 317696 | 317696 | 315382 | 11758403 | 11745691 | 11707360 | 11261654 |
| | 1400000 | 0 | 95 | 81 | 68 | 61 | 53 | 666817 | 1.84 | 7 | 7 | 8.47 | 7 + 0 part | 96.553 | 13 | 14 | 14 | 14 | 285688 | 279454 | 279454 | 279454 | 11759040 | 11750741 | 11709072 | 11352276 |
| | 1600000 | 0 | 101 | 82 | 65 | 58 | 50 | 707881 | 1.58 | 11 | 11 | 7.49 | 7 + 0 part | 96.848 | 12 | 12 | 13 | 13 | 356620 | 326499 | 312595 | 312595 | 11794980 | 11782838 | 11738856 | 11374380 |
| | 2000000 | 0 | 98 | 81 | 65 | 58 | 49 | 707891 | 1.63 | 11 | 11 | 8.57 | 7 + 0 part | 96.811 | 12 | 12 | 12 | 13 | 356416 | 326499 | 356416 | 279454 | 11790726 | 11780107 | 11735246 | 11346208 |
| | 2500000 | 0 | 98 | 81 | 65 | 58 | 49 | 707891 | 1.63 | 11 | 11 | 8.57 | 7 + 0 part | 96.811 | 12 | 12 | 12 | 13 | 356416 | 326499 | 356416 | 279454 | 11790726 | 11780107 | 11735246 | 11346208 |
| | 3000000 | 0 | 98 | 81 | 65 | 58 | 49 | 707891 | 1.63 | 11 | 11 | 8.57 | 7 + 0 part | 96.811 | 12 | 12 | 12 | 13 | 356416 | 326499 | 356416 | 279454 | 11790726 | 11780107 | 11735246 | 11346208 |
| | 3500000 | 0 | 98 | 81 | 65 | 58 | 49 | 707891 | 1.63 | 11 | 11 | 8.57 | 7 + 0 part | 96.811 | 12 | 12 | 12 | 13 | 356416 | 326499 | 356416 | 279454 | 11790726 | 11780107 | 11735246 | 11346208 |
| | 10000 | 0 | 147 | 138 | 122 | 93 | 71 | 512281 | 3.74 | 20 | 24 | 49.15 | 9 + 2 part | 95.254 | 23 | 26 | 25 | 29 | 140239 | 125508 | 132940 | 117620 | 11598257 | 11592376 | 11487232 | 10335469 |
| | 30000 | 0 | 126 | 122 | 109 | 84 | 69 | 505503 | 4.57 | 19 | 25 | 83.5 | 5 + 2 part | 95.453 | 21 | 24 | 22 | 26 | 161657 | 144664 | 160396 | 134009 | 11623751 | 11621610 | 11544303 | 10677820 |
| | 60000 | 0 | 122 | 121 | 104 | 82 | 63 | 580181 | 4.44 | 19 | 24 | 91.23 | 3 + 1 part | 95.31 | 19 | 21 | 20 | 23 | 184052 | 155037 | 182607 | 150252 | 11614836 | 11614331 | 11510722 | 10610327 |
| | 90000 | 0 | 130 | 129 | 114 | 85 | 68 | 495837 | 4.27 | 19 | 25 | 75.67 | 2 + 1 part | 95.298 | 20 | 24 | 21 | 26 | 175543 | 144325 | 174855 | 135102 | 11606934 | 11606429 | 11494918 | 10625229 |
| | 120000 | 0 | 139 | 136 | 121 | 99 | 72 | 451542 | 3.9 | 27 | 33 | 61.53 | 4 + 2 part | 95.293 | 24 | 30 | 26 | 32 | 144845 | 118339 | 137144 | 117865 | 11621776 | 11619826 | 11516043 | 10365974 |
| | 150000 | 0 | 131 | 125 | 112 | 92 | 70 | 755428 | 3.07 | 23 | 33 | 44.67 | 3 + 1 part | 95.428 | 21 | 26 | 22 | 28 | 155133 | 140238 | 153289 | 125393 | 11630351 | 11626645 | 11532748 | 10599013 |
| | 180000 | 0 | 129 | 126 | 110 | 90 | 71 | 497518 | 3.2 | 22 | 30 | 32.83 | 5 + 1 part | 95.254 | 22 | 28 | 24 | 30 | 155326 | 134717 | 151482 | 125667 | 11601431 | 11599491 | 11489301 | 10677126 |
| | 210000 | 0 | 114 | 111 | 98 | 84 | 70 | 1027956 | 3.18 | 22 | 33 | 26.78 | 4 + 1 part | 95.485 | 19 | 26 | 20 | 28 | 155391 | 141102 | 153293 | 134183 | 11625640 | 11623492 | 11572256 | 10915605 |
| | 240000 | 0 | 154 | 129 | 106 | 86 | 68 | 557523 | 2.05 | 11 | 11 | 17.83 | 4 + 1 part | 96.111 | 19 | 21 | 21 | 22 | 173590 | 155677 | 162143 | 155132 | 11711496 | 11694772 | 11573510 | 10776603 |
| | 260000 | 0 | 136 | 99 | 80 | 70 | 60 | 666692 | 2.17 | 7 | 7 | 14.16 | 6 + 2 part | 96.293 | 15 | 15 | 15 | 16 | 244870 | 244870 | 244870 | 239164 | 11739553 | 11714982 | 11644747 | 11157434 |
| | 300000 | 0 | 133 | 99 | 81 | 63 | 53 | 666634 | 2.07 | 10 | 12 | 12.04 | 5 + 2 part | 96.287 | 13 | 14 | 14 | 14 | 315415 | 263156 | 263156 | 263156 | 11736384 | 11713760 | 11612564 | 11094187 |
| | 360019 | 0.54 | 101 | 82 | 68 | 61 | 51 | 666561 | 1.73 | 7 | 7 | 10.11 | 6 + 0 part | 96.658 | 13 | 14 | 14 | 15 | 332829 | 279390 | 279390 | 265699 | 11774218 | 11762442 | 11726310 | 11273804 |
| 1693 | 400000 | 0 | 102 | 81 | 68 | 56 | 50 | 666284 | 1.91 | 7 | 7 | 11.18 | 6 + 1 part | 96.46 | 14 | 14 | 14 | 15 | 314974 | 314974 | 314974 | 260326 | 11748060 | 11734262 | 11674075 | 11324814 |
| | 500000 | 0 | 114 | 92 | 77 | 67 | 58 | 666475 | 1.68 | 5 | 5 | 7.26 | 6 + 2 part | 96.344 | 15 | 15 | 16 | 16 | 239814 | 239814 | 220452 | 219562 | 11734659 | 11719702 | 11657184 | 11231156 |
| | 600000 | 0 | 124 | 100 | 83 | 70 | 57 | 666706 | 1.55 | 5 | 5 | 7.23 | 6 + 1 part | 96.375 | 15 | 16 | 16 | 17 | 237320 | 212429 | 224082 | 204132 | 11740465 | 11723923 | 11668479 | 11007490 |
| | 700000 | 0 | 117 | 91 | 74 | 66 | 53 | 666863 | 1.78 | 8 | 9 | 7.33 | 6 + 1 part | 96.561 | 13 | 14 | 14 | 15 | 315224 | 261352 | 279621 | 259773 | 11763183 | 11745304 | 11693850 | 11097745 |
| | 800000 | 0 | 92 | 73 | 65 | 57 | 49 | 666893 | 1.68 | 6 | 6 | 7.91 | 5 + 1 part | 96.583 | 13 | 13 | 14 | 14 | 319353 | 319029 | 315224 | 315224 | 11762970 | 11750830 | 11725852 | 11306259 |
| | 900000 | 0 | 107 | 89 | 75 | 69 | 54 | 666872 | 1.76 | 8 | 8 | 7.87 | 6 + 0 part | 96.532 | 14 | 14 | 15 | 15 | 261400 | 261400 | 261106 | 261106 | 11758542 | 11746532 | 11711515 | 11086209 |
| | 1000000 | 0 | 110 | 88 | 72 | 65 | 51 | 561640 | 1.65 | 8 | 8 | 8.32 | 5 + 2 part | 96.565 | 14 | 14 | 14 | 15 | 279792 | 265699 | 279792 | 261289 | 11762018 | 11747897 | 11711014 | 11084325 |
| | 1200000 | 0 | 111 | 92 | 76 | 66 | 51 | 666869 | 1.6 | 8 | 8 | 7.87 | 6 + 0 part | 96.524 | 14 | 14 | 15 | 15 | 265973 | 261289 | 261289 | 260323 | 11757371 | 11745242 | 11704860 | 10993491 |
| | 1400000 | 0 | 111 | 90 | 76 | 65 | 53 | 666878 | 1.69 | 4 | 4 | 8.95 | 7 + 0 part | 96.506 | 14 | 14 | 15 | 15 | 265170 | 265170 | 261291 | 261291 | 11755960 | 11742744 | 11707469 | 11071418 |
| | 1600000 | 0 | 97 | 78 | 67 | 60 | 51 | 666560 | 1.68 | 6 | 6 | 9.31 | 5 + 1 part | 96.567 | 13 | 13 | 14 | 14 | 315226 | 315226 | 279859 | 279859 | 11760723 | 11748592 | 11719603 | 11289421 |
| | 2000000 | 0.54 | 106 | 85 | 72 | 64 | 53 | 666103 | 1.65 | 6 | 6 | 9.43 | 6 + 0 part | 96.479 | 14 | 14 | 15 | 15 | 265853 | 265699 | 261294 | 261294 | 11751313 | 11738299 | 11703195 | 11185031 |
| | 2500000 | 0.54 | 101 | 82 | 68 | 61 | 51 | 666561 | 1.73 | 7 | 7 | 10.11 | 6 + 0 part | 96.658 | 13 | 14 | 14 | 15 | 332829 | 279390 | 279390 | 265699 | 11774218 | 11762442 | 11726310 | 11273804 |
| | 3000000 | 0.54 | 101 | 82 | 68 | 61 | 51 | 666561 | 1.73 | 7 | 7 | 10.11 | 6 + 0 part | 96.658 | 13 | 14 | 14 | 15 | 332829 | 279390 | 279390 | 265699 | 11774218 | 11762442 | 11726310 | 11273804 |
| | 3500000 | 0.54 | 101 | 82 | 68 | 61 | 51 | 666561 | 1.73 | 7 | 7 | 10.11 | 6 + 0 part | 96.658 | 13 | 14 | 14 | 15 | 332829 | 279390 | 279390 | 265699 | 11774218 | 11762442 | 11726310 | 11273804 |
| | 10000 | 0 | 141 | 137 | 115 | 84 | 62 | 513394 | 3.13 | 15 | 18 | 35.69 | 6 + 1 part | 95.363 | 19 | 21 | 21 | 23 | 182768 | 165207 | 175695 | 153244 | 11608068 | 11605194 | 11485217 | 10314614 |
| | 30000 | 0 | 128 | 125 | 108 | 78 | 63 | 546691 | 4.16 | 20 | 28 | 64.27 | 4 + 2 part | 95.373 | 18 | 25 | 19 | 26 | 189613 | 144830 | 179224 | 143853 | 11619943 | 11618328 | 11488610 | 10687582 |
| | 60000 | 0 | 159 | 150 | 131 | 99 | 72 | 539100 | 3.69 | 21 | 29 | 74.21 | 6 + 0 part | 95.412 | 23 | 29 | 24 | 31 | 150251 | 118348 | 143513 | 112962 | 11621495 | 11616252 | 11477314 | 10159584 |
| | 90000 | 0 | 171 | 163 | 137 | 104 | 72 | 426269 | 3.51 | 19 | 24 | 62.85 | 6 + 2 part | 95.254 | 26 | 31 | 28 | 33 | 132750 | 122842 | 131191 | 118249 | 11604752 | 11599979 | 11444534 | 9951664 |
| | 120000 | 0 | 152 | 147 | 130 | 105 | 73 | 420201 | 4.28 | 23 | 28 | 52.95 | 4 + 3 part | 95.332 | 28 | 32 | 29 | 34 | 147571 | 122953 | 143989 | 115971 | 11614741 | 11611682 | 11526847 | 10034493 |
| | 150000 | 0 | 146 | 139 | 121 | 94 | 68 | 821628 | 3.76 | 21 | 30 | 39.81 | 3 + 4 part | 95.428 | 24 | 30 | 25 | 32 | 144845 | 132660 | 143990 | 125514 | 11631944 | 11627301 | 11501839 | 10285061 |
| | 180000 | 0 | 128 | 126 | 109 | 85 | 68 | 611363 | 3.38 | 26 | 36 | 32.98 | 3 + 2 part | 95.532 | 20 | 28 | 22 | 30 | 171310 | 126322 | 162250 | 124574 | 11637162 | 11635928 | 11538120 | 10676180 |
| | 210000 | 0 | 120 | 117 | 104 | 83 | 65 | 700271 | 3.55 | 23 | 34 | 31.64 | 4 + 2 part | 95.631 | 17 | 25 | 19 | 27 | 171416 | 133032 | 162916 | 126471 | 11653996 | 11652042 | 11572790 | 10678069 |
| | 240000 | 0 | 137 | 110 | 92 | 70 | 58 | 666584 | 1.95 | 12 | 14 | 14.42 | 4 + 1 part | 96.289 | 13 | 16 | 14 | 17 | 237981 | 187151 | 223877 | 181745 | 11736636 | 11719086 | 11636059 | 10926445 |
| | 260000 | 0.23 | 147 | 122 | 93 | 75 | 60 | 666623 | 1.83 | 9 | 9 | 12.49 | 5 + 1 part | 96.188 | 16 | 17 | 17 | 19 | 196695 | 185917 | 189660 | 176273 | 11722272 | 11706776 | 11604474 | 10805077 |
| | 300000 | 0 | 144 | 93 | 79 | 65 | 51 | 707289 | 1.89 | 8 | 9 | 11.12 | 5 + 2 part | 96.46 | 12 | 13 | 13 | 13 | 356506 | 261924 | 262045 | 261924 | 11758916 | 11726497 | 11662143 | 11022998 |
| | 375385 | 0 | 117 | 88 | 73 | 60 | 53 | 666918 | 1.62 | 6 | 6 | 8.28 | 15 + 1 part | 96.55 | 14 | 14 | 15 | 15 | 313412 | 313412 | 261450 | 261450 | 11767469 | 11748430 | 11684834 | 11277876 |
| 1694 | 400000 | 0 | 113 | 89 | 73 | 62 | 52 | 783617 | 2.01 | 7 | 7 | 9.31 | 4 + 1 part | 96.305 | 12 | 14 | 13 | 15 | 316097 | 260320 | 260320 | 257070 | 11731285 | 11715507 | 11653698 | 11135012 |
| | 500000 | 0 | 119 | 91 | 77 | 62 | 55 | 547905 | 1.67 | 3 | 4 | 8.76 | 6 + 1 part | 96.377 | 13 | 14 | 14 | 14 | 315370 | 285793 | 286602 | 285793 | 11746208 | 11727129 | 11668394 | 11244418 |
| | 600000 | 0.23 | 118 | 91 | 77 | 62 | 55 | 743001 | 1.61 | 7 | 7 | 9.22 | 8 + 1 part | 96.37 | 13 | 14 | 14 | 15 | 261409 | 260325 | 260325 | 239785 | 11741087 | 11723196 | 11655502 | 11242261 |
| | 700000 | 0.74 | 99 | 81 | 69 | 59 | 54 | 546109 | 1.59 | 11 | 11 | 9.32 | 8 + 1 part | 96.506 | 13 | 14 | 14 | 14 | 357274 | 285878 | 286075 | 285878 | 11757384 | 11745221 | 11708738 | 11396577 |
| | 800000 | 0 | 107 | 89 | 74 | 62 | 55 | 536085 | 1.65 | 11 | 11 | 9.48 | 8 + 1 part | 96.614 | 14 | 15 | 15 | 16 | 274011 | 261464 | 261464 | 260325 | 11770449 | 11759219 | 11706138 | 11301418 |
| | 900000 | 0 | 97 | 75 | 65 | 55 | 52 | 666721 | 1.7 | 10 | 10 | 11.09 | 7 + 1 part | 96.794 | 13 | 14 | 14 | 14 | 331660 | 319115 | 319439 | 319115 | 11791511 | 11776739 | 11738918 | 11487469 |
| | 1000000 | 0 | 102 | 77 | 65 | 56 | 51 | 666704 | 1.51 | 12 | 12 | 10.54 | 8 + 2 part | 96.762 | 13 | 14 | 14 | 14 | 331660 | 319127 | 319936 | 319127 | 11791625 | 11775141 | 11722438 | 11448526 |
| | 1200000 | 0.88 | 104 | 78 | 63 | 56 | 52 | 743057 | 1.78 | 8 | 8 | 9.8 | 11 + 2 part | 96.572 | 12 | 13 | 13 | 14 | 331660 | 279248 | 279387 | 261852 | 11767745 | 11750839 | 11703058 | 11482563 |
| | 1400000 | 0 | 111 | 78 | 66 | 56 | 51 | 666703 | 1.69 | 11 | 11 | 9.6 | 10 + 2 part | 96.626 | 13 | 14 | 14 | 14 | 331660 | 312482 | 312621 | 312482 | 11774099 | 11752425 | 11712872 | 11404187 |
| | 1600000 | 0 | 118 | 87 | 71 | 57 | 52 | 666726 | 1.67 | 8 | 9 | 8.54 | 12 + 1 part | 96.6 | 13 | 14 | 14 | 14 | 330074 | 312505 | 312644 | 312505 | 11772736 | 11752460 | 11690269 | 11341651 |
| | 2000000 | 0 | 117 | 89 | 73 | 59 | 52 | 666888 | 1.52 | 6 | 6 | 8.39 | 14 + 1 part | 96.569 | 14 | 14 | 14 | 14 | 312535 | 312535 | 312535 | 312535 | 11769018 | 11751079 | 11690459 | 11266793 |
| | 2500000 | 0 | 117 | 88 | 73 | 60 | 53 | 666918 | 1.62 | 6 | 6 | 8.28 | 15 + 1 part | 96.55 | 14 | 14 | 15 | 15 | 313412 | 313412 | 261450 | 261450 | 11767469 | 11748430 | 11684834 | 11277876 |
| | 3000000 | 0 | 117 | 88 | 73 | 60 | 53 | 666918 | 1.62 | 6 | 6 | 8.28 | 15 + 1 part | 96.55 | 14 | 14 | 15 | 15 | 313412 | 313412 | 261450 | 261450 | 11767469 | 11748430 | 11684834 | 11277876 |
| | 3500000 | 0 | 117 | 88 | 73 | 60 | 53 | 666918 | 1.62 | 6 | 6 | 8.28 | 15 + 1 part | 96.55 | 14 | 14 | 15 | 15 | 313412 | 313412 | 261450 | 261450 | 11767469 | 11748430 | 11684834 | 11277876 |

**Supplementary Table 4. Quality metrics for subassemblies without reference.** Assemblies were repeated using random susamplings of reads to artificially simulate a decrease in sequencing depth. Assembly was performed *de novo* using SPAdes and metrics were calculated by Quast.

| sample | reads | # N's per 100 kbp | # contigs | # contigs (>=1000 bp) | # contigs (>=5000 bp) | # contigs (>=25000 bp) | # contigs (>=50000 bp) | Largest contig | # indels per 100 kbp | # misassembled contigs | # misassemblies | # mismatches per 100 kbp | # unaligned contigs | Genome fraction (%) | L50 | LA50 | LG50 | LGA50 | N50 | NA50 | NG50 | NGA50 | Total length | Total length (>=1000 bp) | Total length (>=10000 bp) | Total length (>=50000 bp) |
|---|---|---|---|---|---|---|---|---|---|---|---|---|---|---|---|---|---|---|---|---|---|---|---|---|---|---|
| | 10000 | 0 | 918 | 78 | 7 | 0 | 0 | 9262 | 10.08 | 36 | 36 | 223.1 | 9 + 0 part | 5.63 | 320 | 341 | | | 684 | 667 | | - | 698996 | 157430 | 0 | 0 |
| | 30000 | 1.41 | 4320 | 775 | 7 | 1 | 1 | 80320 | 8.06 | 111 | 116 | 212.36 | 22 + 2 part | 28.977 | 1495 | 1542 | | | 802 | 787 | | - | 3556960 | 1138974 | 92488 | 80320 |
| | 60000 | 2.69 | 6661 | 2827 | 21 | 1 | 1 | 85902 | 6.86 | 122 | 123 | 184.72 | 16 + 8 part | 60.645 | 1935 | 1987 | 4441 | 4546 | 1220 | 1194 | 730 | 716 | 7432400 | 4700722 | 121863 | 85902 |
| | 90000 | 4.86 | 5953 | 3696 | 137 | 2 | 1 | 85902 | 6.9 | 125 | 129 | 146.53 | 11 + 4 part | 78.988 | 1460 | 1504 | 2153 | 2217 | 2017 | 1967 | 1581 | 1538 | 9671067 | 8025551 | 227238 | 85902 |
| | 120000 | 5.79 | 4275 | 3245 | 456 | 1 | 1 | 85903 | 6.55 | 99 | 103 | 113.65 | 6 + 4 part | 87.597 | 917 | 948 | 1138 | 1176 | 3469 | 3386 | 3045 | 2953 | 10712036 | 9953916 | 832326 | 85903 |
| | 150000 | 5.81 | 2727 | 2325 | 749 | 11 | 0 | 46238 | 6.71 | 76 | 77 | 89.66 | 4 + 3 part | 91.506 | 538 | 558 | 620 | 642 | 6279 | 6079 | 5742 | 5592 | 11180707 | 10884360 | 2972572 | 0 |
| | 180000 | 4.13 | 1719 | 1567 | 786 | 44 | 4 | 85903 | 5.82 | 48 | 50 | 67.92 | 5 + 1 part | 93.204 | 322 | 335 | 359 | 373 | 10875 | 10507 | 10331 | 9700 | 11381878 | 11272509 | 6218665 | 287420 |
| | 210000 | 3.31 | 1034 | 983 | 622 | 114 | 16 | 109764 | 5.48 | 46 | 47 | 51.15 | 4 + 0 part | 94.131 | 183 | 191 | 201 | 210 | 18706 | 18093 | 17864 | 17163 | 11489193 | 11451132 | 8895043 | 1062124 |
| | 240000 | 11.16 | 782 | 692 | 490 | 162 | 35 | 97506 | 3.14 | 21 | 21 | 30.27 | 3 + 1 part | 95.394 | 130 | 132 | 139 | 141 | 29035 | 28448 | 28092 | 27789 | 11651486 | 11588842 | 10280064 | 2353786 |
| | 260000 | 15.81 | 695 | 572 | 411 | 168 | 51 | 198779 | 2.96 | 24 | 26 | 24.3 | 5 + 2 part | 95.564 | 99 | 103 | 106 | 110 | 36009 | 35517 | 35208 | 34938 | 11685540 | 11604039 | 10641672 | 3826858 |
| | 300000 | 14.25 | 496 | 391 | 275 | 156 | 80 | 253025 | 2.96 | 18 | 19 | 17.69 | 5 + 2 part | 95.716 | 62 | 63 | 66 | 67 | 61594 | 60030 | 60174 | 57258 | 11701277 | 11632360 | 11127668 | 6907131 |
| | 400000 | 2.64 | 280 | 226 | 156 | 111 | 80 | 356308 | 2.33 | 9 | 10 | 12.69 | 4 + 0 part | 95.834 | 33 | 35 | 35 | 37 | 118460 | 112961 | 112961 | 109336 | 11691487 | 11654203 | 11428200 | 9739303 |
| 1691 | 500000 | 3.36 | 244 | 190 | 129 | 95 | 73 | 445152 | 2.14 | 7 | 8 | 11.26 | 4 + 0 part | 95.908 | 25 | 26 | 27 | 28 | 149182 | 133862 | 133862 | 132459 | 11696837 | 11659319 | 11427219 | 10303599 |
| | 600000 | 2 | 231 | 172 | 121 | 89 | 70 | 515060 | 1.97 | 9 | 10 | 8.92 | 5 + 2 part | 95.954 | 24 | 25 | 25 | 27 | 151017 | 145472 | 150578 | 132732 | 11702346 | 11661444 | 11454118 | 10434496 |
| | 700000 | 0.43 | 233 | 169 | 110 | 84 | 69 | 552455 | 1.86 | 10 | 11 | 7.18 | 5 + 2 part | 95.971 | 22 | 23 | 23 | 25 | 160541 | 149182 | 155132 | 145367 | 11703141 | 11658429 | 11450842 | 10611287 |
| | 800000 | 0.56 | 223 | 163 | 112 | 84 | 67 | 550066 | 2.28 | 13 | 13 | 8.76 | 5 + 2 part | 95.991 | 22 | 24 | 23 | 25 | 161749 | 149182 | 160089 | 148690 | 11703822 | 11661597 | 11458825 | 10553927 |
| | 900000 | 0.3 | 221 | 162 | 114 | 85 | 67 | 550064 | 1.97 | 13 | 13 | 8.45 | 5 + 2 part | 95.996 | 22 | 24 | 23 | 26 | 161549 | 150254 | 160089 | 133077 | 11702002 | 11660437 | 11466770 | 10468001 |
| | 1000000 | 0.38 | 219 | 164 | 111 | 83 | 66 | 541247 | 1.93 | 11 | 11 | 8.72 | 5 + 2 part | 96.031 | 22 | 23 | 23 | 25 | 161617 | 150254 | 160089 | 145367 | 11706041 | 11667682 | 11471305 | 10516627 |
| | 1200000 | 0.47 | 213 | 152 | 104 | 83 | 67 | 541247 | 1.83 | 13 | 13 | 8.43 | 6 + 2 part | 95.966 | 21 | 22 | 22 | 24 | 160486 | 149184 | 155242 | 145033 | 11697217 | 11653977 | 11497448 | 10631114 |
| | 1400000 | 0.3 | 217 | 160 | 104 | 83 | 69 | 552579 | 1.68 | 11 | 11 | 7.06 | 7 + 0 part | 95.994 | 21 | 21 | 22 | 23 | 160485 | 155418 | 155418 | 149184 | 11700593 | 11660658 | 11486659 | 10720463 |
| | 1600000 | 0.3 | 222 | 164 | 106 | 84 | 67 | 515081 | 1.7 | 13 | 13 | 5.95 | 7 + 0 part | 96.01 | 21 | 21 | 22 | 23 | 168354 | 167932 | 159840 | 150253 | 11704019 | 11663095 | 11496685 | 10604280 |
| | 2000000 | 0.3 | 216 | 162 | 107 | 86 | 67 | 515082 | 1.67 | 11 | 11 | 6.92 | 7 + 0 part | 95.993 | 21 | 22 | 22 | 23 | 168266 | 155132 | 159840 | 151977 | 11700815 | 11662376 | 11501971 | 10543483 |
| | 2500000 | 0.3 | 216 | 162 | 107 | 86 | 67 | 515082 | 1.67 | 11 | 11 | 6.92 | 7 + 0 part | 95.993 | 21 | 22 | 22 | 23 | 168266 | 155132 | 159840 | 151977 | 11700815 | 11662376 | 11501971 | 10543483 |
| | 3000000 | 0.3 | 216 | 162 | 107 | 86 | 67 | 515082 | 1.67 | 11 | 11 | 6.92 | 7 + 0 part | 95.993 | 21 | 22 | 22 | 23 | 168266 | 155132 | 159840 | 151977 | 11700815 | 11662376 | 11501971 | 10543483 |
| | 3500000 | 0.3 | 216 | 162 | 107 | 86 | 67 | 515082 | 1.67 | 11 | 11 | 6.92 | 7 + 0 part | 95.993 | 21 | 22 | 22 | 23 | 168266 | 155132 | 159840 | 151977 | 11700815 | 11662376 | 11501971 | 10543483 |
| | 10000 | 0 | 901 | 61 | 5 | 0 | 0 | 18676 | 13.67 | 31 | 36 | 233.17 | 12 + 2 part | 5.535 | 304 | 327 | | | 682 | 656 | | - | 691306 | 151242 | 18676 | 0 |
| | 30000 | 3.52 | 4073 | 782 | 7 | 1 | 1 | 85569 | 10.44 | 89 | 94 | 227.94 | 19 + 7 part | 27.643 | 1397 | 1447 | | | 826 | 802 | | - | 3405864 | 1147831 | 85569 | 85569 |
| | 60000 | 9.18 | 6480 | 2845 | 26 | 1 | 1 | 85902 | 8.99 | 119 | 124 | 197.6 | 17 + 10 part | 60.402 | 1869 | 1925 | 4329 | 4447 | 1253 | 1220 | 748 | 728 | 7403942 | 4789360 | 129991 | 85902 |
| | 90000 | 10.08 | 5716 | 3651 | 150 | 1 | 1 | 85902 | 8.41 | 112 | 119 | 159.7 | 8 + 7 part | 78.623 | 1383 | 1428 | 2064 | 2131 | 2121 | 2072 | 1632 | 1583 | 9620193 | 8113156 | 231754 | 85902 |
| | 120000 | 10.26 | 4032 | 3097 | 508 | 3 | 1 | 85902 | 7.35 | 76 | 78 | 116.95 | 11 + 5 part | 87.679 | 863 | 890 | 1065 | 1098 | 3773 | 3682 | 3330 | 3230 | 10717043 | 10031528 | 987327 | 85902 |
| | 150000 | 10.16 | 2544 | 2195 | 770 | 12 | 1 | 85902 | 6.63 | 61 | 64 | 87.97 | 6 + 1 part | 91.741 | 492 | 512 | 565 | 588 | 6745 | 6503 | 6194 | 6026 | 11205553 | 10951083 | 3444361 | 85902 |
| | 180000 | 7.8 | 1544 | 1418 | 752 | 49 | 2 | 85902 | 5.89 | 49 | 51 | 63.81 | 8 + 1 part | 93.398 | 282 | 296 | 314 | 330 | 12158 | 11611 | 11148 | 10706 | 11405758 | 11314492 | 6728208 | 147278 |
| | 210000 | 5.39 | 923 | 870 | 610 | 123 | 14 | 97037 | 4.91 | 41 | 44 | 45.95 | 4 + 2 part | 94.216 | 169 | 177 | 185 | 193 | 21246 | 20594 | 20492 | 19311 | 11494736 | 11456495 | 9091378 | 959679 |
| | 240000 | 4.21 | 687 | 606 | 444 | 150 | 53 | 141563 | 3.78 | 17 | 17 | 32.79 | 4 + 4 part | 95.424 | 103 | 106 | 111 | 114 | 33953 | 33392 | 31864 | 30765 | 11650521 | 11592198 | 10446738 | 3786853 |
| | 260000 | 3.25 | 603 | 503 | 363 | 156 | 70 | 148513 | 3.66 | 17 | 17 | 25.87 | 5 + 3 part | 95.623 | 81 | 82 | 86 | 87 | 46480 | 45952 | 44087 | 43585 | 11682918 | 11612324 | 10714169 | 5354699 |
| | 300000 | 2.06 | 410 | 324 | 240 | 141 | 85 | 265916 | 2.71 | 16 | 18 | 16.65 | 6 + 2 part | 95.764 | 50 | 54 | 53 | 57 | 77276 | 70669 | 71262 | 68090 | 11693098 | 11632240 | 11195540 | 8033220 |
| | 400000 | 6.39 | 249 | 192 | 137 | 101 | 76 | 451903 | 2.29 | 12 | 13 | 10.74 | 5 + 2 part | 95.86 | 27 | 29 | 29 | 31 | 132745 | 126470 | 132502 | 124512 | 11688093 | 11647780 | 11428168 | 10154443 |
| 1693 | 500000 | 0.56 | 236 | 177 | 122 | 93 | 74 | 451902 | 2.17 | 7 | 8 | 9.05 | 5 + 1 part | 95.902 | 26 | 26 | 27 | 28 | 150487 | 148689 | 148689 | 132857 | 11692739 | 11650663 | 11464975 | 10459646 |
| | 600000 | 0.38 | 228 | 171 | 122 | 90 | 68 | 515066 | 1.8 | 7 | 8 | 9.31 | 5 + 2 part | 95.916 | 23 | 24 | 25 | 26 | 155951 | 150488 | 151168 | 133075 | 11692707 | 11653697 | 11472014 | 10281363 |
| | 700000 | 0.73 | 228 | 170 | 113 | 87 | 63 | 552781 | 1.78 | 6 | 6 | 7.08 | 6 + 1 part | 95.911 | 20 | 21 | 22 | 22 | 172450 | 162108 | 155950 | 155880 | 11691500 | 11651075 | 11464927 | 10284973 |
| | 800000 | 0.48 | 227 | 169 | 111 | 85 | 65 | 552781 | 1.94 | 8 | 8 | 7.93 | 6 + 1 part | 95.995 | 20 | 21 | 22 | 23 | 172450 | 162108 | 156743 | 155880 | 11701019 | 11660955 | 11486274 | 10422348 |
| | 900000 | 0.9 | 224 | 170 | 110 | 84 | 67 | 552777 | 2.02 | 7 | 7 | 8.33 | 6 + 0 part | 96.037 | 21 | 21 | 22 | 22 | 172449 | 168866 | 162236 | 162109 | 11706223 | 11668151 | 11479182 | 10566636 |
| | 1000000 | 0.38 | 223 | 168 | 109 | 84 | 66 | 552888 | 1.94 | 9 | 9 | 8.72 | 5 + 2 part | 96.009 | 22 | 22 | 23 | 24 | 172450 | 162109 | 162236 | 150577 | 11703605 | 11665150 | 11473408 | 10530679 |
| | 1200000 | 0.47 | 218 | 167 | 108 | 84 | 66 | 515675 | 1.76 | 6 | 6 | 8.32 | 6 + 0 part | 95.994 | 21 | 22 | 22 | 23 | 162235 | 159993 | 160094 | 156557 | 11699859 | 11663755 | 11488964 | 10526687 |
| | 1400000 | 0.56 | 208 | 159 | 104 | 78 | 63 | 552963 | 1.86 | 7 | 7 | 8.35 | 7 + 0 part | 96.005 | 18 | 19 | 20 | 20 | 189039 | 181192 | 177322 | 177322 | 11700455 | 11665894 | 11498935 | 10613309 |
| | 1600000 | 0.56 | 206 | 155 | 102 | 78 | 63 | 552961 | 1.88 | 9 | 9 | 8.3 | 7 + 0 part | 95.977 | 18 | 19 | 20 | 20 | 189041 | 181192 | 177322 | 177322 | 11696462 | 11660207 | 11504689 | 10657802 |
| | 2000000 | 1.01 | 207 | 161 | 103 | 79 | 63 | 527849 | 1.53 | 8 | 8 | 7.69 | 6 + 0 part | 96.008 | 19 | 19 | 20 | 21 | 177322 | 177322 | 172450 | 162200 | 11699350 | 11666871 | 11486052 | 10646223 |
| | 2500000 | 1.09 | 204 | 158 | 101 | 77 | 62 | 552891 | 1.6 | 7 | 7 | 7.94 | 6 + 0 part | 96.026 | 18 | 19 | 19 | 20 | 188980 | 181111 | 181742 | 177322 | 11701059 | 11668214 | 11501635 | 10681523 |
| | 3000000 | 1.09 | 204 | 158 | 101 | 77 | 62 | 552891 | 1.6 | 7 | 7 | 7.94 | 6 + 0 part | 96.026 | 18 | 19 | 19 | 20 | 188980 | 181111 | 181742 | 177322 | 11701059 | 11668214 | 11501635 | 10681523 |
| | 3500000 | 1.09 | 204 | 158 | 101 | 77 | 62 | 552891 | 1.6 | 7 | 7 | 7.94 | 6 + 0 part | 96.026 | 18 | 19 | 19 | 20 | 188980 | 181111 | 181742 | 177322 | 11701059 | 11668214 | 11501635 | 10681523 |
| | 10000 | 2.77 | 938 | 70 | 8 | 0 | 0 | 10219 | 14.79 | 34 | 38 | 238.96 | 12 + 2 part | 5.783 | 322 | 348 | | | 688 | 669 | | - | 722653 | 159968 | 10219 | 0 |
| | 30000 | 3.2 | 4149 | 740 | 10 | 2 | 0 | 36411 | 8.58 | 96 | 98 | 198.25 | 22 + 3 part | 27.99 | 1426 | 1469 | | | 812 | 799 | | - | 3441137 | 1106328 | 86590 | 0 |
| | 60000 | 6.11 | 6518 | 2820 | 26 | 2 | 1 | 60084 | 8.71 | 151 | 157 | 185.5 | 19 + 11 part | 60.029 | 1888 | 1950 | 4414 | 4546 | 1236 | 1210 | 730 | 711 | 7360398 | 4715571 | 120352 | 60084 |
| | 90000 | 8.41 | 5876 | 3653 | 129 | 2 | 1 | 85902 | 8.5 | 147 | 153 | 147.82 | 17 + 11 part | 78.661 | 1421 | 1483 | 2124 | 2209 | 2041 | 1966 | 1599 | 1546 | 9631930 | 8006988 | 232359 | 85902 |
| | 120000 | 9.62 | 4278 | 3243 | 458 | 3 | 1 | 81458 | 7.72 | 86 | 91 | 108.95 | 12 + 5 part | 87.559 | 915 | 949 | 1136 | 1176 | 3508 | 3399 | 3068 | 2985 | 10709040 | 9952226 | 813377 | 81458 |
| | 150000 | 9.01 | 2730 | 2329 | 742 | 8 | 1 | 85902 | 6.28 | 64 | 66 | 75.15 | 5 + 1 part | 91.779 | 538 | 558 | 618 | 642 | 6185 | 5997 | 5601 | 5451 | 11208576 | 10915711 | 3016675 | 85902 |
| | 180000 | 5.43 | 1675 | 1514 | 801 | 37 | 2 | 85902 | 5.36 | 55 | 59 | 56.53 | 4 + 1 part | 93.454 | 319 | 333 | 354 | 369 | 11212 | 10814 | 10600 | 10087 | 11413062 | 11297204 | 6377493 | 139373 |
| | 210000 | 4.86 | 1011 | 951 | 654 | 106 | 14 | 119505 | 4.58 | 40 | 43 | 41.11 | 6 + 0 part | 94.353 | 193 | 202 | 211 | 220 | 18068 | 17873 | 17418 | 17137 | 11516431 | 11472640 | 9008380 | 915284 |
| | 240000 | 3.6 | 728 | 641 | 468 | 156 | 51 | 152556 | 3.7 | 26 | 26 | 27.94 | 5 + 1 part | 95.558 | 109 | 115 | 117 | 124 | 30670 | 29877 | 29732 | 28678 | 11674357 | 11614189 | 10268126 | 3601895 |
| | 260000 | 1.88 | 549 | 478 | 366 | 163 | 71 | 227697 | 2.88 | 23 | 23 | 21.95 | 6 + 2 part | 95.71 | 78 | 82 | 83 | 87 | 45010 | 41957 | 42138 | 40704 | 11681288 | 11630940 | 10898745 | 5543360 |
| | 300000 | 1.71 | 450 | 368 | 254 | 149 | 89 | 230685 | 2.94 | 13 | 16 | 16.1 | 5 + 3 part | 95.849 | 55 | 55 | 58 | 59 | 71575 | 70010 | 69401 | 66080 | 11705264 | 11648286 | 11170828 | 7951294 |
| | 400000 | 8.01 | 244 | 188 | 134 | 99 | 74 | 539859 | 2.56 | 11 | 12 | 9.86 | 3 + 4 part | 95.864 | 26 | 27 | 28 | 29 | 129232 | 124075 | 124216 | 122331 | 11692019 | 11651986 | 11455092 | 10146619 |
| 1694 | 500000 | 0.73 | 245 | 179 | 126 | 96 | 70 | 540161 | 1.95 | 10 | 11 | 8.41 | 5 + 2 part | 95.955 | 24 | 26 | 25 | 27 | 150160 | 126783 | 133313 | 124577 | 11710045 | 11664802 | 11475888 | 10206556 |
| | 600000 | 0.66 | 232 | 170 | 117 | 87 | 64 | 515278 | 1.73 | 12 | 13 | 8.36 | 7 + 2 part | 95.992 | 21 | 23 | 23 | 24 | 176155 | 155257 | 162238 | 150636 | 11710452 | 11667067 | 11481145 | 10298219 |
| | 700000 | 0.47 | 219 | 166 | 111 | 85 | 65 | 514933 | 1.83 | 10 | 11 | 9.11 | 7 + 2 part | 96.028 | 21 | 23 | 22 | 24 | 174330 | 157680 | 163424 | 151593 | 11709813 | 11672075 | 11497547 | 10435364 |
| | 800000 | 0.38 | 214 | 162 | 109 | 84 | 66 | 514989 | 1.7 | 10 | 11 | 8.56 | 7 + 2 part | 96.015 | 20 | 22 | 22 | 23 | 177323 | 162111 | 174331 | 157680 | 11704740 | 11668408 | 11500057 | 10534700 |
| | 900000 | 0.81 | 220 | 165 | 112 | 86 | 68 | 553149 | 1.67 | 11 | 11 | 7.83 | 8 + 0 part | 96.015 | 21 | 22 | 22 | 24 | 162237 | 155244 | 154501 | 135426 | 11707840 | 11669131 | 11504737 | 10499507 |
| | 1000000 | 0.47 | 216 | 162 | 107 | 84 | 67 | 533223 | 1.65 | 11 | 12 | 8.66 | 7 + 3 part | 96.042 | 21 | 22 | 22 | 23 | 176940 | 162110 | 166790 | 155244 | 11713507 | 11676048 | 11507897 | 10583060 |
| | 1200000 | 0.55 | 220 | 161 | 104 | 80 | 64 | 552580 | 1.66 | 11 | 11 | 7.73 | 11 + 2 part | 96.043 | 20 | 20 | 21 | 22 | 177433 | 177433 | 176940 | 162111 | 11712409 | 11670926 | 11498306 | 10602185 |
| | 1400000 | 0.38 | 223 | 164 | 108 | 83 | 67 | 552580 | 1.62 | 9 | 9 | 7.77 | 11 + 1 part | 96.047 | 21 | 22 | 22 | 23 | 168378 | 162111 | 162238 | 155261 | 11712862 | 11671229 | 11508145 | 10584339 |
| | 1600000 | 0.47 | 217 | 160 | 106 | 83 | 67 | 552579 | 1.84 | 8 | 8 | 7.7 | 12 + 1 part | 96.002 | 21 | 22 | 22 | 23 | 168391 | 162360 | 162487 | 156641 | 11708767 | 11668762 | 11511717 | 10602520 |
| | 2000000 | 0.56 | 214 | 159 | 103 | 81 | 65 | 552579 | 1.64 | 10 | 10 | 6.79 | 14 + 1 part | 96.008 | 20 | 21 | 21 | 22 | 177479 | 168285 | 176940 | 162360 | 11709716 | 11670665 | 11504771 | 10628385 |
| | 2500000 | 0.47 | 217 | 157 | 101 | 80 | 66 | 552579 | 1.8 | 8 | 8 | 7.58 | 15 + 1 part | 96.014 | 20 | 20 | 21 | 22 | 177479 | 177479 | 176940 | 174318 | 11712421 | 11669841 | 11503714 | 10719345 |
| | 3000000 | 0.47 | 217 | 157 | 101 | 80 | 66 | 552579 | 1.8 | 8 | 8 | 7.58 | 15 + 1 part | 96.014 | 20 | 20 | 21 | 22 | 177479 | 177479 | 176940 | 174318 | 11712421 | 11669841 | 11503714 | 10719345 |
| | 3500000 | 0.47 | 217 | 157 | 101 | 80 | 66 | 552579 | 1.8 | 8 | 8 | 7.58 | 15 + 1 part | 96.014 | 20 | 20 | 21 | 22 | 177479 | 177479 | 176940 | 174318 | 11712421 | 11669841 | 11503714 | 10719345 |

**Supplementary Table 5: Samtools variant analysis against the S288C reference genome.** Columns 1 and 2 detail the position of the variant. Columns 3 and 4 note the reference allele (Genotype 0) and the alternative allele (Genotype 1). Columns 5-7 and 8-10 denote the called genotype and the likelihood for each of the two genotypes for strains 1691 and 1693 respectively. Likelihoods were calculated from the reported Phread-scaled Genotype Likelihoods, with the maximum likelihood being 1. Columns 11-13 report the location of the variant and its effect. For the sake of brevity,  those variants which are shared between 1691 and 1693 are not shown. The most likely discordant variants (those which have a likelihood of 1 for one genotype and <0.00 for the other) are shaded. Each of these was also called by GATK (see supplemenary table 6). Clear discordand SNPs that could be confirmed by aligning the reads to IGV are shaded dark grey (these all had a high likelihood).

| | | | | 1691 | | | 1693 | | | | | |
|---|---|---|---|---|---|---|---|---|---|---|---|---|
| Chrom | POS | Genotype 0 | Genotype 1 | Genoype (G) | Likelihood G=0 | Likelihood G=1 | Genotype (G) | Likelihood G=0 | Likelihood G=1 | Annotation | Gene | Gene ID |
| I | 26391 | A | G | 0 | 0.37 | 0.17 | 1 | 0.14 | 0.38 | synonymous_variant | FLO9 | YAL063C |
| I | 26940 | A | G | 0 | 0.05 | 0.03 | 1 | 0.01 | 0.02 | synonymous_variant | FLO9 | YAL063C |
| I | 26974 | A | G | 0 | 0.04 | 0.03 | 1 | 0.01 | 0.04 | missense_variant | FLO9 | YAL063C |
| I | 204783 | C | G | 0 | 1.00 | 0.85 | 1 | 0.35 | 1.00 | missense_variant | FLO1 | YAR050W |
| I | 204796 | A | G | 0 | 1.00 | 0.85 | 1 | 0.26 | 1.00 | missense_variant | FLO1 | YAR050W |
| IV | 827861 | G | T | 0 | 1.00 | 0.00 | 1 | 0.00 | 1.00 | missense_variant | CDC1 | YDR182W |
| IV | 1308011 | T | C | 0 | 0.78 | 0.48 | 1 | 0.10 | 1.00 | missense_variant | HKR1 | YDR420W |
| V | 205872 | G | A | 1 | 0.00 | 1.00 | 0 | 1.00 | 0.00 | missense_variant | GCD11 | YER025W |
| V | 362749 | A | T | 0 | 1.00 | 0.00 | 1 | 0.00 | 1.00 | upstream_gene_variant | PRS2 | YER099C |
| VI | 4717 | C | G | 0 | 1.00 | 0.00 | 1 | 0.01 | 0.17 | upstream_gene_variant | YFL066C | YFL066C |
| VIII | 499958 | C | T | 0 | 1.00 | 0.00 | 1 | 0.00 | 1.00 | missense_variant | PPX1 | YHR201C |
| VIII | 511255 | C | T | 0 | 1.00 | 0.00 | 1 | 0.00 | 1.00 | synonymous_variant | SCH9 | YHR205W |
| VIII | 514829 | CTATATATATATATATATATATATATATATA | CTATATATATATATATATATATATATATATATA | 1 | 0.00 | 0.03 | 0 | 0.11 | 0.00 | upstream_gene_variant | BAT1 | YHR208W |
| XI | 59567 | T | G | 0 | 1.00 | 0.00 | 1 | 0.00 | 1.00 | missense_variant | TOR2 | YKL203C |
| XI | 541138 | G | T | 1 | 0.00 | 1.00 | 0 | 1.00 | 0.00 | missense_variant | DYN1 | YKR054C |
| XII | 734703 | T | A | 1 | 0.00 | 1.00 | 0 | 1.00 | 0.00 | upstream_gene_variant | YLR302C | YLR302C |
| XIV | 189585 | C | T | 1 | 0.00 | 1.00 | 0 | 1.00 | 0.00 | missense_variant | SLA2 | YNL243W |
| XIV | 415070 | T | TTC | 0 | 1.00 | 0.00 | 1 | 0.02 | 0.40 | upstream_gene_variant | YNL115C | YNL115C |
| XVI | 667520 | T | G | 0 | 1.00 | 0.00 | 1 | 0.00 | 1.00 | upstream_gene_variant | ATG11 | YPR049C |

**Supplementary Table 6: GATK variant calling against reference S288C.** Columns 1 and 2 detail the position of the variant. Columns 3 and 4 note the reference allele (Genotype 0) and the alternative alleles (Genotypes 1 and 2). Colums 5-6 and 7-8 report the genotype called and the confidence associated with the call for strain 1691 and 1693 respectivly (confidence is out of 100). Blank rows indicate that a genotype was not called for that strain. Columns 9-11 report the location of the variant and its effect. For the sake of brevity, those variants which are shared between 1691 and 1693 are not shown. Discordant variants with high confidence (99 for both calls) are shaded in light grey. Variants that were also called by Samtools (see supplementary table 5) are shaded in dark grey (each of these was also high confidence).

| | | | | strain 1691 | | strain 1693 | | | | |
|---|---|---|---|---|---|---|---|---|---|---|
| CHROM | POS | Genotype 0 | Genotype 1, 2 | Genotype | Confidence | Genotype | Confidence | Annotation | Gene | Gene ID |
| I | 65780 | A | AACCGATGAATTCGAGCTCGTTTTCGACA CTGGATGGCGGCGTTAGTATCGAATCGAC AGCAGTATAGCGACCAGCATTCACATACG ATTGACGCATGATATTACTTTCTGCGCACT TAACTTCGCATCTGGGCAGATGATGTCGA G | 0 | 45 | 1 | 99 | frameshift_variant&stop_gained | CLN3 | YAL040C |
| I | 206176 | CG | C | 0 | 99 | 1 | 99 | frameshift_variant | FLO1 | YAR050W |
| I | 206179 | A | AC | | | 1 | 99 | frameshift_variant | FLO1 | YAR050W |
| III | 148614 | T | TTGTTGGAATAAAAATCAACTATCATCTAC TAACTAGTATTTA | 1 | 99 | | | stop_gained&conservative_inframe_insertion | SRD1 | YCR018C |
| III | 148617 | T | TTACTAGTATATTATCATATACGGTGTTAG AAGATGACGCAAATGATGAGAAATAG | 1 | 99 | | | frameshift_variant&stop_gained | SRD1 | YCR018C |
| IV | 1525361 | GGCTTCTCTCT | G | 0 | 99 | 1 | 99 | frameshift_variant | YDR544C | YDR544C |
| II | 68282 | G | GTTATTATTATTATTATTATTATTATTATTA | 0 | 99 | 1 | 99 | disruptive_inframe_insertion | CDC27 | YBL084C |
| IV | 827861 | G | T | 0 | 99 | 1 | 99 | missense_variant | CDC1 | YDR182W |
| IV | 1160444 | C | T | | | 1 | 64 | missense_variant | HXT6 | YDR343C |
| IV | 1525313 | T | G | | | 1 | 99 | missense_variant | YDR544C | YDR544C |
| IV | 1525335 | C | G | | | 1 | 99 | missense_variant | YDR544C | YDR544C |
| IV | 1525340 | C | A | | | 1 | 99 | missense_variant | YDR544C | YDR544C |
| IV | 1525345 | C | G | | | 1 | 99 | missense_variant | YDR544C | YDR544C |
| IV | 1525351 | C | G | | | 1 | 99 | missense_variant | YDR544C | YDR544C |
| IV | 1525357 | C | G | | | 1 | 99 | missense_variant | YDR544C | YDR544C |
| IX | 515 | A | ATAG | 1 | 90 | 0 | 99 | conservative_inframe_insertion | YIL177C | YIL177C |
| IX | 521 | A | T | 1 | 90 | 0 | 99 | missense_variant | YIL177C | YIL177C |
| V | 205872 | G | A | 1 | 99 | 0 | 99 | missense_variant | GCD11 | YER025W |
| VIII | 1846 | A | AGTAGTAGCACTAGTCCTGACGTTGGTGC TGGCAGTGGTAGTAGCATTAGTGCTGGAG TTGGTACTTTCAGTGGTAGTCGCACTAGT GTTGGAGTTGGTACTTTCAGTGGTAGTCG CACTAGTCCTGACGTTGATGCTGGCAGTG | | | 1 | 99 | conservative_inframe_insertion | YHL050C | YHL050C |
| VIII | 289314 | T | G | 1 | 60 | 0 | 99 | missense_variant | AHT1 | YHR093W |
| VIII | 499958 | C | T | 0 | 99 | 1 | 99 | missense_variant | PPX1 | YHR201C |
| XI | 59567 | T | G | 0 | 99 | 1 | 99 | missense_variant | TOR2 | YKL203C |
| XI | 541138 | G | T | 1 | 99 | 0 | 99 | missense_variant | DYN1 | YKR054C |
| XIV | 189585 | C | T | 1 | 99 | 0 | 99 | missense_variant | SLA2 | YNL243W |
| XV | 708819 | G | C | 1 | 99 | 0 | 99 | missense_variant | YOR192C-B | YOR192C-B |
| I | 206126 | T | C | | | 1 | 99 | synonymous_variant | FLO1 | YAR050W |
| I | 206129 | A | C | | | 1 | 99 | synonymous_variant | FLO1 | YAR050W |
| I | 206132 | G | T | | | 1 | 99 | synonymous_variant | FLO1 | YAR050W |
| I | 206144 | C | T | | | 1 | 99 | synonymous_variant | FLO1 | YAR050W |
| IV | 1160352 | C | T | | | 1 | 99 | synonymous_variant | HXT6 | YDR343C |
| VIII | 511255 | C | T | 0 | 99 | 1 | 99 | synonymous_variant | SCH9 | YHR205W |
| XII | 944060 | G | A | | | 1 | 99 | synonymous_variant | YLR410W-B | YLR410W-B |
| XII | 944099 | A | C | | | 1 | 84 | synonymous_variant | YLR410W-B | YLR410W-B |
| XV | 708878 | T | C | 1 | 99 | 0 | 99 | synonymous_variant | YOR192C-B | YOR192C-B |
| I | 101281 | C | CA | | | 1 | 99 | upstream_gene_variant | DRS2 | YAL026C |
| I | 223127 | C | CTTTTTTTTTTTTTTTTTTT , CTTTTTTTTTTTTTTTTTT | 2 | 99 | 1 | 99 | upstream_gene_variant | PHO11 | YAR071W |
| II | 9022 | G | GA | 0 | 99 | 1 | 99 | upstream_gene_variant | YBL111C | YBL111C |
| II | 165902 | C | CT | | | 1 | 59 | upstream_gene_variant | PET9 | YBL030C |
| III | 84780 | A | T | | | 1 | 99 | upstream_gene_variant | YCL022C | YCL022C |
| IV | 13937 | G | A | 0 | 52 | 1 | 99 | upstream_gene_variant | SOR2 | YDL246C |
| IV | 126537 | C | CTTTTTTTTTTTT | 1 | 99 | | | upstream_gene_variant | UFD2 | YDL190C |
| IV | 678013 | C | CTT , CTTTT | 1 | 99 | 2 | 64 | upstream_gene_variant | YDR109C | YDR109C |
| IV | 892698 | G | GTTTTTTT , GTTTT | 2 | 99 | 1 | 99 | upstream_gene_variant | AHA1 | YDR214W |
| IX | 163245 | AT | A | 1 | 73 | 0 | 99 | upstream_gene_variant | SEC24 | YIL109C |
| IX | 187822 | GA | G | 1 | 78 | 0 | 99 | upstream_gene_variant | BMT5 | YIL096C |
| IX | 241032 | G | GA | 0 | 99 | 1 | 99 | upstream_gene_variant | YIL067C | YIL067C |
| IX | 339225 | C | CT | 0 | 99 | 1 | 87 | upstream_gene_variant | EST3 | YIL009C-A |
| IX | 439814 | AGTGTGGTGTGTGGG TGTGG | A | 1 | 99 | 0 | 99 | upstream_gene_variant | YIR042C | YIR042C |
| V | 362749 | A | T | 0 | 99 | 1 | 99 | upstream_gene_variant | PRS2 | YER099C |
| VII | 555981 | C | CAAAA | 1 | 99 | 0 | 99 | upstream_gene_variant | TIM21 | YGR033C |
| VIII | 340058 | TTA | T | 1 | 70 | 0 | 99 | upstream_gene_variant | YHR112C | YHR112C |
| VIII | 514829 | C | CTA | 1 | 99 | 0 | 99 | upstream_gene_variant | BAT1 | YHR208W |
| VIII | 549762 | C | CTTTTTTTT , CTTTTTTTTTTTTTTTTTT | 2 | 99 | 1 | 99 | upstream_gene_variant | YHR214C-B | YHR214C-B |
| XI | 666614 | TGTGTGGGTGTGGTG TGGGTGTGGTGTGGG TGTGGGTGTG | T | 0 | 99 | 1 | 99 | downstream_gene_variant | GEX2 | YKR106W |
| XI | 666740 | T | G | 0 | 99 | 1 | 90 | downstream_gene_variant | GEX2 | YKR106W |
| XI | 666742 | G | T | 0 | 99 | 1 | 90 | downstream_gene_variant | GEX2 | YKR106W |
| XII | 48855 | C | CAAAAAAAAAAAAA | 1 | 99 | 0 | 99 | upstream_gene_variant | YBT1 | YLL048C |
| XII | 468217 | A | G | | | 1 | 38 | upstream_gene_variant | RDN25-2 | RDN25-2 |
| XII | 585920 | GA | G | 1 | 99 | 0 | 99 | upstream_gene_variant | UTP13 | YLR222C |
| XII | 704138 | TA | T | 1 | 61 | 0 | 99 | upstream_gene_variant | YSH1 | YLR277C |
| XII | 734703 | T | A | 1 | 99 | 0 | 99 | upstream_gene_variant | YLR302C | YLR302C |
| XIII | 5924 | A | C | | | 1 | 52 | upstream_gene_variant | YML133C | YML133C |
| XIII | 483743 | GA | G | | | 1 | 50 | upstream_gene_variant | YKU80 | YMR106C |
| XIV | 154923 | GA | G | 0 | 99 | 1 | 99 | upstream_gene_variant | ORC5 | YNL261W |
| XIV | 413448 | C | CT | | | 1 | 96 | upstream_gene_variant | YNL115C | YNL115C |
| XIV | 415054 | C | CTT | | | 1 | 99 | upstream_gene_variant | YNL115C | YNL115C |
| XIV | 415071 | C | T | | | 1 | 99 | upstream_gene_variant | YNL115C | YNL115C |
| XIV | 415088 | C | T | | | 1 | 99 | upstream_gene_variant | YNL115C | YNL115C |
| XIV | 415096 | C | T | | | 1 | 99 | upstream_gene_variant | YNL115C | YNL115C |
| XIV | 479459 | AT | A | 1 | 61 | 0 | 47 | upstream_gene_variant | SWS2 | YNL081C |
| XVI | 667520 | T | G | 0 | 99 | 1 | 99 | upstream_gene_variant | ATG11 | YPR049C |

**Supplementary Table 7: Samtools variant analysis against the BY4741 draft genome.** Columns 1 and 2 detail the position of the variant. Columns 3 and 4 note the reference allele (Genotype 0) and the alternative allele (Genotype 1). Columns 5-7 and 8-10 denote the called genotype and the likelihood for each of the two genotypes for strains 1691 and 1693 respectively. Likelihoods were calculated from the reported Phread-scaled Genotype Likelihoods, with the maximum likelihood being 1. Columns 11-13 report the location of the variant and its effect. For the sake of brevity, those variants which are shared between 1691 and 1693 and 62 variants called in intergenic regions are not shown. The most likely discordant variants (those which have a likelihood of 1 for one genotype and <0.00 for the other) are shaded light grey. Those variants which were also identified by GATK (supplemental table 8) are shaded in dark grey (these were also high likelihood).

| | | | | 1691 | | | 1693 | | | | | |
|---|---|---|---|---|---|---|---|---|---|---|---|---|
| Scaffold (Genbank ID) | POS | Genotype 0 | Genotype 1 | Genoype (G) | Likelihood G=0 | Likelihood G=1 | Genotype (G) | Likelihood G=0 | Likelihood G=1 | Annotation | Gene | Gene ID |
| gi_696447945_gb_JRIS01000005.1_ | 22253 | G | T | 0 | 1.00 | 0.00 | 1 | 0.00 | 1.00 | missense_variant | CDC1 | YDR182W |
| gi_696447920_gb_JRIS01000013.1_ | 87004 | C | T | 1 | 0.00 | 1.00 | 0 | 1.00 | 0.00 | missense_variant | SLA2 | YNL243W |
| gi_696447894_gb_JRIS01000021.1_ | 99 | G | A | 0 | 1.00 | 0.91 | 1 | 0.19 | 1.00 | upstream_gene_variant | | UNDEF10 |
| gi_696447894_gb_JRIS01000021.1_ | 6781 | T | C | 1 | 0.00 | 1.00 | 0 | 1.00 | 0.00 | synonymous_variant | | UNDEF12 |
| gi_696447894_gb_JRIS01000021.1_ | 10355 | CTATATATATATATATATATATATATATA | CTATATATATATATATATATATATATATATA | 1 | 0.00 | 0.02 | 0 | 0.11 | 0.00 | upstream_gene_variant | | UNDEF18 |
| gi_696447858_gb_JRIS01000030.1_ | 83846 | T | A | 0 | 1.00 | 0.00 | 1 | 0.00 | 1.00 | downstream_gene_variant | STT4 | YLR305C |
| gi_696447821_gb_JRIS01000042.1_ | 106687 | G | T | 1 | 0.00 | 1.00 | 0 | 1.00 | 0.00 | upstream_gene_variant | ATG11 | YPR049C |
| gi_696447795_gb_JRIS01000048.1_ | 48923 | CTATATATATATATATATATATATATATATA | CTATATATATATATATATATATATATATA | 1 | 0.01 | 0.02 | 0 | 0.17 | 0.00 | upstream_gene_variant | TOM71 | YHR117W |
| gi_696447774_gb_JRIS01000053.1_ | 29830 | A | G | 0 | 1.00 | 0.76 | 1 | 0.17 | 1.00 | upstream_gene_variant | PHO87 | YCR037C |
| gi_696447735_gb_JRIS01000063.1_ | 32496 | TTATTATTNNNNNNNNNNNNNNNNNNNNNNNNNNTAT<br>TATT | TTATTATT | 1 | 0.00 | 0.02 | 0 | 1.00 | 0.00 | frameshift_variant | | UNDEF61 |
| gi_696447735_gb_JRIS01000063.1_ | 32502 | TTNNNNNNNNNNNNNNNNNNNNNNNNNNT | TTATTATTATTATTATTATNNNNNNNNNNNNNNNN<br>NNNNNNNNNT | 1 | 0.00 | 0.10 | 0 | 0.56 | 0.00 | frameshift_variant&stop_gained | | UNDEF61 |
| gi_696447735_gb_JRIS01000063.1_ | 130139 | CTC | CTCTTTTTTTTTTTTTTTTTTTTTTTTTTC | 0 | 0.76 | 0.04 | 1 | 0.01 | 0.09 | upstream_gene_variant | PET9 | YBL030C |
| gi_696447727_gb_JRIS01000065.1_ | 83619 | ATAATAATAATAATAATAATAATAATAATNNNNNNN<br>NNNNNNNNNNNNNNNNNNNNNNNNNNNNNNNNNNN<br>NNNNNTAATAATAATAATAATAATAATAATAAT | ATAATAATAATAATAATAATAATAATAAT | 1 | 0.00 | 0.01 | 0 | 0.00 | 0.00 | upstream_gene_variant | FKH1 | YIL131C |
| gi_696447727_gb_JRIS01000065.1_ | 83622 | ATAATAATAATAATAATAATAATAATNNNNNNNNNN<br>NNNNNNNNNNNNNNNNNNNNNNNNNNNNNNNNNNN<br>NNTAATAATAATAATAATAATAATAAT | ATAATAATAATAATAATAATAATAAT | 1 | 0.00 | 0.01 | 0 | 0.00 | 0.00 | upstream_gene_variant | FKH1 | YIL131C |
| gi_696447727_gb_JRIS01000065.1_ | 83625 | ATAATAATAATAATAATAATAATNNNNNNNNNNNNN<br>NNNNNNNNNNNNNNNNNNNNNNNNNNNNNNNNNNT<br>AATAATAATAATAATAATAAT | ATAATAATAATAATAATAATAAT | 1 | 0.00 | 0.01 | 0 | 0.00 | 0.00 | upstream_gene_variant | FKH1 | YIL131C |
| gi_696447727_gb_JRIS01000065.1_ | 116337 | ACCCCCCCCC | ACCCCCCCCCCCCCC | 0 | 0.06 | 0.04 | 1 | 0.01 | 0.03 | upstream_gene_variant | RPI1 | YIL119C |
| gi_696447695_gb_JRIS01000074.1_ | 1172 | C | T | 0 | 0.02 | 0.01 | 1 | 0.00 | 0.01 | missense_variant&splice_region_variant | YNL019C | YNL019C |
| gi_696447659_gb_JRIS01000084.1_ | 56773 | T | G | 0 | 1.00 | 0.00 | 1 | 0.00 | 1.00 | missense_variant | TOR2 | YKL203C |
| gi_696447656_gb_JRIS01000085.1_ | 20647 | C | T | 0 | 0.32 | 0.20 | 1 | 0.04 | 0.32 | synonymous_variant | CDC4 | YFL009W |
| gi_696447656_gb_JRIS01000085.1_ | 20650 | G | A | 0 | 0.35 | 0.15 | 1 | 0.04 | 0.33 | synonymous_variant | CDC4 | YFL009W |
| gi_696447656_gb_JRIS01000085.1_ | 20656 | T | C | 0 | 0.37 | 0.10 | 1 | 0.04 | 0.15 | synonymous_variant | CDC4 | YFL009W |
| gi_696447656_gb_JRIS01000085.1_ | 20658 | G | A | 0 | 0.33 | 0.10 | 1 | 0.04 | 0.13 | synonymous_variant | CDC4 | YFL009W |
| gi_696447656_gb_JRIS01000085.1_ | 20691 | A | G | 0 | 0.12 | 0.06 | 1 | 0.03 | 0.12 | synonymous_variant | CDC4 | YFL009W |
| gi_696447656_gb_JRIS01000085.1_ | 20715 | T | C | 0 | 0.12 | 0.04 | 1 | 0.71 | 0.10 | missense_variant | CDC4 | YFL009W |
| gi_696447656_gb_JRIS01000085.1_ | 20717 | G | A | 0 | 0.17 | 0.04 | 1 | 0.04 | 0.10 | missense_variant | CDC4 | YFL009W |
| gi_696447656_gb_JRIS01000085.1_ | 20751 | T | A | 0 | 0.08 | 0.04 | 1 | 0.02 | 0.11 | missense_variant | CDC4 | YFL009W |
| gi_696447656_gb_JRIS01000085.1_ | 20752 | C | T | 0 | 0.07 | 0.03 | 1 | 0.02 | 0.11 | synonymous_variant | CDC4 | YFL009W |
| gi_696447656_gb_JRIS01000085.1_ | 20756 | G | A | 0 | 0.07 | 0.04 | 1 | 0.02 | 0.11 | missense_variant | CDC4 | YFL009W |
| gi_696447646_gb_JRIS01000088.1_ | 46918 | G | A | 1 | 0.32 | 1.00 | 0 | 1.00 | 0.91 | upstream_gene_variant | CBF2 | YGR140W |
| gi_696447632_gb_JRIS01000092.1_ | 1848 | A | G | 1 | 0.40 | 1.00 | 0 | 1.00 | 0.91 | downstream_gene_variant | YNR066C | YNR066C |
| gi_696447626_gb_JRIS01000094.1_ | 17777 | TTTNNNNNNNNNNNNNNNNNNN | T | 1 | 0.10 | 0.54 | 0 | 1.00 | 0.17 | upstream_gene_variant | RPA34 | YJL148W |
| gi_696447615_gb_JRIS01000097.1_ | 24314 | TCCCCC | TCCCCCCCCCCCC | 1 | 0.05 | 0.13 | 0 | 0.93 | 0.15 | upstream_gene_variant | DMC1 | YER179W |
| gi_696447601_gb_JRIS01000101.1_ | 48 | T | C | 0 | 1.00 | 0.00 | 1 | 0.04 | 0.25 | upstream_gene_variant | | UNDEF199 |
| gi_696447592_gb_JRIS01000103.1_ | 68377 | T | A | 0 | 1.00 | 0.00 | 1 | 0.00 | 1.00 | upstream_gene_variant | RTT105 | YER104W |
| gi_696447592_gb_JRIS01000103.1_ | 224570 | C | T | 1 | 0.00 | 1.00 | 0 | 1.00 | 0.00 | missense_variant | GCD11 | YER025W |
| gi_696447536_gb_JRIS01000118.1_ | 15018 | C | A | 1 | 0.00 | 1.00 | 0 | 1.00 | 0.00 | missense_variant | DYN1 | YKR054C |
| gi_696447526_gb_JRIS01000121.1_ | 77 | G | A | 0 | 0.02 | 0.01 | 1 | 0.00 | 0.03 | upstream_gene_variant | YOL166C | YOL166C |
| gi_696447526_gb_JRIS01000121.1_ | 95 | G | A | 0 | 0.02 | 0.01 | 1 | 0.00 | 0.01 | upstream_gene_variant | YOL166C | YOL166C |
| gi_696447464_gb_JRIS01000138.1_ | 81304 | T | TAATTTGAAAAAAAAAAAAAAAAAAAAAAAAAACTA<br>G | 0 | 0.81 | 0.17 | 1 | 0.00 | 0.05 | upstream_gene_variant | RRG7 | YOR305W |
| gi_696447394_gb_JRIS01000158.1_ | 454036 | C | G | 0 | 0.19 | 0.05 | 1 | 0.13 | 0.35 | upstream_gene_variant | ITR2 | YOL103W |
| gi_696447304_gb_JRIS01000182.1_ | 177 | A | G | 1 | 0.25 | 0.65 | 0 | 1.00 | 0.06 | synonymous_variant | SSB1 | YDL229W |
| gi_696447304_gb_JRIS01000182.1_ | 192 | G | A | 1 | 0.22 | 0.69 | 0 | 1.00 | 0.08 | synonymous_variant | SSB1 | YDL229W |
| gi_696447304_gb_JRIS01000182.1_ | 209 | A | G | 1 | 0.23 | 0.66 | 0 | 0.85 | 0.07 | synonymous_variant | SSB1 | YDL229W |
| gi_696447304_gb_JRIS01000182.1_ | 234 | G | A | 1 | 0.11 | 0.25 | 0 | 0.28 | 0.06 | synonymous_variant | SSB1 | YDL229W |
| gi_696447304_gb_JRIS01000182.1_ | 321 | A | G | 1 | 0.05 | 0.11 | 0 | 0.14 | 0.04 | synonymous_variant | SSB1 | YDL229W |
| gi_696447304_gb_JRIS01000182.1_ | 489 | G | A | 0 | 0.02 | 0.02 | 1 | 0.02 | 0.05 | synonymous_variant | SSB1 | YDL229W |
| gi_696447304_gb_JRIS01000182.1_ | 579 | A | T | 1 | 0.00 | 0.01 | 0 | 0.00 | 0.00 | synonymous_variant | SSB1 | YDL229W |
| gi_696447300_gb_JRIS01000183.1_ | 5253 | A | G | 1 | 0.00 | 1.00 | 0 | 1.00 | 0.00 | missense_variant | PPX1 | YHR201C |
| gi_696447292_gb_JRIS01000185.1_ | 3281 | G | A | 0 | 1.00 | 0.91 | 1 | 0.04 | 1.00 | upstream_gene_variant | YPS5 | YGL259W |
| gi_696447292_gb_JRIS01000185.1_ | 3292 | C | T | 0 | 1.00 | 0.91 | 1 | 0.08 | 1.00 | upstream_gene_variant | YPS5 | YGL259W |
| gi_696447257_gb_JRIS01000194.1_ | 24723 | A | G | 0 | 1.00 | 0.93 | 1 | 0.21 | 1.00 | synonymous_variant | YNR065C | YNR065C |
| gi_696447249_gb_JRIS01000196.1_ | 187624 | G | GAAAAAAAAAAAAAAAAAAAAAAAAAAAA | 0 | 0.07 | 0.03 | 1 | 0.00 | 0.06 | upstream_gene_variant | SPC3 | YLR066W |
| gi_696447197_gb_JRIS01000210.1_ | 103 | T | G | 0 | 1.00 | 0.03 | 1 | 0.05 | 0.16 | upstream_gene_variant | YHL034C | YHL034C |
| gi_696447193_gb_JRIS01000211.1_ | 8034 | AAANNNNNNNNNNNNNNNNNNNNNNN | A | 0 | 1.00 | 0.26 | 1 | 0.14 | 1.00 | upstream_gene_variant | VMA1 | YDL185W |
| gi_696447136_gb_JRIS01000226.1_ | 7 | C | T | 0 | 0.87 | 0.08 | 1 | 0.07 | 1.00 | upstream_gene_variant | ARN2 | YHL047C |
| gi_696446798_gb_JRIS01000321.1_ | 4918 | C | A | 0 | 0.02 | 0.01 | 1 | 0.02 | 0.07 | upstream_gene_variant | PHO12 | YHR215W |
| gi_696446726_gb_JRIS01000339.1_ | 84 | C | G | 0 | 0.05 | 0.04 | 1 | 0.09 | 0.30 | upstream_gene_variant | YMR326C | YMR326C |
| gi_696446714_gb_JRIS01000342.1_ | 401 | T | C | 0 | 0.13 | 0.06 | 1 | 0.04 | 0.11 | upstream_gene_variant | MAL12 | YGR292W |
| gi_696446714_gb_JRIS01000342.1_ | 406 | C | T | 0 | 0.12 | 0.06 | 1 | 0.04 | 0.11 | upstream_gene_variant | MAL12 | YGR292W |

**Supplementary Table 8: GATK variant calling against the BY4741 draft genome.** Columns 1 and 2 detail the position of the variant. Columns 3 and 4 note the reference allele (Genotype 0) and the alternative alleles (Genotypes 1 and 2). Colums 5-6 and 7-8 report the genotype called and the confidence associated with the call for strain 1691 and 1693 respectivly (confidence is out of 100). Blank rows indicate that a genotype was not called for that strain. Columns 9-11 report the location of the variant and its effect. For the sake of brevity, those variants which are shared between 1691 and 1693 and 226 variants called in intergenic regions are not shown. Discordant variants with high confidence (99 for both calls) are shaded in light grey. Variants that were also called by Samtools (see supplementary table 7) are shaded in dark grey (each of these was also high confidence).

| | | | | 1691 | | 1693 | | | | |
|---|---|---|---|---|---|---|---|---|---|---|
| Scaffold Genbank ID | POS | Genotype 0 | Genotype 1, 2 | Genotype | Confidence | Genotype | Confidence | Annotation | Gene | Gene ID |
| gi\|696447945\|gb\|JRIS01000005.1\| | 22253 | G | T | 0 | 99 | 1 | 99 | missense_variant | CDC1 | YDR182W |
| gi\|696447920\|gb\|JRIS01000013.1\| | 36866 | T | A | 0 | 99 | 1 | 99 | synonymous_variant | LYP1 | YNL268W |
| gi\|696447920\|gb\|JRIS01000013.1\| | 36872 | T | C | 0 | 99 | 1 | 99 | synonymous_variant | LYP1 | YNL268W |
| gi\|696447920\|gb\|JRIS01000013.1\| | 36889 | A | G | | | 1 | 99 | missense_variant | LYP1 | YNL268W |
| gi\|696447920\|gb\|JRIS01000013.1\| | 36893 | G | A | | | 1 | 99 | synonymous_variant | LYP1 | YNL268W |
| gi\|696447920\|gb\|JRIS01000013.1\| | 87004 | C | T | 1 | 99 | 0 | 99 | missense_variant | SLA2 | YNL243W |
| gi\|696447894\|gb\|JRIS01000021.1\| | 99 | G | A | | | 1 | 86 | upstream_gene_variant | | UNDEF10 |
| gi\|696447894\|gb\|JRIS01000021.1\| | 162 | T | C | | | 1 | 99 | upstream_gene_variant | | UNDEF10 |
| gi\|696447894\|gb\|JRIS01000021.1\| | 6781 | T | C | 1 | 99 | 0 | 99 | synonymous_variant | | UNDEF12 |
| gi\|696447894\|gb\|JRIS01000021.1\| | 10355 | C | CTA | 1 | 99 | 0 | 99 | upstream_gene_variant | | UNDEF18 |
| gi\|696447881\|gb\|JRIS01000023.1\| | 10511 | A | G | | | 1 | 99 | synonymous_variant | PAU5 | YFL020C |
| gi\|696447881\|gb\|JRIS01000023.1\| | 10517 | T | A | | | 1 | 99 | synonymous_variant | PAU5 | YFL020C |
| gi\|696447881\|gb\|JRIS01000023.1\| | 10522 | T | C | | | 1 | 99 | missense_variant | PAU5 | YFL020C |
| gi\|696447881\|gb\|JRIS01000023.1\| | 10532 | A | G | | | 1 | 99 | synonymous_variant | PAU5 | YFL020C |
| gi\|696447881\|gb\|JRIS01000023.1\| | 10535 | A | G | | | 1 | 99 | synonymous_variant | PAU5 | YFL020C |
| gi\|696447881\|gb\|JRIS01000023.1\| | 10550 | G | A | 1 | 99 | 0 | 99 | synonymous_variant | PAU5 | YFL020C |
| gi\|696447881\|gb\|JRIS01000023.1\| | 10553 | A | G | | | 1 | 99 | synonymous_variant | PAU5 | YFL020C |
| gi\|696447881\|gb\|JRIS01000023.1\| | 10558 | G | A | 1 | 99 | 0 | 99 | stop_gained | PAU5 | YFL020C |
| gi\|696447881\|gb\|JRIS01000023.1\| | 10562 | G | C,A | 2 | 99 | 1 | 99 | synonymous_variant | PAU5 | YFL020C |
| gi\|696447881\|gb\|JRIS01000023.1\| | 10563 | G | A | | | 1 | 99 | missense_variant | PAU5 | YFL020C |
| gi\|696447881\|gb\|JRIS01000023.1\| | 10564 | A | T | | | 1 | 99 | missense_variant | PAU5 | YFL020C |
| gi\|696447881\|gb\|JRIS01000023.1\| | 10573 | C | G | | | 1 | 99 | missense_variant | PAU5 | YFL020C |
| gi\|696447881\|gb\|JRIS01000023.1\| | 10574 | A | G | | | 1 | 99 | synonymous_variant | PAU5 | YFL020C |
| gi\|696447881\|gb\|JRIS01000023.1\| | 10580 | A | G | | | 1 | 99 | synonymous_variant | PAU5 | YFL020C |
| gi\|696447881\|gb\|JRIS01000023.1\| | 10595 | G | A | | | 1 | 99 | synonymous_variant | PAU5 | YFL020C |
| gi\|696447881\|gb\|JRIS01000023.1\| | 10598 | A | G | | | 1 | 99 | synonymous_variant | PAU5 | YFL020C |
| gi\|696447881\|gb\|JRIS01000023.1\| | 10601 | A | G | | | 1 | 99 | synonymous_variant | PAU5 | YFL020C |
| gi\|696447881\|gb\|JRIS01000023.1\| | 10604 | G | A | 1 | 99 | 0 | 99 | synonymous_variant | PAU5 | YFL020C |
| gi\|696447875\|gb\|JRIS01000025.1\| | 49526 | C | T | | | 1 | 99 | missense_variant | RPL2B | YIL018W |
| gi\|696447875\|gb\|JRIS01000025.1\| | 49532 | T | C | | | 1 | 99 | missense_variant | RPL2B | YIL018W |
| gi\|696447875\|gb\|JRIS01000025.1\| | 49544 | A | G | | | 1 | 99 | missense_variant | RPL2B | YIL018W |
| gi\|696447858\|gb\|JRIS01000030.1\| | 83846 | T | A | 0 | 99 | 1 | 99 | downstream_gene_variant | STT4 | YLR305C |
| gi\|696447858\|gb\|JRIS01000030.1\| | 83863 | G | GACGAACAAAGACACTGTTGGAATAAAAATCCACTATTGTCTATCAACTAATACTTATACTATCAATATATTATAATAGACGGCGTTAGATGATGACATAAGTTATGAGAAGATGTCATCGAGGTTA | 0 | 99 | 1 | 99 | downstream_gene_variant | STT4 | YLR305C |
| gi\|696447847\|gb\|JRIS01000034.1\| | 898 | A | T | | | 1 | 99 | upstream_gene_variant | YAL065C | YAL065C |
| gi\|696447821\|gb\|JRIS01000042.1\| | 106687 | G | T | 1 | 99 | 0 | 99 | upstream_gene_variant | ATG11 | YPR049C |
| gi\|696447821\|gb\|JRIS01000042.1\| | 140522 | C | T | | | 1 | 99 | synonymous_variant | TEF1 | YPR080W |
| gi\|696447799\|gb\|JRIS01000047.1\| | 950 | C | T | | | 1 | 90 | synonymous_variant | HSP82 | YPL240C |
| gi\|696447799\|gb\|JRIS01000047.1\| | 971 | C | T | 0 | 43 | 1 | 90 | synonymous_variant | HSP82 | YPL240C |
| gi\|696447795\|gb\|JRIS01000048.1\| | 99699 | A | C | 1 | 99 | 0 | 99 | missense_variant | AHT1 | YHR093W |
| gi\|696447795\|gb\|JRIS01000048.1\| | 158738 | G | A | | | 1 | 99 | missense_variant | | UNDEF44 |
| gi\|696447795\|gb\|JRIS01000048.1\| | 158740 | A | G | | | 1 | 99 | synonymous_variant | | UNDEF44 |
| gi\|696447795\|gb\|JRIS01000048.1\| | 158785 | T | A | 1 | 99 | 0 | 99 | synonymous_variant | | UNDEF44 |
| gi\|696447795\|gb\|JRIS01000048.1\| | 158786 | G | A | 1 | 99 | 0 | 99 | missense_variant | | UNDEF44 |
| gi\|696447795\|gb\|JRIS01000048.1\| | 158789 | T | C | 1 | 99 | 0 | 99 | missense_variant | | UNDEF44 |
| gi\|696447795\|gb\|JRIS01000048.1\| | 158800 | C | T | 1 | 45 | 0 | 99 | synonymous_variant | | UNDEF44 |
| gi\|696447795\|gb\|JRIS01000048.1\| | 158805 | T | C | 1 | 45 | 0 | 99 | missense_variant | | UNDEF44 |
| gi\|696447795\|gb\|JRIS01000048.1\| | 158806 | A | G | 1 | 90 | 0 | 99 | synonymous_variant | | UNDEF44 |
| gi\|696447795\|gb\|JRIS01000048.1\| | 158812 | C | T | 1 | 99 | 0 | 99 | synonymous_variant | | UNDEF44 |
| gi\|696447795\|gb\|JRIS01000048.1\| | 158813 | T | G | 1 | 99 | 0 | 99 | missense_variant | | UNDEF44 |
| gi\|696447795\|gb\|JRIS01000048.1\| | 158816 | T | A | 1 | 99 | 0 | 99 | missense_variant | | UNDEF44 |
| gi\|696447795\|gb\|JRIS01000048.1\| | 158820 | C | T | 1 | 99 | 0 | 99 | missense_variant | | UNDEF44 |
| gi\|696447795\|gb\|JRIS01000048.1\| | 158821 | G | A | 1 | 99 | 0 | 99 | synonymous_variant | | UNDEF44 |
| gi\|696447795\|gb\|JRIS01000048.1\| | 158824 | C | G | 1 | 99 | 0 | 99 | synonymous_variant | | UNDEF44 |
| gi\|696447795\|gb\|JRIS01000048.1\| | 158831 | C | T | 1 | 99 | 0 | 99 | missense_variant | | UNDEF44 |
| gi\|696447795\|gb\|JRIS01000048.1\| | 158836 | G | C | 1 | 99 | 0 | 99 | synonymous_variant | | UNDEF44 |
| gi\|696447795\|gb\|JRIS01000048.1\| | 158840 | T | A | 1 | 99 | 0 | 99 | missense_variant | | UNDEF44 |
| gi\|696447795\|gb\|JRIS01000048.1\| | 158841 | T | G | 1 | 99 | 0 | 99 | missense_variant | | UNDEF44 |
| gi\|696447795\|gb\|JRIS01000048.1\| | 158843 | A | G | 1 | 99 | 0 | 99 | missense_variant | | UNDEF44 |
| gi\|696447795\|gb\|JRIS01000048.1\| | 158846 | C | G | 1 | 99 | 0 | 99 | missense_variant | | UNDEF44 |
| gi\|696447795\|gb\|JRIS01000048.1\| | 158857 | C | T | 1 | 99 | 0 | 99 | synonymous_variant | | UNDEF44 |
| gi\|696447795\|gb\|JRIS01000048.1\| | 158884 | T | C | 1 | 99 | 0 | 99 | synonymous_variant | | UNDEF44 |
| gi\|696447795\|gb\|JRIS01000048.1\| | 158893 | G | A | 1 | 99 | 0 | 99 | synonymous_variant | | UNDEF44 |
| gi\|696447795\|gb\|JRIS01000048.1\| | 158941 | A | G | 0 | 88 | 1 | 99 | synonymous_variant | | UNDEF44 |
| gi\|696447795\|gb\|JRIS01000048.1\| | 159298 | T | C | | | 1 | 88 | upstream_gene_variant | DYS1 | YHR068W |
| gi\|696447790\|gb\|JRIS01000049.1\| | 6924 | C | T | | | 1 | 67 | upstream_gene_variant | CDC53 | YDL132W |
| gi\|696447774\|gb\|JRIS01000053.1\| | 29830 | A | G | | | 1 | 99 | upstream_gene_variant | PHO87 | YCR037C |
| gi\|696447770\|gb\|JRIS01000054.1\| | 589 | G | A | | | 1 | 86 | missense_variant | | UNDEF50 |
| gi\|696447770\|gb\|JRIS01000054.1\| | 631 | T | G | | | 1 | 80 | missense_variant | | UNDEF50 |
| gi\|696447766\|gb\|JRIS01000055.1\| | 31006 | A | G | | | 1 | 99 | upstream_gene_variant | GPI2 | YPL076W |
| gi\|696447766\|gb\|JRIS01000055.1\| | 31015 | A | G | | | 1 | 99 | upstream_gene_variant | GPI2 | YPL076W |
| gi\|696447766\|gb\|JRIS01000055.1\| | 31018 | C | T | | | 1 | 99 | upstream_gene_variant | GPI2 | YPL076W |
| gi\|696447766\|gb\|JRIS01000055.1\| | 31030 | C | T | | | 1 | 99 | upstream_gene_variant | GPI2 | YPL076W |
| gi\|696447766\|gb\|JRIS01000055.1\| | 31042 | G | A | | | 1 | 99 | upstream_gene_variant | GPI2 | YPL076W |
| gi\|696447766\|gb\|JRIS01000055.1\| | 31051 | C | T | | | 1 | 99 | upstream_gene_variant | GPI2 | YPL076W |
| gi\|696447766\|gb\|JRIS01000055.1\| | 31054 | G | A | | | 1 | 99 | upstream_gene_variant | GPI2 | YPL076W |
| gi\|696447766\|gb\|JRIS01000055.1\| | 31060 | A | G | | | 1 | 99 | upstream_gene_variant | GPI2 | YPL076W |
| gi\|696447766\|gb\|JRIS01000055.1\| | 31069 | C | T | | | 1 | 99 | upstream_gene_variant | GPI2 | YPL076W |
| gi\|696447766\|gb\|JRIS01000055.1\| | 31071 | A | G | | | 1 | 99 | upstream_gene_variant | GPI2 | YPL076W |
| gi\|696447766\|gb\|JRIS01000055.1\| | 31075 | T | C | | | 1 | 99 | upstream_gene_variant | GPI2 | YPL076W |
| gi\|696447766\|gb\|JRIS01000055.1\| | 31078 | G | A | | | 1 | 99 | upstream_gene_variant | GPI2 | YPL076W |
| gi\|696447766\|gb\|JRIS01000055.1\| | 31084 | G | A | | | 1 | 99 | upstream_gene_variant | GPI2 | YPL076W |
| gi\|696447766\|gb\|JRIS01000055.1\| | 31126 | A | T | | | 1 | 99 | upstream_gene_variant | GPI2 | YPL076W |
| gi\|696447766\|gb\|JRIS01000055.1\| | 31129 | T | C | | | 1 | 99 | upstream_gene_variant | GPI2 | YPL076W |
| gi\|696447766\|gb\|JRIS01000055.1\| | 31141 | G | T | | | 1 | 99 | upstream_gene_variant | GPI2 | YPL076W |
| gi\|696447766\|gb\|JRIS01000055.1\| | 31156 | T | C | | | 1 | 99 | upstream_gene_variant | GPI2 | YPL076W |
| gi\|696447766\|gb\|JRIS01000055.1\| | 31162 | C | T | | | 1 | 99 | upstream_gene_variant | GPI2 | YPL076W |
| gi\|696447766\|gb\|JRIS01000055.1\| | 31980 | G | GA | 1 | 99 | | | upstream_gene_variant | MOT1 | YPL082C |
| gi\|696447735\|gb\|JRIS01000063.1\| | 130139 | C | CTCTTTTTTTTTTTTTTTTTTTTTTTTTT,CTCTTTTTTTTTTTTTTTTTTTTTTTT | 2 | 99 | 1 | 99 | upstream_gene_variant | PET9 | YBL030C |
| gi\|696447727\|gb\|JRIS01000065.1\| | 653 | G | C | 1 | 64 | | | downstream_gene_variant | YIL169C | YIL169C |
| gi\|696447717\|gb\|JRIS01000068.1\| | 62054 | C | CAACAAAAAAAAAAAAAAAAAAAAAAAAAAAAAAAGA,CAACAAAAAAAAAAAAAAAAAAAAAAAAAAAAAAGA | 2 | 88 | 1 | 99 | upstream_gene_variant | MET28 | YIR017C |
| gi\|696447713\|gb\|JRIS01000069.1\| | 149 | G | A | | | 1 | 99 | upstream_gene_variant | MRH4 | YGL064C |
| gi\|696447713\|gb\|JRIS01000069.1\| | 246160 | T | A,TA | 2 | 88 | 1 | 31 | upstream_gene_variant | COX13 | YGL191W |
| gi\|696447699\|gb\|JRIS01000073.1\| | 885 | G | A | 1 | 99 | | | synonymous_variant | PAU17 | YLL025W |
| gi\|696447699\|gb\|JRIS01000073.1\| | 921 | A | G | 1 | 99 | | | synonymous_variant | PAU17 | YLL025W |
| gi\|696447699\|gb\|JRIS01000073.1\| | 924 | T | C | 1 | 99 | | | synonymous_variant | PAU17 | YLL025W |
| gi\|696447699\|gb\|JRIS01000073.1\| | 939 | A | G,T | 2 | 99 | 1 | 99 | synonymous_variant | PAU17 | YLL025W |
| gi\|696447699\|gb\|JRIS01000073.1\| | 945 | G | A | 1 | 99 | | | synonymous_variant | PAU17 | YLL025W |
| gi\|696447699\|gb\|JRIS01000073.1\| | 950 | T | A | 1 | 99 | | | missense_variant | PAU17 | YLL025W |
| gi\|696447699\|gb\|JRIS01000073.1\| | 951 | A | G | 1 | 99 | | | synonymous_variant | PAU17 | YLL025W |
| gi\|696447699\|gb\|JRIS01000073.1\| | 959 | C | G | 1 | 99 | | | missense_variant | PAU17 | YLL025W |
| gi\|696447699\|gb\|JRIS01000073.1\| | 981 | A | G | 1 | 99 | | | synonymous_variant | PAU17 | YLL025W |
| gi\|696447699\|gb\|JRIS01000073.1\| | 990 | G | A | 1 | 99 | | | synonymous_variant | PAU17 | YLL025W |
| gi\|696447699\|gb\|JRIS01000073.1\| | 996 | T | C | 1 | 99 | | | synonymous_variant | PAU17 | YLL025W |
| gi\|696447699\|gb\|JRIS01000073.1\| | 1002 | A | G | 1 | 99 | 0 | 49 | synonymous_variant | PAU17 | YLL025W |
| gi\|696447699\|gb\|JRIS01000073.1\| | 1007 | G | A | 1 | 99 | | | synonymous_variant | PAU17 | YLL025W |
| gi\|696447695\|gb\|JRIS01000074.1\| | 1052 | G | A | | | 1 | 99 | missense_variant | YNL019C | YNL019C |
| gi\|696447695\|gb\|JRIS01000074.1\| | 1172 | C | T | 0 | 99 | 1 | 99 | missense_variant&splice_region_variant | YNL019C | YNL019C |
| gi\|696447695\|gb\|JRIS01000074.1\| | 1371 | T | C | 0 | 99 | 1 | 99 | upstream_gene_variant | YNL019C | YNL019C_circ |
| gi\|696447695\|gb\|JRIS01000074.1\| | 1433 | T | G | 0 | 99 | 1 | 61 | upstream_gene_variant | YNL019C | YNL019C_circ |
| gi\|696447695\|gb\|JRIS01000074.1\| | 1624 | A | G | 1 | 99 | | | upstream_gene_variant | YNL019C | YNL019C_circ |
| gi\|696447695\|gb\|JRIS01000074.1\| | 1744 | A | G | 0 | 99 | 1 | 99 | upstream_gene_variant | YNL019C | YNL019C_circ |
| gi\|696447695\|gb\|JRIS01000074.1\| | 1797 | G | A | 1 | 99 | 0 | 99 | upstream_gene_variant | YNL019C | YNL019C_circ |
| gi\|696447695\|gb\|JRIS01000074.1\| | 1811 | T | C | 1 | 99 | 0 | 27 | upstream_gene_variant | YNL019C | YNL019C_circ |
| gi\|696447695\|gb\|JRIS01000074.1\| | 1855 | C | T | 1 | 99 | 0 | 99 | upstream_gene_variant | YNL019C | YNL019C_circ |
| gi\|696447695\|gb\|JRIS01000074.1\| | 1977 | T | G | 0 | 65 | 1 | 99 | upstream_gene_variant | YNL019C | YNL019C_circ |
| gi\|696447695\|gb\|JRIS01000074.1\| | 1978 | A | G | 0 | 8 | 1 | 99 | upstream_gene_variant | YNL019C | YNL019C_circ |
| gi\|696447695\|gb\|JRIS01000074.1\| | 1988 | C | G | 0 | 14 | 1 | 90 | upstream_gene_variant | YNL019C | YNL019C_circ |
| gi\|696447695\|gb\|JRIS01000074.1\| | 2004 | A | G | 0 | 95 | 1 | 45 | upstream_gene_variant | YNL019C | YNL019C_circ |
| gi\|696447695\|gb\|JRIS01000074.1\| | 2066 | C | T | | | 1 | 43 | upstream_gene_variant | YNL019C | YNL019C |
| gi\|696447692\|gb\|JRIS01000075.1\| | 64342 | C | T | 1 | 99 | 0 | 99 | upstream_gene_variant | ALE1 | YOR175C |
| gi\|696447667\|gb\|JRIS01000082.1\| | 24360 | G | A | 0 | 99 | 1 | 83 | upstream_gene_variant | YGR111W | YGR111W |
| gi\|696447667\|gb\|JRIS01000082.1\| | 24364 | C | T | 0 | 99 | 1 | 80 | upstream_gene_variant | YGR111W | YGR111W |
| gi\|696447667\|gb\|JRIS01000082.1\| | 25927 | G | A | 0 | 99 | 1 | 47 | upstream_gene_variant | CLD1 | YGR110W |
| gi\|696447667\|gb\|JRIS01000082.1\| | 26011 | G | A | 1 | 99 | 0 | 93 | upstream_gene_variant | CLD1 | YGR110W |
| gi\|696447667\|gb\|JRIS01000082.1\| | 26930 | A | G | 1 | 41 | 0 | 99 | upstream_gene_variant | CAB2 | YIL083C |
| gi\|696447667\|gb\|JRIS01000082.1\| | 27524 | C | T | 0 | 99 | 1 | 99 | synonymous_variant | YIL082W | YIL082W |
| gi\|696447659\|gb\|JRIS01000084.1\| | 56773 | T | G | 0 | 99 | 1 | 99 | missense_variant | TOR2 | YKL203C |
| gi\|696447656\|gb\|JRIS01000085.1\| | 20480 | T | C | 1 | 53 | 0 | 65 | missense_variant | CDC4 | YFL009W |
| gi\|696447656\|gb\|JRIS01000085.1\| | 33894 | ATTTTTTTTTTTTTTTTTTTTTTTTTTTTTTTTTTTTTT | A | 0 | 99 | 1 | 99 | upstream_gene_variant | HSP12 | YFL014W |
| gi\|696447656\|gb\|JRIS01000085.1\| | 33952 | T | A | | | 1 | 99 | upstream_gene_variant | HSP12 | YFL014W |
| gi\|696447656\|gb\|JRIS01000085.1\| | 33956 | T | A | 0 | 99 | 1 | 99 | upstream_gene_variant | HSP12 | YFL014W |
| gi\|696447646\|gb\|JRIS01000088.1\| | 46918 | G | A | 1 | 84 | . | . | upstream_gene_variant | CBF2 | YGR140W |
| gi\|696447632\|gb\|JRIS01000092.1\| | 1785 | C | T | 1 | 88 | | | downstream_gene_variant | YNR066C | YNR066C |
| gi\|696447632\|gb\|JRIS01000092.1\| | 1806 | C | T | 1 | 87 | | | downstream_gene_variant | YNR066C | YNR066C |
| gi\|696447632\|gb\|JRIS01000092.1\| | 1830 | A | G | 1 | 82 | | | downstream_gene_variant | YNR066C | YNR066C |
| gi\|696447632\|gb\|JRIS01000092.1\| | 1848 | A | G | 1 | 84 | | | downstream_gene_variant | YNR066C | YNR066C |
| gi\|696447629\|gb\|JRIS01000093.1\| | 63035 | C | CGGGGGGGGG | | | 1 | 90 | upstream_gene_variant | STE24 | YJR117W |
| gi\|696447626\|gb\|JRIS01000094.1\| | 33656 | A | AGTAGCTTGAACTTGACCATCACCGATTTGAGAAACGGCAGCAGCGGTGCTCTTTGGGGCTAAAGTAGTGGTG | 0 | 99 | 1 | 99 | disruptive_inframe_insertion | HSP150 | YJL159W |
| gi\|696447622\|gb\|JRIS01000095.1\| | 13665 | T | C | | | 1 | 99 | upstream_gene_variant | YNR073C | YNR073C |
| gi\|696447622\|gb\|JRIS01000095.1\| | 13685 | A | C | | | 1 | 99 | upstream_gene_variant | YNR073C | YNR073C |
| gi\|696447622\|gb\|JRIS01000095.1\| | 13690 | T | TC | | | 1 | 99 | upstream_gene_variant | YNR073C | YNR073C |
| gi\|696447622\|gb\|JRIS01000095.1\| | 13695 | C | T | | | 1 | 99 | upstream_gene_variant | YNR073C | YNR073C |
| gi\|696447622\|gb\|JRIS01000095.1\| | 13703 | G | A | | | 1 | 99 | upstream_gene_variant | YNR073C | YNR073C |
| gi\|696447622\|gb\|JRIS01000095.1\| | 13716 | G | C | | | 1 | 99 | upstream_gene_variant | YNR073C | YNR073C |
| gi\|696447622\|gb\|JRIS01000095.1\| | 13726 | T | C | | | 1 | 99 | upstream_gene_variant | YNR073C | YNR073C |
| gi\|696447622\|gb\|JRIS01000095.1\| | 13745 | C | T | | | 1 | 99 | upstream_gene_variant | YNR073C | YNR073C |
| gi\|696447622\|gb\|JRIS01000095.1\| | 13751 | C | T | | | 1 | 99 | upstream_gene_variant | YNR073C | YNR073C |
| gi\|696447622\|gb\|JRIS01000095.1\| | 13766 | G | C | | | 1 | 99 | upstream_gene_variant | YNR073C | YNR073C |
| gi\|696447622\|gb\|JRIS01000095.1\| | 13769 | T | G | | | 1 | 99 | upstream_gene_variant | YNR073C | YNR073C |
| gi\|696447622\|gb\|JRIS01000095.1\| | 13771 | A | T | | | 1 | 99 | upstream_gene_variant | YNR073C | YNR073C |
| gi\|696447622\|gb\|JRIS01000095.1\| | 13774 | C | T | | | 1 | 99 | upstream_gene_variant | YNR073C | YNR073C |
| gi\|696447622\|gb\|JRIS01000095.1\| | 13788 | C | T | | | 1 | 99 | upstream_gene_variant | YNR073C | YNR073C |
| gi\|696447622\|gb\|JRIS01000095.1\| | 13800 | G | A | | | 1 | 99 | upstream_gene_variant | YNR073C | YNR073C |
| gi\|696447622\|gb\|JRIS01000095.1\| | 13807 | C | G | | | 1 | 99 | upstream_gene_variant | YNR073C | YNR073C |
| gi\|696447622\|gb\|JRIS01000095.1\| | 13808 | T | G | | | 1 | 99 | upstream_gene_variant | YNR073C | YNR073C |
| gi\|696447622\|gb\|JRIS01000095.1\| | 13833 | C | CAA | | | 1 | 99 | upstream_gene_variant | YNR073C | YNR073C |
| gi\|696447622\|gb\|JRIS01000095.1\| | 14073 | A | G | | | 1 | 99 | synonymous_variant | YNR073C | YNR073C |
| gi\|696447622\|gb\|JRIS01000095.1\| | 14172 | T | C | | | 1 | 99 | synonymous_variant | YNR073C | YNR073C |
| gi\|696447609\|gb\|JRIS01000099.1\| | 1810 | A | G | 0 | 99 | 1 | 97 | upstream_gene_variant | TDA8 | YAL064C-A |
| gi\|696447605\|gb\|JRIS01000100.1\| | 33431 | T | C | 0 | 99 | 1 | 99 | upstream_gene_variant | EFT1 | YOR133W |
| gi\|696447592\|gb\|JRIS01000103.1\| | 68377 | T | A | 0 | 99 | 1 | 99 | upstream_gene_variant | RTT105 | YER104W |
| gi\|696447592\|gb\|JRIS01000103.1\| | 107515 | C | T | | | 1 | 99 | synonymous_variant | SER3 | YER081W |
| gi\|696447592\|gb\|JRIS01000103.1\| | 107518 | G | C | | | 1 | 99 | synonymous_variant | SER3 | YER081W |
| gi\|696447592\|gb\|JRIS01000103.1\| | 107532 | G | A | | | 1 | 99 | synonymous_variant | SER3 | YER081W |
| gi\|696447592\|gb\|JRIS01000103.1\| | 107563 | G | A | | | 1 | 99 | synonymous_variant | SER3 | YER081W |
| gi\|696447592\|gb\|JRIS01000103.1\| | 107581 | C | G | | | 1 | 99 | synonymous_variant | SER3 | YER081W |
| gi\|696447592\|gb\|JRIS01000103.1\| | 107584 | G | A | | | 1 | 99 | synonymous_variant | SER3 | YER081W |
| gi\|696447592\|gb\|JRIS01000103.1\| | 107599 | G | A | | | 1 | 99 | synonymous_variant | SER3 | YER081W |
| gi\|696447592\|gb\|JRIS01000103.1\| | 107608 | A | G | | | 1 | 99 | synonymous_variant | SER3 | YER081W |
| gi\|696447592\|gb\|JRIS01000103.1\| | 168974 | A | ATTTTTTTTTTTTTTTTTTTT,ATTTTTTTTTTTTTTTTTTTTT | 1 | 81 | 2 | 99 | upstream_gene_variant | PIC2 | YER053C |
| gi\|696447592\|gb\|JRIS01000103.1\| | 224570 | C | T | 1 | 99 | 0 | 99 | missense_variant | GCD11 | YER025W |

| | | | | | | | | | | |
|---|---|---|---|---|---|---|---|---|---|---|
| gi\|696447583\|gb\|JRIS01000105.1\| | 14205 | T | C | 1 | 99 | | | upstream_gene_variant | NTC20 | YBR188C |
| gi\|696447583\|gb\|JRIS01000105.1\| | 14209 | A | C | 1 | 99 | | | upstream_gene_variant | NTC20 | YBR188C |
| gi\|696447583\|gb\|JRIS01000105.1\| | 14211 | G | T | 1 | 99 | | | upstream_gene_variant | NTC20 | YBR188C |
| gi\|696447583\|gb\|JRIS01000105.1\| | 14214 | C | T | 1 | 99 | | | upstream_gene_variant | NTC20 | YBR188C |
| gi\|696447583\|gb\|JRIS01000105.1\| | 14217 | C | T | 1 | 99 | | | upstream_gene_variant | NTC20 | YBR188C |
| gi\|696447583\|gb\|JRIS01000105.1\| | 14227 | A | G | 1 | 99 | | | upstream_gene_variant | NTC20 | YBR188C |
| gi\|696447574\|gb\|JRIS01000107.1\| | 47417 | TA | T | 1 | 99 | 0 | 99 | upstream_gene_variant | YSH1 | YLR277C |
| gi\|696447565\|gb\|JRIS01000110.1\| | 160312 | T | C | 0 | 99 | 1 | 59 | upstream_gene_variant | YNL115C | YNL115C |
| gi\|696447565\|gb\|JRIS01000110.1\| | 161918 | T | TC | 0 | 99 | 1 | 86 | upstream_gene_variant | YNL115C | YNL115C |
| gi\|696447565\|gb\|JRIS01000110.1\| | 161945 | C | T | | | 1 | 45 | upstream_gene_variant | YNL115C | YNL115C |
| gi\|696447550\|gb\|JRIS01000114.1\| | 1163 | T | TTATATAAA | 1 | 99 | | | upstream_gene_variant | CDS1 | YBR029C |
| gi\|696447536\|gb\|JRIS01000118.1\| | 15018 | C | A | 1 | 99 | 0 | 99 | missense_variant | DYN1 | YKR054C |
| gi\|696447536\|gb\|JRIS01000118.1\| | 32097 | GA | G | 1 | 80 | 0 | 99 | upstream_gene_variant | TRK2 | YKR050W |
| gi\|696447526\|gb\|JRIS01000121.1\| | 622 | T | A | 1 | 99 | | | synonymous_variant | YOL166C | YOL166C |
| gi\|696447478\|gb\|JRIS01000134.1\| | 42257 | G | GAAAAAAAAAAAAAAAAACACAGAAAAGAATGCAGAAAGATGTCAACTGAAAAAAAAAAAAGGTGAACACAGGAAAAAAAAT | | | 1 | 99 | upstream_gene_variant | SSC1 | YJR045C |
| gi\|696447464\|gb\|JRIS01000138.1\| | 16156 | T | TCGAAAAAAAAAAAAAAAAAAAAAAAAGAAAAGAAGC,TCGAAAAAAAAAAAAAAAAAAAAAAAAAAAAAGAAAAGAAGC | 2 | 99 | 1 | 98 | upstream_gene_variant | YOR338W | YOR338W |
| gi\|696447464\|gb\|JRIS01000138.1\| | 25814 | A | AGAAG | 1 | 99 | 0 | 99 | upstream_gene_variant | MRS2 | YOR334W |
| gi\|696447464\|gb\|JRIS01000138.1\| | 25825 | A | AGAGAC | 1 | 99 | 0 | 99 | upstream_gene_variant | MRS2 | YOR334W |
| gi\|696447456\|gb\|JRIS01000140.1\| | 23490 | A | T | | | 1 | 99 | downstream_gene_variant | ADD37 | YMR184W |
| gi\|696447456\|gb\|JRIS01000140.1\| | 23493 | G | A | | | 1 | 99 | downstream_gene_variant | ADD37 | YMR184W |
| gi\|696447456\|gb\|JRIS01000140.1\| | 23499 | T | C | | | 1 | 99 | downstream_gene_variant | ADD37 | YMR184W |
| gi\|696447456\|gb\|JRIS01000140.1\| | 23517 | G | A | | | 1 | 99 | downstream_gene_variant | ADD37 | YMR184W |
| gi\|696447456\|gb\|JRIS01000140.1\| | 23523 | G | T | | | 1 | 99 | downstream_gene_variant | ADD37 | YMR184W |
| gi\|696447456\|gb\|JRIS01000140.1\| | 23526 | C | T | | | 1 | 99 | downstream_gene_variant | ADD37 | YMR184W |
| gi\|696447456\|gb\|JRIS01000140.1\| | 23532 | A | T | | | 1 | 99 | downstream_gene_variant | ADD37 | YMR184W |
| gi\|696447456\|gb\|JRIS01000140.1\| | 23539 | T | C | | | 1 | 99 | downstream_gene_variant | ADD37 | YMR184W |
| gi\|696447456\|gb\|JRIS01000140.1\| | 23540 | TAG | T | 0 | 99 | 1 | 99 | downstream_gene_variant | ADD37 | YMR184W |
| gi\|696447456\|gb\|JRIS01000140.1\| | 23544 | C | CTA | | | 1 | 99 | downstream_gene_variant | ADD37 | YMR184W |
| gi\|696447456\|gb\|JRIS01000140.1\| | 23559 | T | C | | | 1 | 99 | downstream_gene_variant | ADD37 | YMR184W |
| gi\|696447456\|gb\|JRIS01000140.1\| | 23850 | C | T | | | 1 | 51 | downstream_gene_variant | ADD37 | YMR184W |
| gi\|696447451\|gb\|JRIS01000142.1\| | 39391 | GA | G | 0 | 99 | 1 | 60 | upstream_gene_variant | MRS6 | YOR370C |
| gi\|696447449\|gb\|JRIS01000143.1\| | 20050 | A | T | 1 | 99 | | | upstream_gene_variant | ARP8 | YOR141C |
| gi\|696447444\|gb\|JRIS01000145.1\| | 308 | T | C | | | 1 | 99 | synonymous_variant | YCR102C | YCR102C |
| gi\|696447444\|gb\|JRIS01000145.1\| | 318 | A | G | | | 1 | 99 | missense_variant | YCR102C | YCR102C |
| gi\|696447444\|gb\|JRIS01000145.1\| | 12282 | AAT | A | 0 | 99 | 1 | 99 | upstream_gene_variant | YCR097W | YCR097W |
| gi\|696447433\|gb\|JRIS01000148.1\| | 92199 | A | AT | 1 | 99 | | | upstream_gene_variant | SHE3 | YBR130C |
| gi\|696447433\|gb\|JRIS01000148.1\| | 113252 | A | G | | | 1 | 99 | downstream_gene_variant | GRS1 | YBR121C |
| gi\|696447429\|gb\|JRIS01000149.1\| | 47506 | A | ATAACT | 0 | 99 | 1 | 99 | upstream_gene_variant | RTN2 | YDL204W |
| gi\|696447429\|gb\|JRIS01000149.1\| | 47510 | A | AGAT | 0 | 99 | 1 | 99 | upstream_gene_variant | RTN2 | YDL204W |
| gi\|696447429\|gb\|JRIS01000149.1\| | 47513 | A | AGGAGATAAGG | 0 | 99 | 1 | 99 | upstream_gene_variant | RTN2 | YDL204W |
| gi\|696447394\|gb\|JRIS01000158.1\| | 133106 | GA | G | 1 | 55 | 0 | 99 | upstream_gene_variant | MSA1 | YOR066W |
| gi\|696447394\|gb\|JRIS01000158.1\| | 261800 | C | CTT | 1 | 99 | | | upstream_gene_variant | SIN3 | YOL004W |
| gi\|696447367\|gb\|JRIS01000165.1\| | 96765 | G | GA | 1 | 99 | 0 | 99 | upstream_gene_variant | VMS1 | YDR049W |
| gi\|696447367\|gb\|JRIS01000165.1\| | 100581 | A | T | 0 | 99 | 1 | 99 | upstream_gene_variant | HEM13 | YDR044W |
| gi\|696447353\|gb\|JRIS01000169.1\| | 9116 | GA | G | 1 | 99 | | | upstream_gene_variant | ERG11 | YHR007C |
| gi\|696447326\|gb\|JRIS01000175.1\| | 75535 | T | A | | | 1 | 90 | missense_variant | ENO1 | YGR254W |
| gi\|696447326\|gb\|JRIS01000175.1\| | 75536 | G | A | | | 1 | 90 | synonymous_variant | ENO1 | YGR254W |
| gi\|696447317\|gb\|JRIS01000178.1\| | 53039 | A | G | | | 1 | 99 | synonymous_variant | RPL4A | YBR031W |
| gi\|696447304\|gb\|JRIS01000182.1\| | 123 | G | A | | | 1 | 99 | synonymous_variant | SSB1 | YDL229W |
| gi\|696447304\|gb\|JRIS01000182.1\| | 234 | G | A | 1 | 81 | 0 | 99 | synonymous_variant | SSB1 | YDL229W |
| gi\|696447304\|gb\|JRIS01000182.1\| | 249 | G | A | 1 | 57 | 0 | 99 | synonymous_variant | SSB1 | YDL229W |
| gi\|696447304\|gb\|JRIS01000182.1\| | 312 | A | G | 1 | 99 | 0 | 99 | synonymous_variant | SSB1 | YDL229W |
| gi\|696447304\|gb\|JRIS01000182.1\| | 321 | A | G | 1 | 99 | 0 | 99 | synonymous_variant | SSB1 | YDL229W |
| gi\|696447304\|gb\|JRIS01000182.1\| | 489 | G | A | 0 | 41 | 1 | 99 | synonymous_variant | SSB1 | YDL229W |
| gi\|696447304\|gb\|JRIS01000182.1\| | 537 | AG | A | 1 | 99 | 0 | 99 | frameshift_variant | SSB1 | YDL229W |
| gi\|696447304\|gb\|JRIS01000182.1\| | 539 | C | CTG | 1 | 99 | 0 | 99 | frameshift_variant | SSB1 | YDL229W |
| gi\|696447304\|gb\|JRIS01000182.1\| | 542 | AT | A | 1 | 99 | 0 | 99 | frameshift_variant | SSB1 | YDL229W |
| gi\|696447304\|gb\|JRIS01000182.1\| | 546 | A | T | 1 | 99 | 0 | 99 | synonymous_variant | SSB1 | YDL229W |
| gi\|696447304\|gb\|JRIS01000182.1\| | 549 | A | G | 1 | 99 | 0 | 99 | synonymous_variant | SSB1 | YDL229W |
| gi\|696447304\|gb\|JRIS01000182.1\| | 579 | A | T | 1 | 99 | 0 | 99 | synonymous_variant | SSB1 | YDL229W |
| gi\|696447304\|gb\|JRIS01000182.1\| | 591 | A | G | 1 | 90 | 0 | 99 | synonymous_variant | SSB1 | YDL229W |
| gi\|696447304\|gb\|JRIS01000182.1\| | 693 | G | A | 1 | 99 | 0 | 99 | synonymous_variant | SSB1 | YDL229W |
| gi\|696447300\|gb\|JRIS01000183.1\| | 5253 | A | G | 1 | 99 | 0 | 99 | missense_variant | PPX1 | YHR201C |
| gi\|696447292\|gb\|JRIS01000185.1\| | 3263 | A | ATC | | | 1 | 99 | upstream_gene_variant | YPS5 | YGL259W |
| gi\|696447292\|gb\|JRIS01000185.1\| | 3281 | G | A | | | 1 | 99 | upstream_gene_variant | YPS5 | YGL259W |
| gi\|696447292\|gb\|JRIS01000185.1\| | 3292 | C | T | | | 1 | 99 | upstream_gene_variant | YPS5 | YGL259W |
| gi\|696447289\|gb\|JRIS01000186.1\| | 7587 | C | T | | | 1 | 99 | synonymous_variant | ENA5 | YDR038C |
| gi\|696447289\|gb\|JRIS01000186.1\| | 7613 | G | T | | | 1 | 99 | missense_variant | ENA5 | YDR038C |
| gi\|696447282\|gb\|JRIS01000188.1\| | 762 | C | T | | | 1 | 99 | upstream_gene_variant | MDM36 | YPR083W |
| gi\|696447274\|gb\|JRIS01000190.1\| | 519 | C | T | | | 1 | 99 | synonymous_variant | LYS20 | YDL182W |
| gi\|696447274\|gb\|JRIS01000190.1\| | 549 | T | C | | | 1 | 99 | synonymous_variant | LYS20 | YDL182W |
| gi\|696447274\|gb\|JRIS01000190.1\| | 555 | C | T | | | 1 | 99 | synonymous_variant | LYS20 | YDL182W |
| gi\|696447274\|gb\|JRIS01000190.1\| | 86616 | A | G | 1 | 99 | | | upstream_gene_variant | RDI1 | YDL135C |
| gi\|696447274\|gb\|JRIS01000190.1\| | 86628 | G | A | 1 | 99 | | | upstream_gene_variant | RDI1 | YDL135C |
| gi\|696447270\|gb\|JRIS01000191.1\| | 103480 | C | CT | 1 | 84 | | | upstream_gene_variant | RRP46 | YGR095C |
| gi\|696447265\|gb\|JRIS01000192.1\| | 137194 | GA | G | 1 | 99 | 0 | 99 | upstream_gene_variant | BLI1 | YKL061W |
| gi\|696447257\|gb\|JRIS01000194.1\| | 24773 | C | T | | | 1 | 91 | missense_variant | YNR065C | YNR065C |
| gi\|696447257\|gb\|JRIS01000194.1\| | 24778 | G | C | | | 1 | 55 | synonymous_variant | YNR065C | YNR065C |
| gi\|696447246\|gb\|JRIS01000197.1\| | 3999 | G | GA | 1 | 56 | 0 | 99 | upstream_gene_variant | SPC2 | YML055W |
| gi\|696447246\|gb\|JRIS01000197.1\| | 9995 | C | CT | | | 1 | 99 | upstream_gene_variant | CMP2 | YML057W |
| gi\|696447242\|gb\|JRIS01000198.1\| | 42814 | T | C | 1 | 99 | 0 | 99 | upstream_gene_variant | COX15 | YER141W |
| gi\|696447197\|gb\|JRIS01000210.1\| | 96 | T | C | 0 | 99 | 1 | 99 | upstream_gene_variant | YHL034C | YHL034C |
| gi\|696447197\|gb\|JRIS01000210.1\| | 103 | T | G | 0 | 99 | 1 | 99 | upstream_gene_variant | YHL034C | YHL034C |
| gi\|696447193\|gb\|JRIS01000211.1\| | 315 | G | A | | | 1 | 90 | upstream_gene_variant | YDL183C | YDL183C |
| gi\|696447193\|gb\|JRIS01000211.1\| | 325 | A | G | | | 1 | 90 | upstream_gene_variant | YDL183C | YDL183C |
| gi\|696447193\|gb\|JRIS01000211.1\| | 328 | T | C | | | 1 | 90 | upstream_gene_variant | YDL183C | YDL183C |
| gi\|696447193\|gb\|JRIS01000211.1\| | 329 | C | T | | | 1 | 90 | upstream_gene_variant | YDL183C | YDL183C |
| gi\|696447193\|gb\|JRIS01000211.1\| | 478 | G | A | | | 1 | 45 | upstream_gene_variant | YDL183C | YDL183C |
| gi\|696447193\|gb\|JRIS01000211.1\| | 484 | A | G | | | 1 | 45 | upstream_gene_variant | YDL183C | YDL183C |
| gi\|696447193\|gb\|JRIS01000211.1\| | 514 | G | A | | | 1 | 52 | upstream_gene_variant | YDL183C | YDL183C |
| gi\|696447174\|gb\|JRIS01000216.1\| | 68214 | G | T | | | 1 | 90 | upstream_gene_variant | ADH4 | YGL256W |
| gi\|696447174\|gb\|JRIS01000216.1\| | 68217 | T | C | | | 1 | 90 | upstream_gene_variant | ADH4 | YGL256W |
| gi\|696447169\|gb\|JRIS01000217.1\| | 81 | TCCACACCCACACCCACACA | T | 1 | 99 | . | . | intergenic_region | | |
| gi\|696447165\|gb\|JRIS01000218.1\| | 38619 | A | AACCGATGAATTCGAGCTCGTTTTCGACACTGGATGGCGGCGTTAGTATCGAATCGACAGCAGTATAGCGACCAGCATTCACATACGATTGACGCATGATATTACTTTCTGCGCACTTAACTTCGCATCTGGGCAGATGATGTCGAG | 0 | 45 | 1 | 99 | frameshift_variant&stop_gained | CLN3 | YAL040C |
| gi\|696447150\|gb\|JRIS01000222.1\| | 49905 | C | T | 1 | 45 | 0 | 13 | upstream_gene_variant | ECM29 | YHL030W |
| gi\|696447132\|gb\|JRIS01000227.1\| | 96460 | G | GA | | | 1 | 99 | upstream_gene_variant | TCP1 | YDR212W |
| gi\|696447076\|gb\|JRIS01000242.1\| | 416 | C | G | 0 | 99 | 1 | 99 | synonymous_variant | YHR214W | YHR214W |
| gi\|696447076\|gb\|JRIS01000242.1\| | 1034 | G | A | 1 | 99 | | | upstream_gene_variant | YHR214W | YHR214W |
| gi\|696447076\|gb\|JRIS01000242.1\| | 1340 | A | T | 1 | 99 | | | upstream_gene_variant | YHR214W | YHR214W |
| gi\|696446946\|gb\|JRIS01000279.1\| | 2083 | G | T | 1 | 99 | | | stop_gained | YJR157W | YJR157W |
| gi\|696446946\|gb\|JRIS01000279.1\| | 2145 | T | A | 1 | 99 | 0 | 99 | stop_gained | YJR157W | YJR157W |
| gi\|696446866\|gb\|JRIS01000302.1\| | 1153 | T | TA | | | 1 | 48 | downstream_gene_variant | HSP32 | YPL280W |
| gi\|696446851\|gb\|JRIS01000306.1\| | 123 | C | T | 1 | 99 | 0 | 99 | upstream_gene_variant | GEX1 | YCL073C |
| gi\|696446851\|gb\|JRIS01000306.1\| | 249 | A | C | 0 | 99 | 1 | 99 | upstream_gene_variant | GEX1 | YCL073C |
| gi\|696446851\|gb\|JRIS01000306.1\| | 1313 | A | C | | | 1 | 99 | synonymous_variant | GEX1 | YCL073C |
| gi\|696446851\|gb\|JRIS01000306.1\| | 1469 | T | C | | | 1 | 99 | synonymous_variant | GEX1 | YCL073C |
| gi\|696446851\|gb\|JRIS01000306.1\| | 1548 | G | A | 0 | 99 | 1 | 99 | missense_variant | GEX1 | YCL073C |
| gi\|696446851\|gb\|JRIS01000306.1\| | 1769 | A | G | 0 | 99 | 1 | 99 | synonymous_variant | GEX1 | YCL073C |
| gi\|696446851\|gb\|JRIS01000306.1\| | 2216 | T | C | 0 | 99 | 1 | 99 | synonymous_variant | GEX1 | YCL073C |
| gi\|696446851\|gb\|JRIS01000306.1\| | 2297 | C | T | 0 | 99 | 1 | 99 | synonymous_variant | GEX1 | YCL073C |
| gi\|696446851\|gb\|JRIS01000306.1\| | 2468 | G | C | 1 | 99 | 0 | 99 | synonymous_variant | GEX1 | YCL073C |
| gi\|696446851\|gb\|JRIS01000306.1\| | 2704 | T | C | 1 | 99 | 0 | 22 | missense_variant | GEX1 | YCL073C |
| gi\|696446851\|gb\|JRIS01000306.1\| | 2738 | C | CA | 1 | 99 | 0 | 99 | downstream_gene_variant | GEX1 | YCL073C |
| gi\|696446844\|gb\|JRIS01000308.1\| | 641 | G | T | 0 | 99 | 1 | 99 | upstream_gene_variant | DDI3 | YNL335W |
| gi\|696446844\|gb\|JRIS01000308.1\| | 674 | G | A | 0 | 99 | 1 | 99 | upstream_gene_variant | DDI3 | YNL335W |
| gi\|696446844\|gb\|JRIS01000308.1\| | 799 | T | G | 0 | 95 | 1 | 91 | upstream_gene_variant | DDI3 | YNL335W |
| gi\|696446844\|gb\|JRIS01000308.1\| | 918 | T | C | | | 1 | 99 | upstream_gene_variant | DDI3 | YNL335W |
| gi\|696446844\|gb\|JRIS01000308.1\| | 1218 | T | C | 0 | 99 | 1 | 99 | upstream_gene_variant | DDI3 | YNL335W |
| gi\|696446844\|gb\|JRIS01000308.1\| | 1615 | T | C | 1 | 99 | 0 | 99 | synonymous_variant | DDI3 | YNL335W |
| gi\|696446844\|gb\|JRIS01000308.1\| | 2349 | T | G | 0 | 35 | 1 | 99 | missense_variant | SNO2 | YNL334C |
| gi\|696446844\|gb\|JRIS01000308.1\| | 2699 | A | G | 1 | 99 | 0 | 99 | upstream_gene_variant | SNO2 | YNL334C |
| gi\|696446844\|gb\|JRIS01000308.1\| | 2903 | T | G | 0 | 99 | 1 | 99 | upstream_gene_variant | SNO2 | YNL334C |
| gi\|696446844\|gb\|JRIS01000308.1\| | 3006 | G | A | 1 | 99 | 0 | 99 | upstream_gene_variant | SNO2 | YNL334C |
| gi\|696446844\|gb\|JRIS01000308.1\| | 3831 | T | C | 1 | 99 | | | synonymous_variant | SNZ2 | YNL333W |
| gi\|696446844\|gb\|JRIS01000308.1\| | 3843 | G | A | 1 | 99 | | | synonymous_variant | SNZ2 | YNL333W |
| gi\|696446844\|gb\|JRIS01000308.1\| | 3850 | A | G | 1 | 99 | | | missense_variant | SNZ2 | YNL333W |
| gi\|696446798\|gb\|JRIS01000321.1\| | 1578 | T | G | 1 | 99 | 0 | 99 | missense_variant | PHO12 | YHR215W |
| gi\|696446798\|gb\|JRIS01000321.1\| | 2365 | T | C | 1 | 99 | 0 | 99 | synonymous_variant | PHO12 | YHR215W |
| gi\|696446798\|gb\|JRIS01000321.1\| | 3266 | A | T | 0 | 99 | 1 | 99 | upstream_gene_variant | PHO12 | YHR215W |
| gi\|696446794\|gb\|JRIS01000322.1\| | 403 | A | G | 1 | 99 | 0 | 99 | synonymous_variant | YOR390W | YOR390W |
| gi\|696446794\|gb\|JRIS01000322.1\| | 1589 | G | T | 0 | 99 | 1 | 99 | upstream_gene_variant | YOR390W | YOR390W |
| gi\|696446794\|gb\|JRIS01000322.1\| | 1599 | CT | C | 0 | 99 | 1 | 99 | upstream_gene_variant | YOR390W | YOR390W |
| gi\|696446794\|gb\|JRIS01000322.1\| | 1644 | T | G | 1 | 99 | | | upstream_gene_variant | YOR390W | YOR390W |
| gi\|696446794\|gb\|JRIS01000322.1\| | 1680 | T | A | 1 | 79 | | | upstream_gene_variant | YOR390W | YOR390W |
| gi\|696446729\|gb\|JRIS01000338.1\| | 477 | C | T | | | 1 | 99 | missense_variant | YBL109W | YBL109W |
| gi\|696446729\|gb\|JRIS01000338.1\| | 494 | CA | C | | | 1 | 45 | frameshift_variant | YBL109W | YBL109W |
| gi\|696446729\|gb\|JRIS01000338.1\| | 548 | T | C | | | 1 | 99 | missense_variant | YBL109W | YBL109W |
| gi\|696446729\|gb\|JRIS01000338.1\| | 561 | G | A | | | 1 | 99 | missense_variant | YBL109W | YBL109W |
| gi\|696446729\|gb\|JRIS01000338.1\| | 569 | T | C | | | 1 | 99 | missense_variant | YBL109W | YBL109W |
| gi\|696446726\|gb\|JRIS01000339.1\| | 510 | G | A | 1 | 99 | | | synonymous_variant | YMR326C | YMR326C |
| gi\|696446726\|gb\|JRIS01000339.1\| | 519 | A | C | 1 | 99 | | | missense_variant | YMR326C | YMR326C |
| gi\|696446726\|gb\|JRIS01000339.1\| | 547 | A | T | | | 1 | 99 | missense_variant | YMR326C | YMR326C |
| gi\|696446726\|gb\|JRIS01000339.1\| | 582 | T | TTGAAGTATGAGCACTACCATAACGTCATTAACGTAAAAGTTCCTTAATATTCCCATTTGCTTGAACGGATGCCACTTCAGAATATTTCGTACTTACACAGACTTCACATTAGAATATTATGTCATCCCATTGTTGTGACACTCTTTATTCACCGAGCAATAATACGGTAGTGGCTCACACACA | 0 | 99 | 1 | 90 | stop_gained&disruptive_inframe_insertion | YMR326C | YMR326C |
| gi\|696446714\|gb\|JRIS01000342.1\| | 305 | A | G | 0 | 40 | 1 | 44 | upstream_gene_variant | MAL12 | YGR292W |
| gi\|696446714\|gb\|JRIS01000342.1\| | 329 | G | T | 0 | 99 | 1 | 44 | upstream_gene_variant | MAL12 | YGR292W |
| gi\|696446714\|gb\|JRIS01000342.1\| | 401 | T | C | 0 | 99 | 1 | 99 | upstream_gene_variant | MAL12 | YGR292W |
| gi\|696446714\|gb\|JRIS01000342.1\| | 406 | C | T | 0 | 99 | 1 | 99 | upstream_gene_variant | MAL12 | YGR292W |
| gi\|696446714\|gb\|JRIS01000342.1\| | 464 | A | T | 1 | 99 | 0 | 67 | upstream_gene_variant | MAL12 | YGR292W |
| gi\|696446714\|gb\|JRIS01000342.1\| | 720 | T | C | 1 | 83 | 0 | 99 | synonymous_variant | MAL12 | YGR292W |
| gi\|696446714\|gb\|JRIS01000342.1\| | 2238 | G | A | 1 | 99 | 0 | 99 | downstream_gene_variant | MAL12 | YGR292W |
| gi\|696446714\|gb\|JRIS01000342.1\| | 2272 | C | G | 1 | 99 | 0 | 99 | downstream_gene_variant | MAL12 | YGR292W |
| gi\|696446714\|gb\|JRIS01000342.1\| | 2327 | G | A | 1 | 99 | 0 | 99 | downstream_gene_variant | MAL12 | YGR292W |
| gi\|696446714\|gb\|JRIS01000342.1\| | 2388 | C | T | 1 | 99 | 0 | 99 | downstream_gene_variant | MAL12 | YGR292W |
| gi\|696446714\|gb\|JRIS01000342.1\| | 2446 | A | G | 0 | 99 | 1 | 99 | downstream_gene_variant | MAL12 | YGR292W |
| gi\|696446714\|gb\|JRIS01000342.1\| | 2623 | T | C | 1 | 99 | 0 | 99 | downstream_gene_variant | MAL12 | YGR292W |
| gi\|696446714\|gb\|JRIS01000342.1\| | 2626 | G | A | 1 | 99 | 0 | 99 | downstream_gene_variant | MAL12 | YGR292W |
| gi\|696446714\|gb\|JRIS01000342.1\| | 2711 | G | A | 0 | 99 | 1 | 99 | downstream_gene_variant | MAL12 | YGR292W |
| gi\|696446714\|gb\|JRIS01000342.1\| | 2735 | T | C | 0 | 99 | 1 | 99 | downstream_gene_variant | MAL12 | YGR292W |
| gi\|696446714\|gb\|JRIS01000342.1\| | 2878 | C | T | 1 | 99 | 0 | 52 | downstream_gene_variant | MAL12 | YGR292W |
| gi\|696446714\|gb\|JRIS01000342.1\| | 2898 | G | T | 1 | 99 | | | downstream_gene_variant | MAL12 | YGR292W |
| gi\|696446714\|gb\|JRIS01000342.1\| | 2928 | A | T | 1 | 99 | 0 | 99 | downstream_gene_variant | MAL12 | YGR292W |
| gi\|696446714\|gb\|JRIS01000342.1\| | 2941 | G | A | 1 | 99 | 0 | 99 | downstream_gene_variant | MAL12 | YGR292W |
| gi\|696446714\|gb\|JRIS01000342.1\| | 2949 | T | C | 1 | 99 | 0 | 99 | downstream_gene_variant | MAL12 | YGR292W |
| gi\|696446714\|gb\|JRIS01000342.1\| | 2977 | A | C | 1 | 99 | 0 | 99 | downstream_gene_variant | MAL12 | YGR292W |
| gi\|696446714\|gb\|JRIS01000342.1\| | 2988 | C | T | 1 | 99 | 0 | 99 | downstream_gene_variant | MAL12 | YGR292W |
| gi\|696446714\|gb\|JRIS01000342.1\| | 2999 | G | T | 1 | 99 | 0 | 99 | downstream_gene_variant | MAL12 | YGR292W |
| gi\|696446714\|gb\|JRIS01000342.1\| | 3000 | A | G | 1 | 99 | 0 | 99 | downstream_gene_variant | MAL12 | YGR292W |
| gi\|696446714\|gb\|JRIS01000342.1\| | 3016 | C | G | 1 | 99 | 0 | 99 | downstream_gene_variant | MAL12 | YGR292W |
| gi\|696446714\|gb\|JRIS01000342.1\| | 3033 | A | G | 1 | 99 | 0 | 99 | downstream_gene_variant | MAL12 | YGR292W |
| gi\|696446714\|gb\|JRIS01000342.1\| | 3043 | G | T | 1 | 99 | 0 | 99 | downstream_gene_variant | MAL12 | YGR292W |
| gi\|696446714\|gb\|JRIS01000342.1\| | 3056 | C | T | 1 | 99 | 0 | 99 | downstream_gene_variant | MAL12 | YGR292W |
| gi\|696446714\|gb\|JRIS01000342.1\| | 3062 | G | A | 1 | 99 | 0 | 99 | downstream_gene_variant | MAL12 | YGR292W |

**Supplementary Table 9: Samtools variant analysis of contigs against the S288C reference genome.** Columns 1 and 2 detail the position of the variant. Columns 3 and 4 note the reference allele (Genotype 0) and the alternative allele (Genotype 1). Columns 5 and 6 denote the called genotype for strains 1691 and 1693 respectively. Columns 7-9 report the location of the variant and its effect. For the sake of brevity, those variants which are shared between 1691 and 1693 are not shown, and for cases where there were five or more discordant variants called for a single gene, those variants are not listed individually (this was the case for many genes such as FLO9 that appear to contain a structural variation). Those variants that were also called when analyzing the trimmed reads (as opposed to the assembled contigs) are shaded. Clear discordand SNPs that could be confirmed by aligning the reads to IGV are shaded dark grey. Each of these was also called when analyzing trimmed reads (Supplemental Tables 5 & 6).

| CHROM | POS | Genotype 0 | Genotype 1 | 1691 | 1693 | Annotation | Gene | Gene ID |
|---|---|---|---|---|---|---|---|---|
| I | 12690 | AGAAGTAATTTCCTGACTTGTTGTTGCACTGGTAA<br>CAGGTGGTAATGATGAAGTAATTTCCTGACTTGT<br>TGTTGTACTGGTAACAGGTGGTAATGATGAAG | AGAAGTAATTTCCTGACTTGTTGTTGCACTGGTAACAGGTGGTAATGATGAAGTAATTTCCTGACTTGTTGTTGTACTGGTAACAGGTGGTAATGATGAAGTAATTTC<br>CTGACTTGTTGTTGTACTGGTAACAGGTGGTAATGAAGAAGTAATTTCCTGACTTGTTGTTGCACTGGTAACAGGTGGTAATGATGAAGTAATTTCCTGACTTGTTGT<br>TGTACTGGTAACAGGTGGTAATGAAGAAGTAATTTCCTGACTTGTTGTTGTACTGGTAACAGGTGGTAATGATGAAGTAATTTCCTGACTTGCTGTCGCACTGGTAA<br>CAGGTGGTAATGAAGAAGTAATTTCCTGACTTGTTGTTGCACTGGTAACAGGTGGTAATGATGAAGTAATTTCCTGACTTGTTGTTGCACTGGTAACAGGTGGTAAT<br>GATGAAGTAATTTCCTGACTTGTTGTTGTACTGGTAACAGGTGGTAATGATGAAG | 0 | 1 | upstream_gene_variant | SEO1 | YAL067C |
| I | | | 53 SNPs between 26221 and 26799 | - | - | missense_variant | FLO9 | YAL063C |
| I | 141030 | T | A | 1 | 0 | missense_variant | YAL004W | YAL004W |
| I | | | 59 SNPs between 203639 and 212211 | - | - | synonymous_variant | FLO1 | YAR050W |
| I | 213637 | A | T | 1 | 0 | upstream_gene_variant | YAR061W | YAR061W |
| I | 214057 | C | A | 1 | 0 | upstream_gene_variant | YAR061W | YAR061W |
| I | 214061 | C | A | 1 | 0 | upstream_gene_variant | YAR061W | YAR061W |
| I | 219252 | T | C | 1 | 0 | upstream_gene_variant | YAR060C | YAR060C |
| I | 220292 | G | C | 1 | 0 | missense_variant | YAR064W | YAR064W |
| I | 220382 | C | A | 1 | 0 | missense_variant | YAR064W | YAR064W |
| I | 220499 | T | A | 1 | 0 | upstream_gene_variant | YAR060C | YAR060C |
| I | 220805 | C | T | 1 | 0 | upstream_gene_variant | YAR060C | YAR060C |
| I | 222254 | T | C | 1 | 0 | upstream_gene_variant | YAR060C | YAR060C |
| II | 68282 | GTTATTATTATTATTATTATTATTATTATTATTATT<br>ATTATTATTATTATTATTATTATTATTATT | GTTATTATTATTATTATTATTATTATTATTATTATTATTATTATTATTATTATTATTATTATTATTATTATTATTATTATTATTATTATTATTATT | 0 | 1 | conservative_inframe_insertion | CDC27 | YBL084C |
| II | 221505 | C | T | 0 | 1 | missense_variant | YBL005W-A | YBL005W-A |
| II | 224777 | G | A | 1 | 0 | synonymous_variant | YBL005W-B | YBL005W-B |
| II | 466955 | A | C | 1 | 0 | upstream_gene_variant | CYC8 | YBR112C |
| III | | | 7 SNPs between 206 and 586 | - | - | upstream_gene_variant | YCL076W | YCL076W |
| III | 200119 | G | T | 1 | 0 | synonymous_variant | MATALPHA2 | YCR039C |
| III | 200145 | G | T | 1 | 0 | missense_variant | MATALPHA2 | YCR039C |
| III | 200158 | T | C | 1 | 0 | synonymous_variant | MATALPHA2 | YCR039C |
| III | 200166 | G | A | 1 | 0 | missense_variant | MATALPHA2 | YCR039C |
| III | | | 36 SNPs between 200188 and 200434 | - | - | upstream_gene_variant | PHO87 | YCR037C |
| III | | | 42 SNPs between 200444 and 200850 | - | - | start_lost | MATALPHA1 | YCR040W |
| III | | | 55 SNPs between 268850 and 759248 | - | - | missense_variant | FIG2 | YCR089W |
| IV | 827861 | G | T | 0 | 1 | missense_variant | CDC1 | YDR182W |
| IV | 892698 | GTTTTTTTTTTTTTTTTTTTTTTTTTTTTTTTTTT | GTTTTTTTTTTTTTTTTTTTTTTTTTTTTTTTTTTTTTTTTTTTTTTTTTTTTTTTTTTTTTT | 1 | 0 | upstream_gene_variant | AHA1 | YDR214W |
| IV | | | 6 SNPs between 1525363 and 1525431 | - | - | missense_variant | YDR544C | YDR544C |
| V | 2084 | T | C | 1 | 0 | missense_variant | YEL077C | YEL077C |
| V | 2116 | T | C | 1 | 0 | missense_variant | YEL077C | YEL077C |
| V | 2146 | T | C | 1 | 0 | missense_variant | YEL077C | YEL077C |
| V | 20824 | G | A | 0 | 1 | synonymous_variant | DSF1 | YEL070W |
| V | 205872 | G | A | 1 | 0 | missense_variant | GCD11 | YER025W |
| V | 362749 | A | T | 0 | 1 | upstream_gene_variant | PRS2 | YER099C |
| VI | 4716 | CCCACACCCACACACCACACCCACACCACACCCACAC<br>ACCACACCCACACCACACC | CCCACACCCACACACCACACCCACACCACACC | 0 | 1 | upstream_gene_variant | YFL066C | YFL066C |
| VI | 9922 | T | C | 1 | 0 | synonymous_variant | DDI2 | YFL061W |
| VI | 11006 | A | G | 1 | 0 | upstream_gene_variant | SNO3 | YFL060C |
| VI | 11211 | G | T | 1 | 0 | upstream_gene_variant | SNO3 | YFL060C |
| VI | 11314 | G | A | 1 | 0 | upstream_gene_variant | SNO3 | YFL060C |
| VII | 10981 | T | A | 1 | 0 | upstream_gene_variant | PAU11 | YGL261C |
| VII | 364650 | C | G | 1 | 0 | synonymous_variant | RPL7A | YGL076C |
| VII | 364785 | G | A | 0 | 1 | synonymous_variant | RPL7A | YGL076C |
| VII | 535885 | A | G | 0 | 1 | upstream_gene_variant | THG1 | YGR024C |
| VIII | | | 5 SNPs between 4844 and 6105 | - | - | upstream_gene_variant | YHL050C | YHL050C |
| VIII | 289318 | A | G | 0 | 1 | synonymous_variant | AHT1 | YHR093W |
| VIII | 340058 | TTATATATATATATATATATATATATATATA | TTATATATATATATATATATATATATATATA | 1 | 0 | upstream_gene_variant | YHR112C | YHR112C |
| VIII | 499958 | C | T | 0 | 1 | missense_variant | PPX1 | YHR201C |
| VIII | 511255 | C | T | 0 | 1 | synonymous_variant | SCH9 | YHR205W |
| VIII | 514829 | CTATATATATATATATATATATATATATA | CTATATATATATATATATATATATATATATATA | 1 | 0 | upstream_gene_variant | BAT1 | YHR208W |
| VIII | | | 18 SNPs between 525628 and 529805 | - | - | synonymous_variant | FLO5 | YHR211W |
| VIII | 549816 | T | G | 1 | 0 | upstream_gene_variant | YHR214C-B | YHR214C-B |
| VIII | 551421 | G | A | 1 | 0 | missense_variant | YHR214C-E | YHR214C-E |
| VIII | 551423 | A | C | 1 | 0 | missense_variant | YHR214C-E | YHR214C-E |
| VIII | 551439 | A | T | 1 | 0 | missense_variant | YHR214C-E | YHR214C-E |
| VIII | 551468 | A | T | 1 | 0 | missense_variant | YHR214C-E | YHR214C-E |
| VIII | 552371 | G | A | 0 | 1 | synonymous_variant | PHO12 | YHR215W |
| IX | | | 6 SNPs between 206802 and 210095 | - | - | missense_variant | YIL082W-A | YIL082W-A |
| IX | 210176 | C | T | 1 | 0 | upstream_gene_variant | tI(AAU)I2 | tI(AAU)I2 |
| IX | 387143 | C | A | 0 | 1 | upstream_gene_variant | MET28 | YIR017C |
| IX | | | 21 SNPs between 391339 and 392245 | - | - | synonymous_variant | FLO11 | YIR019C |
| IX | | | 5 SNPs between 439814 and 439877 | - | - | upstream_gene_variant | YIR042C | YIR042C |
| X | 639694 | C | A | 1 | 0 | upstream_gene_variant | YJR111C | YJR111C |
| X | 639709 | C | A | 1 | 0 | upstream_gene_variant | YJR111C | YJR111C |
| X | 652909 | G | T | 0 | 1 | missense_variant | YJR124C | YJR124C |
| X | 652916 | A | T | 0 | 1 | synonymous_variant | YJR124C | YJR124C |
| X | | | 9 SNPs between 715095 and 715141 | - | - | missense_variant | DAN4 | YJR151C |
| X | 730773 | A | T | 1 | 0 | stop_gained | YJR157W | YJR157W |
| X | 730835 | C | A | 1 | 0 | stop_gained | YJR157W | YJR157W |
| X | 731773 | C | T | 1 | 0 | upstream_gene_variant | THI11 | YJR156C |
| X | 733265 | G | A | 1 | 0 | missense_variant | HXT16 | YJR158W |
| X | 733998 | C | T | 1 | 0 | missense_variant | HXT16 | YJR158W |
| X | 734783 | G | A | 1 | 0 | upstream_gene_variant | SOR1 | YJR159W |
| X | | | 6 SNPs between 740157 and 742235 | - | - | upstream_gene_variant | MPH3 | YJR160C |
| X | 743001 | C | G | 1 | 0 | missense_variant | COS5 | YJR161C |
| X | 743217 | T | G | 1 | 0 | synonymous_variant | COS5 | YJR161C |
| X | 745680 | G | T | 1 | 0 | upstream_gene_variant | COS5 | YJR161C |
| XI | 28 | A | C | 1 | 0 | upstream_gene_variant | YKL225W | YKL225W |
| XI | 59567 | T | G | 0 | 1 | missense_variant | TOR2 | YKL203C |
| XI | 164844 | CTATATATATATATATATATATATATATATATATAT | CTATATATATATATATATATATATATATATATATATAT | 0 | 1 | upstream_gene_variant | RSM22 | YKL155C |
| XI | 541138 | G | T | 1 | 0 | missense_variant | DYN1 | YKR054C |
| XII | 12277 | C | T | 1 | 0 | upstream_gene_variant | YLL066C | YLL066C |
| XII | 13020 | C | T | 1 | 0 | upstream_gene_variant | YLL066C | YLL066C |
| XII | | | 6 SNPs between 13122 and 13170 | - | - | missense_variant | PAU18 | YLL064C |
| XII | 96700 | A | G | . | 1 | synonymous_variant | SSA2 | YLL024C |
| XII | 220717 | G | A | 1 | 0 | upstream_gene_variant | YLR035C-A | YLR035C-A |
| XII | 460291 | A | C | 1 | 0 | upstream_gene_variant | RDN37-1 | RDN37-1 |
| XII | 474417 | ACC | ACCC | 1 | 0 | upstream_gene_variant | ASP3-1 | YLR155C |
| XII | 734703 | T | A | 1 | 0 | upstream_gene_variant | YLR302C | YLR302C |
| XII | | | 9 SNPs between 1059473 and 1061528 | - | - | upstream_gene_variant | NBP1 | YLR457C |
| XIII | 6154 | A | T | 0 | 1 | upstream_gene_variant | YML133C | YML133C |
| XIII | 6933 | C | T | 1 | 0 | upstream_gene_variant | YML133C | YML133C |
| XIII | 6941 | G | A | 1 | 0 | upstream_gene_variant | YML133C | YML133C |
| XIII | 812314 | A | C | 0 | 1 | missense_variant | ZDS1 | YMR273C |
| XIII | 924337 | T | G | 1 | 0 | upstream_gene_variant | YMR324C | YMR324C |
| XIII | 924341 | G | T | 1 | 0 | upstream_gene_variant | YMR324C | YMR324C |
| XIV | 189585 | C | T | 1 | 0 | missense_variant | SLA2 | YNL243W |
| XIV | 415047 | CTTTTTT | CTTTTTTTTTTTTTTTTTTTTTCGTTTTTTATTTTTTTTTTTTTTTTTTTTTATTTATTTTTTTTTTTTTTTTTTATTTTTTTTTTTTTTTTTTTTTTTTTTTTTTTTTT | 0 | 1 | upstream_gene_variant | YNL115C | YNL115C |
| XIV | 415054 | C | T | 0 | 1 | upstream_gene_variant | YNL115C | YNL115C |
| XIV | 415088 | C | T | 1 | 0 | upstream_gene_variant | YNL115C | YNL115C |
| XIV | 415096 | C | T | 1 | 0 | upstream_gene_variant | YNL115C | YNL115C |
| XIV | 601677 | T | C | 1 | 0 | missense_variant | YNL018C | YNL018C |
| XIV | 601742 | G | A | 1 | 0 | synonymous_variant | YNL018C | YNL018C |
| XV | 78 | CA | C | 0 | 1 | upstream_gene_variant | YOL166W-A | YOL166W-A |
| XV | 97 | A | C | 1 | 0 | upstream_gene_variant | YOL166W-A | YOL166W-A |
| XV | 98 | C | A | 1 | 0 | upstream_gene_variant | YOL166W-A | YOL166W-A |
| XV | 108 | A | C | 1 | 0 | upstream_gene_variant | YOL166W-A | YOL166W-A |
| XV | 2294 | G | C | 0 | 1 | upstream_gene_variant | YOL166C | YOL166C |
| XV | 2345 | G | A | 0 | 1 | upstream_gene_variant | YOL166C | YOL166C |
| XV | 2398 | T | C | 0 | 1 | upstream_gene_variant | YOL166C | YOL166C |
| XV | | | 9 SNPsbetween 970965 and 971069 | - | - | synonymous_variant | YOR343W-A | YOR343W-A |
| XVI | 174252 | T | C | 1 | 0 | synonymous_variant | RPL7B | YPL198W |
| XVI | 174387 | C | G | 0 | 1 | missense_variant | YPL197C | YPL197C |
| XVI | 560663 | T | A | 1 | 0 | upstream_gene_variant | YPR002C-A | YPR002C-A |
| XVI | 667520 | T | G | 0 | 1 | upstream_gene_variant | ATG11 | YPR049C |
| XVI | 769498 | T | C | 1 | 0 | upstream_gene_variant | tK(UUU)P | tK(UUU)P |
| XVI | 769532 | G | A | 1 | 0 | upstream_gene_variant | tK(UUU)P | tK(UUU)P |
| XVI | 769580 | A | G | 1 | 0 | upstream_gene_variant | tK(UUU)P | tK(UUU)P |
| XVI | 838970 | T | A | 1 | 0 | upstream_gene_variant | TPO3 | YPR156C |
| XVI | | | 13 SNPs between 853008-853335 | - | - | synonymous_variant | YPR158C-D | YPR158C-D |

**Supplemetary Table 10: Breakdancer CNV Analysis.** The first six columns record the position(s) of the event and the open reading frame(s) detected. If the detected event is not a translocation, columns 1-3 are the same as coumns 4-6. Column 7 and 8 detail the type and size of the event: DEL (deletions), INS (insertion), INV (inversion), ITX (intra-chromosomal translocation), CTX (inter-chromosomal translocation). Columns 8-10 list the confidence score and the number of supporting read pairs for each strain. Events that are located in repetative regions such as telomeres, transposons, and ribosomal RNA genes are shaded light grey. Events which could be visually confirmed by alighning the reads to the S288C

| Chr1 | Pos1 | ORF | Chr2 | Pos2 | ORF | Type | Size | Score | num_Reads_1691 | num_Reads_1693 |
|---|---|---|---|---|---|---|---|---|---|---|
| I | 12474 | intergenic (downstream of YAL064W-B) | I | 12684 | intergenic (downstream of YAL064W-B) | ITX | -287 | 99 | 23 | 18 |
| I | 26074 | YAL063C (FLO9) | I | 26510 | YAL063C (FLO9) | ITX | -336 | 71 | 3 | 1 |
| I | 26891 | YAL063C (FLO9) | I | 27051 | YAL063C (FLO9) | ITX | -304 | 99 | 3 | 7 |
| II | 469882 | YBR115C (LYS2) | II | 474220 | YBR115C (LYS2) | DEL | 4506 | 99 | 25 | 33 |
| II | 469882 | YBR115C (LYS2) | II | 474469 | YBR115C (LYS2) | DEL | 4740 | 70 | 2 | 1 |
| II | 469882 | YBR115C (LYS2) | II | 474602 | YBR115C (LYS2) | DEL | 4862 | 71 | 1 | 2 |
| III | 17414 | YCL063W | III | 17458 | YCL063W | INS | -471 | 99 | 1 | 2 |
| III | 84509 | YCL019W (LEU2) | III | 92480 | YCL019W (LEU2) | DEL | 7917 | 99 | 1 | 4 |
| III | 84939 | YCL019W (LEU2) | III | 92480 | YCL019W (LEU2) | DEL | 7692 | 99 | 24 | 51 |
| III | 268980 | YCR089W (known duplication) | III | 269334 | YCR089W (known duplication) | ITX | -338 | 94 | 2 | 2 |
| III | 200940 | YCR039C/YCR040W (ALPHA2/ALPHA1 | III | 294607 | YCR098C (GIT1) | ITX | 92452 | 99 | 23 | 26 |
| IV | 758478 | YDR150W | IV | 758586 | YDR150W | ITX | -334 | 73 | 3 | 1 |
| IV | 1525372 | telomere | IV | 1525452 | YDR544C | ITX | -315 | 99 | 4 | 6 |
| IX | 25362 | telomere | IX | 25535 | YIL169C | ITX | -293 | 99 | 3 | 4 |
| Mito | 6913 | | Mito | 47815 | | INV | 23166 | 99 | 13 | 23 |
| Mito | 11215 | | Mito | 47815 | | INV | 1890 | 99 | 7 | 16 |
| Mito | 13618 | | Mito | 47815 | | INV | 23840 | 93 | 3 | 9 |
| Mito | 47815 | | Mito | 73182 | | INV | 6251 | 99 | 17 | 23 |
| Mito | 47815 | | Mito | 80215 | | INV | 6079 | 99 | 7 | 10 |
| Mito | 47815 | | Mito | 85980 | | INV | 153 | 99 | 5 | 9 |
| Mito | 1201 | | Mito | 83430 | | INV | 81133 | 93 | 4 | 4 |
| Mito | 1 | | Mito | 85980 | | ITX | 84733 | 99 | 180 | 350 |
| Mito | 6913 | | Mito | 16970 | | INV | 1105 | 69 | 2 | 6 |
| Mito | 6913 | | Mito | 31030 | | INV | 25843 | 99 | 8 | 9 |
| Mito | 6913 | | Mito | 36806 | | INV | 28862 | 71 | 4 | 6 |
| Mito | 6913 | | Mito | 52951 | | DEL | 51683 | 73 | 4 | 9 |
| Mito | 6913 | | Mito | 64262 | | DEL | 65288 | 99 | 12 | 11 |
| Mito | 6913 | | Mito | 83430 | | INV | 77862 | 99 | 21 | 17 |
| Mito | 6913 | | Mito | 85980 | | INV | 80059 | 99 | 8 | 5 |
| Mito | 11215 | | Mito | 80027 | | DEL | 71235 | 99 | 7 | 6 |
| Mito | 11990 | | Mito | 80027 | | DEL | 70487 | 60 | 2 | 4 |
| Mito | 13618 | | Mito | 31030 | | INV | 17056 | 99 | 2 | 7 |
| Mito | 13618 | | Mito | 83430 | | INV | 69308 | 80 | 3 | 4 |
| Mito | 31030 | | Mito | 60441 | | INV | 24625 | 99 | 9 | 9 |
| Mito | 29316 | | Mito | 83430 | | ITX | 51070 | 58 | 13 | 12 |
| Mito | 36806 | | Mito | 53145 | | INV | 15660 | 99 | 8 | 7 |
| Mito | 36806 | | Mito | 60441 | | INV | 22111 | 99 | 14 | 18 |
| Mito | 53145 | | Mito | 73182 | | INV | 19234 | 53 | 4 | 5 |
| Mito | 53145 | | Mito | 83430 | | INV | 29092 | 74 | 4 | 5 |
| Mito | 60441 | | Mito | 73182 | | INV | 11493 | 97 | 6 | 9 |
| Mito | 60441 | | Mito | 83430 | | INV | 26832 | 99 | 16 | 14 |
| Mito | 60441 | | Mito | 85980 | | INV | 27581 | 68 | 4 | 5 |
| V | 116108 | YEL021W (URA3) | V | 117045 | YEL021W (URA3) | DEL | 1135 | 99 | 30 | 33 |
| VII | 356826 | YGL082W (known duplication) | VII | 356858 | YGL082W (known duplication) | INS | -490 | 99 | 54 | 111 |
| I | 1 | telomere | VIII | 562821 | intergenic (downsteram of YHR219W) | ITX | -427 | 51 | 4 | 4 |
| X | 508773 | YJR040W | X | 508807 | YJR040W | INS | -466 | 99 | 1 | 1 |
| X | 713026 | YJR151C (DAN4) | X | 713371 | YJR151C (DAN4) | ITX | -305 | 99 | 3 | 2 |
| VIII | 556799 | telomere | XII | 363 | telomere | ITX | -355 | 99 | 7 | 4 |
| II | 165870 | intergenic (upstream of YBL029W) | XII | 460787 | intergenic (upstream of RDN37-2) | ITX | -346 | 99 | 19 | 23 |
| XII | 732412 | YLR303W (MET15) | XII | 734686 | YLR303W (MET15) | DEL | 2456 | 99 | 31 | 0 |
| XIII | 908139 | telomere | XIII | 908743 | telomere | ITX | -378 | 99 | 17 | 12 |
| XIV | 704446 | YNR044W (known duplication) | XIV | 704473 | YNR044W (known duplication) | ITX | -282 | 67 | 1 | 1 |
| IX | 23044 | telomere | XV | 28722 | YOL019W (known duplication) | CTX | -510 | 99 | 6 | 15 |
| IX | 23880 | telomere | XV | 28722 | YOL019W (known duplication) | CTX | -502 | 99 | 17 | 16 |

**Supplemetary Table 11: CNV.kit Analysis.** Only those genome regions with a copy number different than 1 are shown. Events that are located in repetative regions such as telomeres, transposons, and ribosomal RNA genes are shaded light grey. Events which could be visually confirmed by alighning the reads to the S288C reference genome in IGV are shaded dark grey.

| 1691 | | | | |
|---|---|---|---|---|
| chromosome | start | end | ORF | copy numer |
| X | 472681 | 483859 | YJR026W, YJR027W, YJR029W (transposons) | 2 |
| Mito | 0 | 85779 | mitochondria | 22 |
| VII | 535087 | 543074 | YGR027W-A (transposon), YGR027W-B (transposon), tE(UUC)G3, YGR028W | 2 |
| XII | 450438 | 469605 | RDN25-1, RDN37-1, RDN18-1, RDN5-1, RDN37-2, RDN18-2, RDN5-2 | 94 |
| XV | 116638 | 123029 | YOL104C, YOL103W-B (transposon) | 2 |
| XVI | 844146 | 856936 | YPR158W-A, YPR158W-B, YPR158C-D, YPR158C-C (transposons) | 2 |

| 1693 | | | | |
|---|---|---|---|---|
| chromosome | start | end | ORF | copy number |
| II | 469902 | 474122 | YBR115C (lys2) | 0 |
| IV | 528009 | 537866 | YDR38C (ENA5), YDR39C (ENA2), YDR40C (ENA1) | 2 |
| IV | 987026 | 992658 | YDR261C-D, YDR261C-C (transposons) | 2 |
| Mito | 0 | 85779 | mitochondria | 29 |
| VII | 534912 | 541950 | YGR027W-A (transposon), YGR027W-B (transposon), tE(UUC)G3, YGR028W | 2 |
| VIII | 212397 | 216617 | RUF5-1, YHR054C (known partial duplication), RUF5-2, YHR056C (known duplicat | 15 |
| XII | 451820 | 468711 | RDN25-1, RDN37-1, RDN18-1, RDN5-1, RDN37-2, RDN18-2, RDN5-2 | 94 |
| XII | 1068324 | 1078177 | telomere | 2 |

**Supplemetary Table 12: CNVnator Analysis, bin size = 20.** Events that are located in repetative regions such as telomeres, transposons, and ribosomal RNA genes are shaded light grey. Events which could be visually confirmed by alighning the reads to the S288C reference genome in IGV are shaded dark grey.

| 1691 | | | |
|---|---|---|---|
| CNV type | coordinates | context/ORF | CNV_size |
| duplication | I:1-23720 | telomere | 23720 |
| duplication | I:23761-30780 | YAL063C (flo9) | 7020 |
| duplication | I:48301-50480 | YAL051W (known duplicate) | 2180 |
| duplication | I:63561-65760 | YAL04W1 | 2200 |
| deletion | I:65761-67520 | YAL040C (cln3) | 1760 |
| duplication | I:159741-189920 | YAR009C (transposon) | 30180 |
| deletion | I:189921-192100 | YAR035W | 2180 |
| duplication | I:196221-203860 | telomere | 7640 |
| deletion | I:204601-205760 | telomere | 1160 |
| duplication | I:206261-210840 | telomere | 4580 |
| duplication | I:214221-230220 | telomere | 16000 |
| duplication | II:1-30640 | telomere | 30640 |
| deletion | II:30641-31860 | YBL100W-B (transposon) | 1220 |
| duplication | II:31861-54220 | YBL100W-B (transposon) | 22360 |
| duplication | II:108181-110580 | YBL060W, YBL059C-A | 2400 |
| duplication | II:222841-226960 | YBL005W-B (transposon) | 4120 |
| deletion | II:261281-261520 | YBR012W-B (transposon) | 240 |
| duplication | II:261541-263020 | YBR012W-B (transposon) | 1480 |
| deletion | II:264141-264400 | YBR012W-B (transposon) | 260 |
| duplication | II:295201-296940 | YBR028C, YBR029C | 1740 |
| deletion | II:455401-457500 | YBR108W | 2100 |
| deletion | II:469701-474240 | YBR115C (lys2) | 4540 |
| duplication | II:642341-647840 | YBR209W, tE(UUC)B, YBR210W (known duplicate) , YBR211C | 5500 |
| duplication | II:653161-657600 | YBR215W, YBR216C (known duplicate) | 4440 |
| duplication | II:751621-813200 | telomere | 61580 |
| duplication | III:1-12140 | telomere | 12140 |
| duplication | III:13921-47320 | YCL067C (HMLALPHA2), YCL066W (HLMAPHA1) | 33400 |
| duplication | III:80141-84780 | YCL024W (known duplicate), YCL022C, tE(UUC)C, YCL021W-A | 4640 |
| deletion | III:84781-88320 | YCL019W (leu2) | 3540 |
| deletion | III:90661-92480 | YCL019W (leu2) | 1820 |
| deletion | III:148601-151720 | telomere | 3120 |
| duplication | III:168381-170600 | telomere | 2220 |
| deletion | III:250941-252440 | YCR077C | 1500 |
| duplication | III:255001-316580 | telomere | 61580 |
| duplication | IV:1-31360 | telomere | 31360 |
| duplication | IV:46521-48460 | YDL227C | 1940 |
| deletion | IV:352881-355380 | YDL056W (mbp1) | 2500 |
| duplication | IV:514301-515700 | YDR034C-D (transposon) | 1400 |
| duplication | IV:528741-531500 | YDR038C (ENA5) | 2760 |
| duplication | IV:531881-537560 | YDR039C (ENA2) | 5680 |
| duplication | IV:645761-651740 | YDR098C-B (transposon) | 5980 |
| duplication | IV:875561-883880 | YDR210C-D (transposon) | 8320 |
| duplication | IV:985221-992800 | YDR261C-D (transposon) | 7580 |
| duplication | IV:1095961-1102200 | YDR316W-B (transposon) | 6240 |
| duplication | IV:1206821-1212420 | YDR36W-B (transposon) | 5600 |
| duplication | IV:1480681-1531940 | telomere | 51260 |
| duplication | IX:1-2220 | telomere | 2220 |
| duplication | IX:3861-18480 | telomere | 14620 |
| deletion | IX:18741-18980 | telomere | 240 |
| duplication | IX:18981-22000 | telomere | 3020 |
| duplication | IX:23461-36820 | YIL169C, YIL168W, YIL166C, YIL165C | 13360 |
| duplication | IX:42761-46520 | YIL159W, YIL158W (known duplicate) | 3760 |
| duplication | IX:270881-274140 | YIL045W (known duplicate), YIL044C | 3260 |
| duplication | IX:391661-439800 | telomere | 48140 |
| duplication | Mito:1-8980 | mitochondrial | 8980 |
| duplication | Mito:9021-47660 | mitochondrial | 38640 |
| duplication | Mito:47881-56900 | mitochondrial | 9020 |
| duplication | Mito:57021-66700 | mitochondrial | 9680 |
| duplication | Mito:66741-70560 | mitochondrial | 3820 |
| duplication | Mito:70661-72860 | mitochondrial | 2200 |
| duplication | Mito:72961-85780 | mitochondrial | 12820 |
| duplication | V:1-40040 | telomere | 40040 |
| duplication | V:45001-65300 | YEL056W, YEL055C, snR80, YEL053C, YEL052W, YEL050C, snR67, YEL049W | 20300 |
| deletion | V:115921-117060 | YEL021W (URA3) | 1140 |
| duplication | V:443761-449040 | YER138C (transposon) | 5280 |
| deletion | V:473021-476100 | YER152C, YER152W-A, YER153C, YER154W | 3080 |
| duplication | V:492601-498740 | YER160C (transposon) | 6140 |
| duplication | V:523081-544100 | YER169W (known duplication), YER170W, YER171W, YER172C, YER172C-A, YER174C (known duplication), YER175W-A, YER176W | 21020 |
| duplication | V:545621-576880 | telomere | 31260 |
| duplication | VI:1-48120 | telomere | 48120 |
| duplication | VI:84961-88940 | YFL025C, YFL024C | 3980 |
| duplication | VI:109881-112600 | YFL012W-A, YFL012W, YFL011W | 2720 |
| duplication | VI:137041-138260 | YFL003C, tN(GUU)F | 1220 |
| duplication | VI:140721-145400 | YFL002W-A (transposon) | 4680 |
| duplication | VI:188481-197060 | YFR019W, tS(GCU)F, YFR020W, YFR012W (known duplicate) | 8580 |
| duplication | VI:199821-201580 | YFR023W (known duplicate) | 1760 |
| duplication | VI:232061-240040 | YFR039C (known duplicate), YFR040W (known duplicate), YFR041C, YFR042W, YFR043C | 7980 |
| duplication | VI:242341-270140 | telomere | 27800 |
| duplication | VII:1-65380 | telomere | 65380 |
| duplication | VII:69501-71000 | YGL227W | 1500 |
| duplication | VII:110381-113880 | tH(GUG)G1, YGL204C, YGL203C | 3500 |
| duplication | VII:147381-148880 | YGL189C (known duplication) | 1500 |
| deletion | VII:383781-385360 | YGL063W (known duplication), YGL063C-A, YGL062W (known duplication) | 1580 |
| duplication | VII:496241-498020 | YGL001C, YGR001C | 1780 |
| duplication | VII:534901-541700 | YGR027W-A (transposon) | 6800 |
| duplication | VII:562341-569040 | YGR038C-B (transposon) | 6700 |
| duplication | VII:618041-623060 | YGR065C, YGR066C, YGR067C | 5020 |
| duplication | VII:711601-713960 | YGR109W-B (transposon), YGR110W | 2360 |
| duplication | VII:818021-823500 | YGR161C-D (transposon) | 5480 |
| duplication | VII:909761-912260 | YGR205W, YGR206W, YGR207C, YGR208W | 2500 |
| duplication | VII:922541-923860 | YGR216C | 1320 |
| duplication | VII:930881-932000 | tG(GCC)G2 | 1120 |
| duplication | VII:956381-958260 | YGR233C | 1880 |
| duplication | VII:1007441-1009500 | YGR258C | 2060 |
| duplication | VII:1037641-1090960 | telomere | 53320 |
| duplication | VIII:1-48700 | telomere | 48700 |
| duplication | VIII:71021-72800 | YHL017W (known duplication), YHL016C | 1780 |
| duplication | VIII:74481-92460 | YHL009W-B (transposon) | 17980 |

| 1693 | | | |
|---|---|---|---|
| CNV type | coordinates | context/ORF | CNV_size |
| duplication | I:1-25540 | telomere | 25540 |
| duplication | I:26901-31460 | YAL063C (FLO9) | 4560 |
| duplication | I:38541-40700 | YAL058W, YAL056W (known duplication) | 2160 |
| deletion | I:65781-69840 | YAL040C (CLN3) | 4060 |
| duplication | I:159981-162760 | YAR009C (transposon) | 2780 |
| duplication | I:165241-166480 | YAR010C (transposon), tA(UGC)A | 1240 |
| duplication | I:175041-189220 | YAR020C, YAR023C, tL(CAA)A, tS(AGA)A, YAR027W, YAR028W, | 14180 |
| duplication | I:194561-203780 | telomere | 9220 |
| deletion | I:204581-205340 | telomere | 760 |
| duplication | I:205781-211960 | telomere | 6180 |
| deletion | I:211961-212200 | telomere | 240 |
| deletion | I:213941-214180 | telomere | 240 |
| duplication | I:214181-220000 | telomere | 5820 |
| duplication | I:223341-230220 | telomere | 6880 |
| duplication | II:1-30200 | telomere | 30200 |
| deletion | II:30201-31780 | YBL100W-B (transposon) | 1580 |
| duplication | II:31781-36300 | YBL100W-B (transposon) | 4520 |
| deletion | II:469701-474240 | YBR115C (LYS2) | 4540 |
| duplication | III:1-11880 | telomere | 11880 |
| duplication | III:14041-46920 | YCL064C, YCL061C, YCL059C, YCL057W, YCL055W, | 32880 |
| deletion | III:84781-88340 | YCL019W (LEU2) | 3560 |
| deletion | III:90521-92480 | YCL019W (LEU2) | 1960 |
| deletion | III:148601-151720 | YCR018C, tM(CAU)C, tK(CUU)C | 3120 |
| duplication | III:293421-314100 | telomere | 20680 |
| duplication | IV:1-15560 | telomere | 15560 |
| deletion | IV:302941-304400 | YDL085W (known duplication) | 1460 |
| deletion | IV:352881-355380 | YDL056W (MBP1) | 2500 |
| duplication | IV:528561-531500 | YDR038C (ENA5) | 2940 |
| deletion | IV:531501-531760 | YDR039C (ENA2) | 260 |
| duplication | IV:532581-537120 | YDR40C (ENA1) | 4540 |
| duplication | IV:645381-652160 | YDR098C-B,A (transposn) | 6780 |
| duplication | IV:987561-992660 | YDR261C-D,C (transposon) | 5100 |
| duplication | IV:1096641-1101680 | YDR261C-D,C (transposon) | 5040 |
| duplication | IV:1207321-1212260 | YDR365W-B (transposon) | 4940 |
| deletion | IV:1333441-1334700 | YDR435C | 1260 |
| duplication | IV:1525881-1531940 | telomere | 6060 |
| duplication | IX:1-2280 | telomere | 2280 |
| duplication | IX:7261-18440 | telomere | 11180 |
| duplication | IX:196041-197840 | telomere | 1800 |
| duplication | IX:383481-439900 | telomere | 56420 |
| duplication | Mito:1-2960 | mitochondrial | 2960 |
| duplication | Mito:3021-34060 | mitochondrial | 31040 |
| duplication | Mito:34161-52380 | mitochondrial | 18220 |
| duplication | Mito:52501-70560 | mitochondrial | 18060 |
| duplication | Mito:70661-72900 | mitochondrial | 2240 |
| duplication | Mito:73201-77100 | mitochondrial | 3900 |
| duplication | Mito:77221-81480 | mitochondrial | 4260 |
| duplication | Mito:81521-85780 | mitochondrial | 4260 |
| duplication | V:1-39080 | telomere | 39080 |
| deletion | V:115921-117060 | YEL021W (URA3) | 1140 |
| duplication | V:443621-448260 | YER138C (transposon) | 4640 |
| deletion | V:472361-475940 | YER152C, YER152W-A, YER153C, YER154W | 3580 |
| duplication | V:492601-497520 | YER160C (transposon) | 4920 |
| duplication | V:561701-576880 | telomere | 15180 |
| duplication | VI:4981-12860 | YFL061W (DDI2), YFL060C (SNO3) | 7880 |
| duplication | VI:143381-146340 | YFL002W-A (transposon) | 2960 |
| duplication | VII:1-52960 | | 52960 |
| deletion | VII:270801-272640 | YGL126W, YGL125W | 1840 |
| duplication | VII:535521-538220 | YGR027W-A (transposon) | 2700 |
| duplication | VII:539621-541540 | YGR027W-A (transposon) | 1920 |
| duplication | VII:562101-568880 | YGR038C-B (trasnposon) | 6780 |
| duplication | VII:817521-823940 | YGR161C-D (transposon) | 6420 |
| duplication | VII:1046041-1047900 | YGR277C, YGR278W | 1860 |
| duplication | VII:1080841-1090960 | telomere | 10120 |
| duplication | VIII:1-23520 | telomere | 23520 |
| duplication | VIII:212381-216140 | RUF5-1, YHR054C (known partial duplication), RUF5-2, YHR056C (known duplication) | 3760 |
| deletion | VIII:390601-391600 | YHR146W (known duplication) | 1000 |
| duplication | VIII:526301-537700 | telomere | 11400 |
| duplication | VIII:543621-545680 | telomere | 2060 |
| duplication | VIII:547241-557160 | telomere | 9920 |
| deletion | VIII:557721-557960 | telomere | 240 |
| duplication | VIII:557961-562620 | telomere | 4660 |
| duplication | X:1-19220 | telomere | 19220 |
| duplication | X:72701-74820 | YJL192C, YJL191W (ribosomal protein) | 2120 |
| deletion | X:360221-362040 | YJL043W, YJL042W | 1820 |
| deletion | X:362081-363520 | YJL042W | 1440 |
| duplication | X:471541-483840 | YJR026W, YJR027W, YJR029W (transposons) | 12300 |
| deletion | X:637941-640340 | upstream of YJR115W (known duplication) | 2400 |
| duplication | X:715481-731580 | telomere | 16100 |
| deletion | X:731621-731900 | telomere | 280 |
| deletion | X:732801-734900 | telomere | 2100 |
| duplication | X:735141-737200 | telomere | 2060 |
| duplication | X:737241-745760 | telomere | 8520 |
| duplication | XI:1-12840 | telomere | 12840 |
| deletion | XI:66581-69500 | YKL201C (known duplication), YKL198C (known duplication) | 2920 |
| deletion | XI:170201-173780 | YKL148C, YKL147C, YKL146W | 3580 |
| deletion | XI:525481-527260 | YKR047W, YKR048C, YKR049C | 1780 |
| duplication | XI:635721-666820 | telomere | 31100 |
| duplication | XII:5541-20060 | telomere | 14520 |
| deletion | XII:216321-217240 | YLR035C-A (transposon) | 920 |
| deletion | XII:329801-330960 | YLR095C | 1160 |
| duplication | XII:451381-468840 | RDN25-1, RDN37-1, YLR154W-F, RND5-1 RDN37-2, RDN18-2 | 17460 |
| duplication | XII:476221-478220 | YLR157C-B (transposon) | 2000 |
| deletion | XII:479341-479900 | YLR157C-B (transposon) | 560 |
| deletion | XII:481721-482140 | YLR157C-A (transposon) | 420 |
| duplication | XII:490041-490320 | upstream of YLR162W-A | 280 |
| duplication | XII:593781-598680 | YLR227W-B (transposon) | 4900 |
| duplication | XII:651481-652880 | YLR256W-A (transposon) | 1400 |

| | | | |
|---|---|---|---|
| duplication | VIII:112881-120180 | YHR005C, YHR005C-A, YHR006W (known duplication) | 7300 |
| duplication | VIII:212381-216140 | RUF5, YHR054C | 3760 |
| duplication | VIII:472581-475180 | YHR184W, YHR185C | 2600 |
| duplication | VIII:525161-548600 | telomere | 23440 |
| deletion | VIII:548601-548840 | telomere | 240 |
| duplication | VIII:548841-557720 | telomere | 8880 |
| deletion | VIII:557721-557960 | telomere | 240 |
| duplication | VIII:557961-562660 | telomere | 4700 |
| duplication | X:1-20100 | telomere | 20100 |
| duplication | X:28221-35320 | YJL214W, YJL213W, YJL212C | 7100 |
| duplication | X:38601-48180 | YJL209W, YJL208C, YJL207C, YJL206C | 9580 |
| deletion | X:120421-121800 | YJL159W (known duplication) | 1380 |
| duplication | X:472521-483960 | YJR026W, YJR027W, YJR029W (transposons) | 11440 |
| deletion | X:638901-640260 | YJR113C (ribosomal protein), YJR115W (known duplication) | 1360 |
| duplication | X:721361-731460 | telomere | 10100 |
| deletion | X:731461-734220 | telomere | 2760 |
| deletion | X:734641-734900 | telomere | 260 |
| duplication | X:734901-745760 | telomere | 10860 |
| duplication | XI:1-51140 | telomere | 51140 |
| duplication | XI:361501-363400 | YKL040C, YKL039W (known duplication) | 1900 |
| duplication | XI:373521-378920 | YKL034W, YKL033W-A, YKL033W | 5400 |
| duplication | XI:430241-431780 | YKL006C, snR87 | 1540 |
| deletion | XI:527861-529340 | YKR050W (known duplication) | 1480 |
| duplication | XI:602601-606000 | YKR086W, YKR087C, YKR088C, YKR089C (known duplication) | 3400 |
| duplication | XI:619541-666820 | telomere | 47280 |
| deletion | XII:441-700 | telomere | 260 |
| duplication | XII:701-20140 | telomere | 19440 |
| duplication | XII:20181-59380 | telomere | 39200 |
| duplication | XII:208741-216400 | YLR035C-A (transposon) | 7660 |
| deletion | XII:216401-217720 | YLR035C-A (transposon) | 1320 |
| duplication | XII:217721-244020 | YLR035C-A (transposon), YLR036C, YLR038C, YLR039C, YLR040C, YLR043C (known duplication), YLR045C, YLR046C (known duplication), YLR047C, YR048W (known duplication) | 26300 |
| duplication | XII:412301-413760 | YLR134W, YLR135W | 1460 |
| duplication | XII:436201-437460 | YLR148W | 1260 |
| duplication | XII:451461-469960 | RDN25-1, RDN37-1, YLR154W-F, RDN5-1, RDN37-2, RDN18-2, RDN5-2 | 18500 |
| deletion | XII:474301-474580 | intergenic, adjacent to transposon | 280 |
| deletion | XII:475841-476060 | YLR157W-D (known duplication, adjacent to transposon) | 220 |
| duplication | XII:476081-478180 | YLR157C-B (transposon) | 2100 |
| duplication | XII:480081-481620 | YLR157C-A (transposon) | 1540 |
| duplication | XII:483721-485860 | YLR159W (known duplication), RDN5-5 | 2140 |
| duplication | XII:487141-489140 | YLR160C (known duplication, near rNDA), YLR161W (known duplication, near rDNA) | 2000 |
| duplication | XII:490041-490320 | intergenic near unknown proteins | 280 |
| duplication | XII:593861-599740 | YLR227W-B (transposon) | 5880 |
| duplication | XII:650901-657860 | YLR256W-A (transposon) | 6960 |
| deletion | XII:732281-734700 | YLR303W (MET15) | 2420 |
| duplication | XII:748701-751900 | YLR309C | 3200 |
| duplication | XII:792421-794180 | tD(GUC)L2 | 1760 |
| duplication | XII:794221-795640 | snR61, snR55, snR57, YLR333C (ribosomal) | 1420 |
| duplication | XII:944341-947220 | YLR410W-B (transposon) | 2880 |
| duplication | XII:976761-981600 | | 4840 |
| duplication | XII:1028761-1034320 | YLR448W (ribosomal), YLR449W, YLR450W | 5560 |
| duplication | XII:1036941-1078180 | telomere | 41240 |
| duplication | XIII:1-32760 | telomere | 32760 |
| duplication | XIII:184261-190060 | YML045W (transposon) | 5800 |
| duplication | XIII:197501-198380 | YML039W (transposon) | 880 |
| duplication | XIII:198621-202340 | YML039W (transposon) | 3720 |
| duplication | XIII:356301-363100 | YMR045C (transposon) | 6800 |
| duplication | XIII:372681-378700 | YMR050C (transposon) | 6020 |
| deletion | XIII:388021-389960 | YMR056C, YMR057C, YMR058W | 1940 |
| duplication | XIII:502741-505060 | tL(CAA)M | 2320 |
| duplication | XIII:648641-650060 | YMR129W | 1420 |
| duplication | XIII:841081-857240 | YMR287C, YMR288W (snRNP), YMR289W, YMR291W, YMR292W, YMR293C | 16160 |
| duplication | XIII:863401-924440 | telomere | 61040 |
| duplication | XIV:6621-43160 | telomere | 36540 |
| duplication | XIV:96621-100700 | YNL284C-B | 4080 |
| deletion | XIV:196481-200380 | YNL241C, YNL240C | 3900 |
| duplication | XIV:520941-525120 | YNL054W-B (transposon) | 4180 |
| duplication | XIV:561181-563820 | | 2640 |
| duplication | XIV:672441-674320 | YNR024W, YNR025C, YNR026C (known duplication) | 1880 |
| duplication | XIV:715921-754020 | YNR052C, snR191, tL(UAA)N, YNR055C, YNR056C, YNR058W, YNR059W, YNR060W, YNR062C, YNR063W, YNR065C | 38100 |
| duplication | XIV:754061-784340 | telomere | 30280 |
| duplication | XV:1-53600 | telomere | 53600 |
| duplication | XV:97381-99040 | YOL117W | 1660 |
| duplication | XV:110821-113620 | tG(UCC)O, YOL108C, YOL107W, YOL106W | 2800 |
| duplication | XV:117841-123720 | YOL103W-B (transposon) | 5880 |
| duplication | XV:179641-181920 | YOL081W (known duplication), YOL080C, YOL079W, YOL078W | 2280 |
| duplication | XV:226641-228760 | tG(GCC)O1, tN(GUU)O1, YOL054W | 2120 |
| duplication | XV:299281-300620 | YOL014W | 1340 |
| duplication | XV:436221-439080 | YOR058C, tK(UUU)O, YOR059C | 2860 |
| deletion | XV:479621-481280 | YOR083W (WHI5) | 1660 |
| duplication | XV:594301-600620 | YDR142W-B (transposon) | 6320 |
| deletion | XV:637081-639680 | YOR161C, YOR161W-B, YOR161C-C, YOR162C (known duplication) | 2600 |
| deletion | XV:676061-677940 | YOR181W | 1880 |
| duplication | XV:703441-709040 | YOR192C-B (transposon) | 5600 |
| deletion | XV:722241-722460 | YOR202W (HIS3) | 220 |
| duplication | XV:972901-976300 | YOR343W-B (transposon) | 3400 |
| duplication | XV:1065781-1091280 | telomere | 25500 |
| duplication | XVI:1-62880 | telomere | 62880 |
| duplication | XVI:435841-440720 | YPL060C-A (transposon) | 4880 |
| duplication | XVI:804641-810700 | YPR137C-B (transposon) | 6060 |
| duplication | XVI:844301-855920 | YPR158W-A,B, YPR158C-C,D (transposons) | 11620 |
| duplication | XVI:881041-883680 | YPR170C, YPR169W-A | 2640 |
| duplication | XVI:923401-947800 | telomere | 24400 |

| | | | |
|---|---|---|---|
| duplication | XII:654281-656760 | YLR256W-A (transposon) | 2480 |
| duplication | XII:945781-946980 | YLR410W-B (transposon) | 1200 |
| duplication | XII:1069281-1078180 | telomere | 8900 |
| duplication | XIII:1-14980 | telomere | 14980 |
| deletion | XIII:68021-73040 | YML102W, YML101C-A, YML101C (known duplication), YML100W (known dupliction) | 5020 |
| duplication | XIII:184041-190120 | YML045W (transposon) | 6080 |
| duplication | XIII:199181-202340 | YML039W (transposon) | 3160 |
| duplication | XIII:372781-377660 | YMR050C (transposon) | 4880 |
| deletion | XIII:489161-490320 | YMR109W (known duplication) | 1160 |
| deletion | XIII:817581-820100 | YMR275C (known duplication), YMR276W | 2520 |
| duplication | XIII:908121-924400 | telomere | 16280 |
| duplication | XIV:1-11520 | telomere | 11520 |
| duplication | XIV:14741-18160 | YNL332W, YNL331C, YNL330C | 3420 |
| duplication | XIV:96781-100660 | YNL284C-B (transposon) | 3880 |
| duplication | XIV:561361-563400 | between tT(AGU)N2 and tP(UGG)N2 | 2040 |
| duplication | XIV:725481-784340 | telomere | 58860 |
| duplication | XV:1-29400 | telomere | 29400 |
| duplication | XV:119001-123400 | YOL103W-B (transposon) | 4400 |
| duplication | XV:226141-228920 | tG(GCC)O1, tN(GUU)O1, YOL054W | 2780 |
| deletion | XV:480041-482540 | YOR083W, YOR084W, YOR085W | 2500 |
| deletion | XV:587881-589800 | YOR140W | 1920 |
| duplication | XV:598561-600640 | YDR142W-B (transposon) | 2080 |
| deletion | XV:676221-677980 | YOR181W (LAS17) | 1760 |
| duplication | XV:708001-709100 | YOR181W (LAS17) | 1100 |
| deletion | XV:921801-922620 | YOR323C | 820 |
| duplication | XV:974801-978240 | YOR343W-B (transposon) | 3440 |
| duplication | XV:1084821-1091300 | telomere | 6480 |
| duplication | XVI:1-12480 | telomere | 12480 |
| duplication | XVI:59921-63140 | YPL257W-B (transposon) | 3220 |
| duplication | XVI:805661-808860 | YPR137C-B (transposon) | 3200 |
| duplication | XVI:847841-855960 | YPR158W-A,B, YPR158C-C,D (transposons) | 8120 |

**Supplemetary Table 13: CNVnator Analysis, bin size = 100.** Events that are located in repetative regions such as telomeres, transposons, and ribosomal RNA genes are shaded light grey. Events which could be visually confirmed by alighning the reads t

| 1691 | | | |
|---|---|---|---|
| CNV type | coordinates | ORF(s) impacted | CNV_size |
| duplication | I:1-27500 | telomere | 27500 |
| duplication | I:192001-230500 | telomere | 38500 |
| duplication | II:1-30500 | telomere | 30500 |
| deletion | II:470001-474000 | YBR115C (LYS2) | 4000 |
| duplication | II:760001-813500 | telomere | 53500 |
| duplication | III:1-32500 | telomere | 32500 |
| deletion | III:85001-89000 | YCL018W (LEU2) | 4000 |
| deletion | III:91001-92500 | YCL018W (LEU2) | 1500 |
| deletion | III:148501-151500 | YCR018C, tM(CAU)C, tK(CUU)C | 3000 |
| duplication | III:293501-317000 | telomere | 23500 |
| duplication | IV:1-14500 | telomere | 14500 |
| deletion | IV:353001-355500 | YDL056W (MBP1) | 2500 |
| duplication | IV:874001-885000 | YDR210W-B, YDR210C-D, YDR210C-C (transposons) | 11000 |
| duplication | IV:1524001-1531500 | telomere | 7500 |
| duplication | IX:405501-440000 | telomere | 34500 |
| duplication | VI:1-13000 | telomere | 13000 |
| duplication | VII:1-28000 | telomere | 28000 |
| duplication | VIII:1-24000 | telomere | 24000 |
| duplication | VIII:529001-563000 | telomere | 34000 |
| duplication | X:1-20000 | telomere | 20000 |
| duplication | X:472001-484000 | YJR026W,YJR027W, YJR029W (transposons) | 12000 |
| duplication | X:704501-746000 | telomere | 41500 |
| duplication | XII:1-54500 | telomere | 54500 |
| duplication | XII:451501-471500 | RDN25-1, RDN37-1, RDN18-1, RDN5-1, RDN37-2, RDN18-2, RDN5-2 | 20000 |
| deletion | XII:732501-734500 | YLR303W (MET17) | 2000 |
| duplication | XII:1056501-1078000 | telomere | 21500 |
| duplication | XIV:754001-784500 | telomere | 30500 |
| duplication | XV:1-29500 | telomere | 29500 |
| duplication | XVI:6001-63000 | telomere | 57000 |
| duplication | XVI:846001-857000 | YPR158W-B, YPR158C-D, YPR158C-C (transposons) | 11000 |

| 1693 | | | |
|---|---|---|---|
| CNV type | coordinates | ORF(s) impacted | CNV_size |
| duplication | I:1-66000 | telomere | 66000 |
| deletion | I:66001-67500 | YAL040C(CLN3) | 1500 |
| duplication | I:194501-230500 | telomere | 36000 |
| duplication | II:1-30000 | telomere | 30000 |
| deletion | II:470001-474000 | YBR115C (LYS2) | 4000 |
| duplication | III:1-29000 | telomere | 29000 |
| deletion | III:85001-88500 | YCL018W (LEU2) | 3500 |
| deletion | III:90501-92500 | YCL018W (LEU2) | 2000 |
| deletion | III:148501-151500 | YCR018C, tM(CAU)C, tK(CUU)C | 3000 |
| duplication | III:293001-317000 | telomere | 24000 |
| duplication | IV:1-14500 | telomere | 14500 |
| deletion | IV:353001-355500 | YDL056W (MBP1) | 2500 |
| deletion | V:116001-117000 | YEL021W (URA3) | 1000 |
| duplication | VI:1-13000 | telomere | 13000 |
| duplication | VII:1-32500 | telomere | 32500 |
| duplication | VIII:516001-563000 | telomere | 47000 |
| duplication | X:471501-484000 | YJR026W,YJR027W, YJR029W (transposons) | 12500 |
| duplication | X:735001-746000 | telomere | 11000 |
| duplication | XII:1-50500 | telomere | 50500 |
| duplication | XII:451501-473500 | RDN25-1, RDN37-1, RDN18-1, RDN5-1, RDN37-2, RDN18-2, RDN5-2, YLR156W | 22000 |
| duplication | XIV:746001-784500 | telomere | 38500 |
| duplication | XV:1-29500 | telomere | 29500 |
| duplication | XVI:1-65500 | telomere | 65500 |
